# Seed-applied multi-kingdom synthetic communities selectively reshape bacterial communities and highlight key criteria for strain selection

**DOI:** 10.64898/2026.07.15.738658

**Authors:** Logan Suteau, Claire Campion, Coralie Marais, Martial Briand, Anaïs Hardouin, Kaat Hellyn, Kenji Maurice, Muriel Marchi, Marie Simonin, Natalia Guschinkaya

## Abstract

Defined microbial communities (also known as synthetic communities) are showing promising results for plant health but are lacking an efficient lab to field transition. This is the due to limited knowledge on how to efficiently modulate plant microbiota by considering complex environments and multi-kingdom interactions. In this study, we aimed to better understand the transmission and impact of multi-kingdom synthetic communities (SynComs) from the seed to seedling stage. We constructed 20 different SynComs using both *a priori* and random approaches, from pool of diverse strains including 24 bacteria, 11 yeasts and 10 filamentous fungi. SynComs were inoculated on *Brassica napus* seeds and we monitored both transmission and impact on the microbiota of 15-day-old seedlings grown in non-sterile soil. Optimization of the inoculation protocol showed that alginate coating improved bacterial, yeast and filamentous fungi concentrations by more than 2 log compared with other approaches. With this inoculation method, we observed contrasted seedling colonization profiles, with SynComs members representing between 1.1-45.7% of bacterial community and 3.2-36.6% of fungal community. Our multiple SynCom design revealed that strain selection is a more critical determinant of SynCom performance than assembly strategy. Even randomly assembled communities performed well, as long as they are drawn from a pool of ecologically relevant, well-adapted taxa. Based on these evidences, we identified key bacterial and fungal traits explaining efficient seedling colonization such as high abundance on inoculated seed and low *in vitro* lag-time. Despite low colonization levels, we observed that SynCom inoculation altered seeding bacterial community assembly in 14 SynComs. A total of 82 native bacterial ASVs were identified as responsive to SynCom inoculation, most likely originating from the soil. This shift indicates that SynComs influence community assembly by modulating the recruitment of environmental taxa, especially when SynCom strains were more integrated in multi-kingdom network structures. Finally, we identified four distinct SynComs profiles which either colonized strongly or not seedlings while shifting of not native microbiota. Altogether, these findings provide actionable directions for improving SynCom design, suggesting that leveraging ecological processes such as host adaptation, optimal inoculation density, and network integration could enhance both colonization efficiency and plant phenotypic outcomes.

## Background

An increasing number of studies use defined microbial communities [1] also known as synthetic communities (SynComs) [2] on plants to modulate plant phenotype [3] and protect against biotic and abiotic stressors [4, 5]. While having beneficial effects in controlled conditions, these methods are not as performant in the field and suffer a lack of efficient transfer from lab to more realistic conditions [6, 7]. With most studies mainly focusing on phenotypic effect and neglecting colonization dynamics of the SynCom, low performance can be often attributed to a weak establishment of the inoculant [7]. Furthermore, the lasting effects of these inoculants on the native microbiota is still underestimated despite the ecological risks they imply [8]. These knowledge gaps highlight the needs for a better monitoring of the colonization process of microbial inoculants on plants.

Microbial interactions play an important role in the assembly and dynamics of plant and soil microbiota. When inoculated, SynCom encounters native communities (e.g. soil, seed), triggering community coalescence described as the joining of previously separate communities into a new entity [9]. During this event, both inoculated and native strains will compete for similar resources [10, 11] and engage in molecular dialogue [12]. Pioneer studies on SynCom inoculation suggest that both the number of organisms (i.e. propagule size [13]) and the timing of arrival (i.e. priority effects [14]) are key drivers of the successful establishment within communities [15]. Thus, a better understanding and consideration of the ecological processes at play during SynCom inoculation is key to improve inoculation success [16].

One way to harness these ecological processes is also to target specific “critical windows” during plant development that could be more favorable for inoculant establishment. For instance, at the seed stage native plant microbiota is sparse, favoring inoculant establishment through priority effect and propagule pressure [14]. Improvement in seed technologies also allows the coating of seed for a more efficient delivery of microorganisms [17]. The inoculation and monitoring of bacterial communities in seed to seedling stages has shown promising results with high colonization [15, 18]. However, like most studies using SynComs inoculation experiments, the focus has primarily been on bacterial communities, while only a few studies consider both bacterial and fungal inoculation (4 out of 86 surveyed in Xu et al., 2025 [19]). Yet, interkingdom interactions between bacteria and fungi naturally occur within plant microbiota and modulate plant phenotype [20]. More recently, an increasing interest has grown in multi-kingdom inoculum as they enhance disease suppressiveness [21, 22]. Still, while we know that complex interactions occur between these kingdoms [23], limited knowledge on how to design and inoculate efficiently these multi-kingdom inoculants is available. An upgraded guideline for the use of multi-kingdom inoculant is then required to fully harness the beneficial effects mediated by these multi-kingdom interactions.

In this study, we aim to gain a deeper understanding of multi-kingdom SynCom inoculation to characterize their colonization success of seeds and seedling, and their impact on the native microbiome. To do so, 20 different SynComs were designed based on distinct criteria and inoculated onto *Brassica napus* L. seeds and grown for 15 days in non-sterile soil. We hypothesized that both fungal and bacterial strains inoculated onto seed would be transmitted to the seedling (H1).

As SynCom outperforms single strain inoculant in colonization and survival [24], a better understanding of efficient strain combination is required. That is why we hypothesized that SynCom designs based on specific assembly criteria would achieve higher colonization than those assembled randomly (H2). We aimed to find easily characterizable traits for an entire microbial collection such as natural abundance and prevalence in target habitat (here seed and seedling) as potential indicators of host adapation [25], taxonomic distance as indicator of reduced niche competition [26] and growth speed for quicker access to resources [27].

Even though SynCom composition should be a main driver of colonization through microbial interactions, specific strain traits might be linked to a higher colonization. Thus, identifying these traits could allow a future better selection of strains for SynCom composition. Here, we also hypothesized that strains traits could influence their colonization success but could differ between bacteria and fungi (H3). We investigated traits relative to the native abundance and prevalence of strains, their proportion in inoculant (relative abundance in inoculated seeds) as well as basic phenotyping traits (*in vitro* growth parameters).

Finally, plant microbiome assembly is a dynamic process that evolves throughout development [28]. In native conditions, seedling microbiota is mostly shaped by its environment [29, 30]. In this context, ecological processes such as priority effects [14] and community coalescence [9] are known to play key roles in shaping microbiome structure and dynamics. We therefore hypothesized that SynCom inoculation would alter the recruitment of both soilborne and native seedborne microbiota during seedling microbiota assembly, and that these shifts would be associated with specific inoculated strains or SynCom composition (H4)

## Methods

### Biological material

#### Microbial strains

The bacterial and fungal strains used in this study were obtained from Simonin *et al*., 2026 that aimed to create a diverse collection of seed-borne microorganisms [31]. Briefly, seeds from four *Brassica napus* L. (oilseed rape) genotypes (Jupiter, Aviso, Mohican, Milena), produced under two production modes (open-field vs self-pollinated cages), were used for isolation from both seeds and seedlings germinated under gnotobiotic conditions. The bacterial strains were isolated on Tryptic Soy Agar (TSA) 10% strength and fungal strains on Malt Agar (MA) medium. This collection resulted in a total of 636 bacterial and 188 fungal strains. A preliminary genotyping of all isolates was performed using metabarcoding of the *gyrB* bacterial marker gene (Illumina Miseq) or NS7/ITS4 + D1/D2 regions for yeasts (Sanger sequencing) and NS7-ITS4 for filamentous fungi (Sanger sequencing). The representativeness of the strain collection was assessed by comparing it to the bacterial and fungal community composition of the same seeds and seedlings used for strain isolation. It showed that 17% of the bacterial diversity and 6% of the fungal diversity detected by metabarcoding in seed lots was recovered through isolation. Even if these numbers appear to be low, the most prevalent and abundant taxa were isolated and the collection represented 69% (bacteria) and 75% (fungi) of the total relative abundance of the seed microbiota.

#### Plant material

We used winter oilseed rape seeds of the Jupiter genotype produced under open-field conditions in 2020 at the IGEPP field station, Le Rheu, Bretagne (48.13916, −1.799776) and stored at 9°C. These seeds are from the same seed lot than the one used for the microbial collection.

#### Strain culture conditions and phenotyping

Bacterial strains were routinely grown on 10% TSA medium at 18°C and fungal strains were grown on MA medium at 20°C. For growth rate assessment, bacterial strains were adjusted to an optical density of 600 nm (OD600) of 0.01 in 10% Tryptic Soy Broth (TSB) and monitored by spectrophotometry. Optical density was measured with a spectrophotometer over 72h at 25°C. Yeasts and filamentous fungi growth was also measured in liquid culture in Malt Broth (MB) at a final concentration of 10^5^ cells.mL^-1^ using nephelometry [32] at 25°C over 72h.

### SynCom strains selection

A subset of strains from the microbial collection was preselected for SynCom design. Further criteria have been used.

#### Taxonomy

First, one representative strain per genus was selected to capture the phylogenetic diversity associated with oilseed rape seeds. For the highly abundant genera (*Pantoea*, Pseu*domonas, Vishniacozyma, Holtermaniella* and *Cladosporium*), multiple strains were selected. This resulted in a preliminary set of 23 filamentous fungi, 27 yeasts and 60 bacterial strains chosen.

#### Strain growth rate *in vitro* condition

Each strain was cultured to verify purity and viability after conservation. Growth parameters were then assessed for all strains using liquid microplate cultures. When multiple strains shared the same unique Amplicon Sequence Variant (ASV), the strain with the highest growth rate was selected; when no differences were observed, one strain was randomly chosen.

#### Sporulation

For filamentous fungi, conidia production and spore viability were assessed, and strains unable to produce conidia were excluded as inoculation required conidia.

The final selection included 24 bacterial strains, 10 filamentous fungi and 11 yeasts with unique *gyrB* (bacteria) or ITS1 (fungi) ASV, enabling their detection in SynCom experiments *via* metabarcoding. Detailed information on all strains is provided in Table S1.

### SynCom design

A total of 20 different SynComs were built, 10 of them using *a priori* criteria and 10 using random strain sampling. 19 out of 20 SynCom consisted of 12 strains: 4 bacteria, 4 yeast, and 4 filamentous fungi, to approach a low microbial richness typically observed in seeds [33]. Using metabarcoding data from Simonin *et al.* (2026) [31], we assessed the natural prevalence and relative abundance of each strain in the seed and seedling microbiota. Based on these data, seven SynComs were constructed: one containing the most abundant strains on seeds, one with the most abundant strains on seedlings, one with the most abundant strains overall, and two containing rare strains (*i.e.,* strains isolated but not detected by metabarcoding). One of them was designed to represent an abundance gradient, including the most abundant strains on seed and seedling, two strains with intermediate abundance, and one rare strain. Finally, a high richness SynCom was assembled by combining all selected strains (45 strains in total). In addition, two SynComs maximizing phylogenetic diversity were constructed by selecting strains from distinct families. One SynCom was composed of the fastest-growing strains in synthetic media. The remaining 10 SynComs were designed randomly by sampling with replacement between independent SynCom, while maintaining the composition of 4 bacteria, 4 filamentous fungi and 4 yeasts. In total, 20 SynComs were designed (Fig. 1A). Across these SynComs, each bacterial strain was included between 2 and 7 times, each yeast strain between 5 and 10 times, and each filamentous fungi strain between 6 and 11 times. The representativeness of the assembled SynComs out of all possible strain combinations was calculated and represented in a theoretical composition space (Supplementary material and methods). To assess the difference in SynCom composition, their compositional dissimilarity was assessed according to supplementary materials and methods (Fig. 1B). In brief, pairwise compositional dissimilarity between all communities (theoretical and effective) was quantified using the binary Jaccard distance and distance matrices were subjected to Principal Coordinates Analysis (PCoA).

**Fig. 1:**
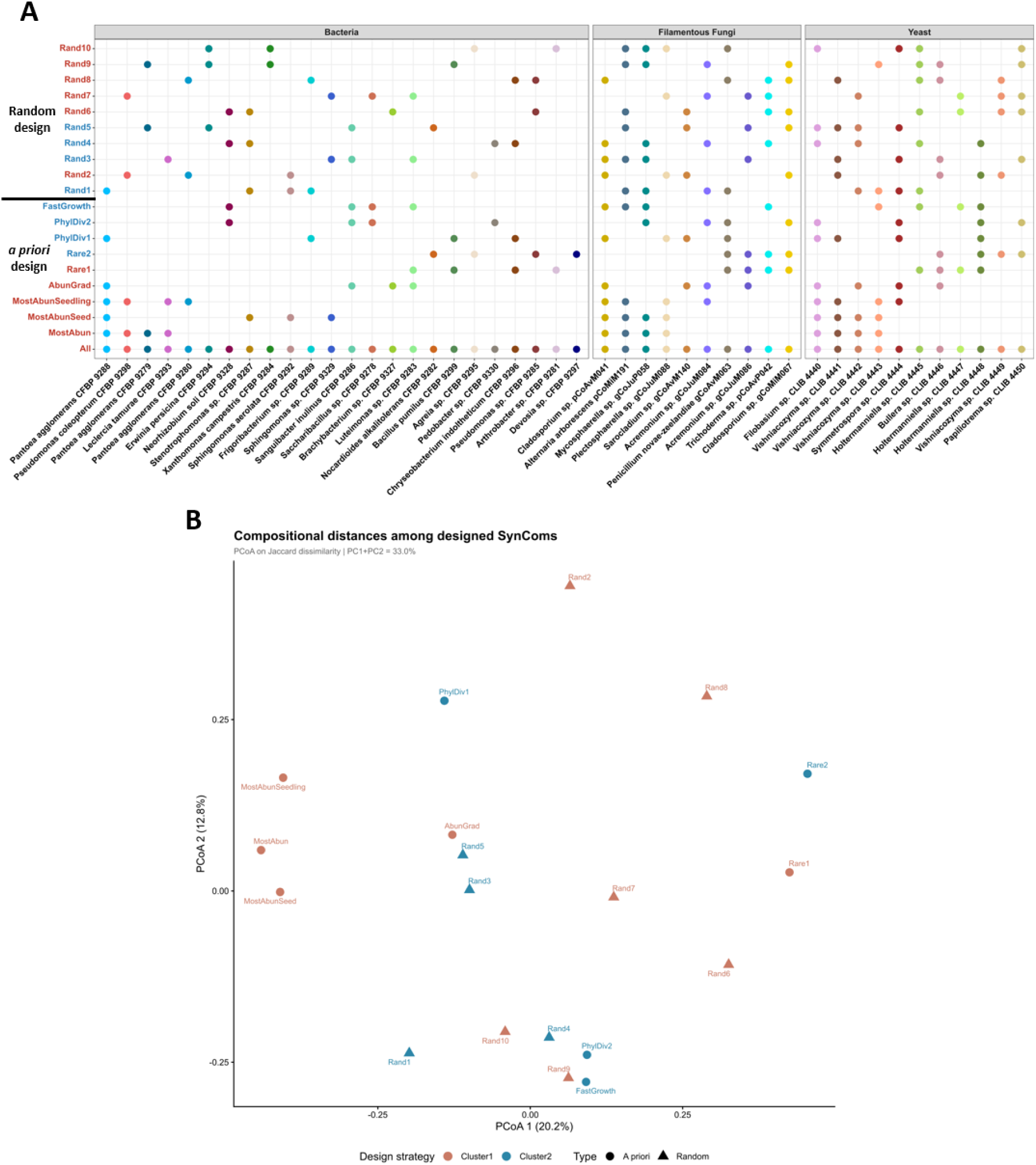
SynCom composition. (A) SynCom strain members and their distribution in the different SynComs used during this study. Strains are divided between Bacteria (n = 24), Filamentous fungi (n = 10) and Yeast (n = 11). SynComs are divided between Random and *a priori* constructions and colored to either Cluster 1 or Cluster 2 based on result presented later in the article. (B) Principal Coordinates Analysis (PCoA) of binary Jaccard distances among the 19 experimentally designed 4B+4F+4Y SynComs, based on a 45-strain binary presence–absence matrix (Cailliez correction). Points are shaped by design strategy and colored by cluster. Axis labels report the percentage of total variance explained. Individual SynCom identifiers are annotated.

### Seed inoculation optimization

To optimize the inoculation of SynComs on *Brassica napus* L. seeds, four inoculation methods were evaluated. The objectives were to increase microbial biomass on seeds, while maintaining a balanced proportion between bacteria, yeasts and filamentous fungi. In addition, we aimed to avoid negative effects on seed germination and to ensure reliable recovery of inoculated microorganisms for subsequent assessment of seed colonization. The four methods tested were seed soaking in different conditions (sterile water, 10mM MgCl_2_, 2.4% glycerol) or seed coating in 2.4% sodium alginate. Each condition was applied to 120 seeds.

Each method was used to inoculate a bacterial strain (*Pantoea agglomerans* CFBP 9288) a yeast (*Filobasium* sp. CLIB 4440), the fungal strain *Cladosporium* sp. pCoAvM041, and a control condition using sterile water instead of microbial suspension resulting in a total of four independent inoculations. Seeds were inoculated at a 10^6^ microorganisms.mL^-1^ concentration and dried for 10 min under a laminar flow hood. To assess seed colonization, eight seeds per each condition and per method were individually macerated in sterile water, or in 1% sodium citrate for alginate-coated seeds. Germination was evaluated by placing 100 seeds per condition on pleated paper, followed by incubation for 2 days in the dark at 20°C and 12 days under a 16h light and 8h dark photoperiod.

### SynCom inoculation on *Brassica napus* seed

The 20 SynComs were inoculated on *Brassica napus* seeds. Each strain was first prepared individually at a target concentration of 10^6^ microorganisms.mL^-1^, then mixed in equal proportions to assemble the SynComs. The resulting cultures were combined with sodium alginate to a final concentration of 2.4% method selected based on the previous experiment). Then, for each SynCom, 0.55 g of *Brassica napus* seeds (20 independent lots) were rinsed three times with sterile water and soaked in the SynCom suspensions for 30 min at 20°C under constant agitation. Seeds were then individually transferred into 10mM CaCl_2_ to induce alginate encapsulation, dried, and 84 seeds were sown in non-sterile potting soil (Traysubstrat 092, Klassmann) per condition. Plants were grown in a greenhouse for 15 days and watered twice prior to harvest. To assess seedling phenotypes based on seed testing standards, we followed the International Seed Testing Association ISTA [34] guidelines. We classified each plant in three potential categories: non germinated, normal seedling, abnormal seedling to estimate the emergence and normality rates. Seedlings were considered abnormal when more than 50% of the cotyledons or leaves were necrotic or rotten, if the hypocotyl or epicotyl were deformed, or if the root system was absent, stunted, or rotten. For each condition, eight entire normal seedlings were collected and whole seedlings microbial communities were analyzed using metabarcoding. Roots were washed in sterile water prior to DNA extraction to minimize the presence of soil-associated organisms. Seedling hypocotyl length was measured on the remaining seedlings to assess the potential impact of SynCom inoculation on plant phenotypes.

### DNA extraction, *gyrB* and ITS1 sequencing

The following metabarcoding approach was performed on the inocula, inoculated seeds and seedlings. For inocula, 200 μL of each fresh inoculum was instantly stored at −80°C prior to DNA extraction. For inoculated seeds, eight seeds per condition were individually macerated in 1% sodium citrate and incubated overnight (∼16h) at 4°C under agitation (150 rpm). A 200µL aliquot from each suspension was plated to assess seed inoculation and then stored at −80°C. Control samples consisted of a pool of 30 seeds macerated in 1mL under the same conditions, and processed identically. Whole seedling samples were crushed using a roller, followed by the addition of 5mL of sterile water. Samples were then ground for 30s using a stomacher. DNA was extracted using 200 μL of the resulting suspension using the NucleoSpin® 96 Food kit (Macherey-Nagel, Düren, Germany), following the manufacturer’s protocol. For potting soil, four replicates of 200 mg were sampled from control samples during seedling harvest and extracted using the DNeasy PowerSoil Pro kit (Qiagen), following the manufacturer’s protocol.

For bacterial communities, the *gyrB* gene was amplified using gyrB_aF64 (5’- MGNCCNGSNATGTAYATHGG-3’) and gyrB_aR353 (5’-ACNCCRTGNARDCCDCCNGA-3’) primers [35]. PCR reactions were performed with Q5 High-Fidelity DNA Polymerase (NEB, Ipswich, Massachusetts, USA) in a 50 μL reaction volume containing 10 μL of 5X Reaction Buffer, 10 µL of 5X High GC Enhancer, 1 µL of 10mM dNTP mix, 1 μL of each forward and reverse primers (100 μM), 0.5 μL of polymerase, and 10 μL of DNA. Cycling conditions included an initial denaturation step at 98◦C for 3 min, followed by 25 cycles of amplification at 98°C (30 s), 55°C (45 s), and 72°C (90 s), with a final elongation at 72°C for 10 min. Fungal communities identification was based on the internal transcribed spacer (ITS) region amplification using primers ITS1F_Mi (5’-CTTGGTCATTTAGAGGAAGTAA-3’) and ITS2_Mi (5’- GCTGCGTTCTTCATCGATGC-3’) [36]. Amplification was performed using the same conditions as described for *gyrB* except a primer concentration of 10µM and a hybridation step at 50°C.

Amplicons were purified using magnetic beads (Sera-MagTM, Merck, Kenilworth, New Jersey). A second PCR was performed to add Illumina adapters and barcodes, with the following cycling conditions: denaturation at 98°C (1 min), followed by 10 cycles at 98°C (1 min), 55°C (1 min), and 72°C (1 min), and a final elongation step at 72°C for 10 min. For ITS purification, an additional PCR product migration step on 1.8% agarose gel (70 V, 90 min) was performed to separate and recover fungal ITS (lower band, around 400 bp) from co-amplified plant ITS (higher band, >600 bp). The lower band was excised, extracted and purified using NucleoSpin Gel and PCR Clean up kit (Macherey-Nagel, Düren, Germany) and bead purified once more.

Library concentration was quantified using quantitative PCR (KAPA Library Quantification Kit, Roche, Basel, Switzerland). Libraries were mixed with 15% PhiX and sequenced using two MiSeq reagent kits v2 500 cycles (Illumina). Quality controls included a blank extraction kit control, a PCR-negative control, PCR-positive control (*Lactococcus piscium* DSM6634, a fish pathogen that is not plant-associated or *Saccharomyces cerevisiae*, a yeast that is not plant-associated) and a bacterial mock community, included in each PCR plate. Raw amplicon sequencing data are available at the European Nucleotide Archive (ENA) under accession number PRJEB91662.

### Long read genome extraction, sequencing, assembly and taxonomic affiliation

#### Genome extraction and sequencing

High molecular weight bacterial DNA was extracted using the Wizard Genomic DNA Purification kit (Promega, A1120) with a slightly modified protocol. Cells were grown for 24 h in 10% TSB and washed three times with 50 mM EDTA. To preserve DNA integrity, no vortexing was applied and pipetting was minimized. Protein precipitation was performed twice to improve DNA purity. High molecular weight fungal DNA was extracted using a modified Möller protocol [37]. In brief, cultures were flash frozen and crushed, then washed three times with methanol-0,1%2-mercaptoethanol. Cells were resuspended in TES (100mM TRIS pH 8.0, 10mM EDTA, 2% SDS) and 100µg of proteinase K at 60°C for 1h for cell lysis. Washing steps included 1.4M NaCl and 1% CTAB, protein precipitation with SEVAG (Chloroform:Isoamyl Alcohol (24:1 V:V)) on ice and 1.6M NH4Ac. DNA was precipitated with isopropanol and washed thrice with 70% ethanol before resuspension in water. DNA quality, concentration and fragment length were assessed using respectively a NanoDrop™ One Spectrophotometer, Qubit™ fluorometer and 1.5% gel electrophoresis. Afterwards, 1.5 µg of each strain’s DNA was used for long-read sequencing with the Ligation sequencing gDNA - Native Barcoding Kit 24 V14 (SQK-NBD114.24) on the MinION Mk1D (Oxford Nanopore Technologies) sequencer using a FLO-MIN114 flow cell. The raw electrical signals obtained were then converted into a DNA sequence using the Dorado software, applying version of the high accuracy basecalling model available at the time the sequencing was carried out.

#### Genome assembly and taxonomic affiliation

Bacterial genomes were assembled using Flye 2.9.5-b1801 [38]. Genomes that were not fully circularized were reassembled using different subsets of reads selected with filtlong. Checks for completeness and contamination were done with CheckM v1.2.2 [39] and taxonomic affiliation with gtdbtk v2.4.0 [40]. For fungal genomes, read filtering was done with NanoFilt 2.8.0, assembly with Flye 2.9.5-b1801, polishing with Racon v1.4.20 and biological completeness was estimated using BUSCO 6.0.0. Taxonomic affiliation was realized on the already known genus of the strain by using specific sequences detailed in Table S2. Each sequence used for comparison was obtained through *in silico* amplification on the assembled genome. For each specific gene, multiple sequence alignment was done using MAFFT (v7) to obtain a same length sequence of the gene for all compared species. All trimmed genes were then aggregated to create a phylogenetic tree (Supplementary Fig. 1 and Supplementary Fig. 2) with IQ-TREEusing Ultrafast bootstrap (1000 iterations). For each investigated genus, the set of strains used for comparison and their sequence accession are available in Table S3 to S11. All assembled genome accession numbers can be found in Table S1.

### Statistical and microbiota analyses

Bioinformatic processing of the amplicons was performed in R (v4.5.2). Primer sequences were removed using cutadapt v2.7 [41] and trimmed FASTQ files were processed with DADA2 v1.22.0 [39]. Reads were truncated at the first instance of a quality score ≤ 5, and chimeric sequences were identified and removed using the removeBimeraDenovo function. ITS1 sequences were processed using forward reads only, following Pauvert *et al.* (2019) [42], to maximize fungal diversity detection. ASVs were taxonomically assigned using a naïve Bayesian classifier with an in-house *gyrB* database (train_set_gyrB_v5.fa.gz, [43]) Unassigned sequences at the phylum level and *parE* sequences (a *gyrB* paralog) were removed. ITS1-derived ASVs were classified against the UNITE v.8 fungal database [44], and non-fungal ASVs were filtered out. All scripts are available at https://forge.inrae.fr/irhs-emersys/2026_logan_inoseed. Then, post-clustering curation was performed using LULU v0.1.0 [45] at a minimum threshold of 98% for *gyrB* and 99% for ITS. Contaminants were identified and removed using the decontam package v1.30.0, followed by manual curation. ASVs with fewer than 10 reads and detected in only one sample were removed from the dataset. Finally, samples with fewer than 1.000 reads after filtering were excluded from downstream analysis. To assess sequencing depth, rarefaction curves were generated using the rarecurve function of the vegan package (v2.7.2 [46]). Based on rarefaction curves, the *gyrB* dataset was rarefied at 3000 reads, and ITS dataset was rarefied at 2600 reads, resulting in 357 bacterial samples and 366 fungal samples. Rarefied ASV tables were then used to estimate species richness. For all other analyses, non-rarefied read counts were transformed into proportions (relative abundance) using the transform function from the microbiome R package. To estimate the effect of inoculation on species richness, observed richness was calculated on the rarefied dataset using the estimate_richness function of the phyloseq package (Version 1.54.0 ; [47]). To test the effect of SynCom inoculation on the microbiota, ordination analyses were performed using Bray-Curtis dissimilarity matrices computed on log(x+1) transformed non-rarefied data followed by Principal Coordinate Analysis (PCoA) using the ordinate function in phyloseq. To explore the percentage of variance explained by the factor SynCom on seed samples and seedling sample separately, permutational multivariate analysis of variance (PERMANOVA) was performed with 999 permutations using the adonis2 function from the vegan R package [46]. To further investigate the effect of SynCom on seedling microbiota, we used a clustering approach based on k-means clustering. The optimal number of clusters was determined using the NbClust package in R by evaluating 26 distinct diagnostic indices. To assess SynCom colonization, inoculated strains were traced back by matching the already known *gyrB* and ITS sequence of the strain used in the dataset. Then, the cumulative contribution of the SynCom strain was quantified by summing the relative abundances of the inoculated ASVs in each condition.

#### Generalized Additive Models

To elucidate the relationships between specific microbial traits and seedling colonization efficiency, we utilized Generalized Additive Models (GAMs) with the package mgcv [48] to explore both linear and non-linear traits correlations with colonization success, measured as the strain relative abundance in seedlings (response variable). A multivariate GAM was constructed using eight predictive variables: log10-transformed relative abundance on inoculated seeds, log10-transformed relative abundance in the initial inoculum, log10-transformed native relative abundance on seeds, native seed prevalence (defined as the proportion of occurrences across all sampled sites), genome size, growth rate, cumulative growth (Area Under the Curve, AUC), and lag phase duration. Relative abundance variables were log10-transformed to reduce the influence of highly abundant taxa. Independent multivariate models were fitted for bacterial and fungal datasets, with strain identity included as a random effect to account for phylogenetic structure. The relative importance of predictors within each model was quantified *via* hierarchical partitioning using the gam.hp package in R [49]. Model predictions were visualized using the ggplot2 package by extracting fitted GAM responses and plotting them against observed data.

#### Differential abundance analysis

To gain insight into how SynCom inoculation reshaped seedling microbial communities, we identified differentially abundant ASVs between each individual SynCom treatment and the non-inoculated control group. To maximize the robustness of our differential abundance analysis and minimize algorithm-specific biases, we implemented a stringent consensus approach requiring an ASV to be significantly enriched or depleted in at least three out of four independent statistical methods. Seedling samples were first filtered using a prevalence threshold of 3% to retain low-prevalence taxa while allowing detection of SynCom-specific strains. Then, each SynCom was tested against the control (n = 16). Differential abundance testing was performed using the DAtest package (Version 2.8.0 [50]), which selects multiple statistical methods and ranks them based on performance metrics. For each method, the average performance score of each method was calculated for each condition, and the four best-performing methods were selected. For the *gyrB* dataset, the selected methods were LIMMA-ARL, EdgeR (exact test with TMM normalization), log-transformed t-test, and permutation tests. For ITS, the selected methods were EdgeR, Quasi-Poisson GLM, LIMMA - ALR and MgSeq Feature.

#### Multi-kingdom co-occurrence network

To elucidate the ecological mechanisms underlying the distinct structural partitions observed with the clustering approach on the bacterial dataset, we constructed three multi-kingdom co-occurrence networks: (i) A global network including all bacterial and fungal ASVs detected across all seedling samples, (ii) a Cluster 1-specific, and (iii) a Cluster 2-specific network. The two clusters are based on SynCom impact on seedling native microbiota, Cluster 1 encompass SynComs inducing community shit and Cluster 2 SynCom with low or no impact. Although clustering was primarily defined by bacterial differential abundance profiles, the full fungal dataset was incorporated into each network to capture interkingdom associations, with particular emphasis on inoculated strains and differentially abundant bacteria. Networks were constructed using the multi domain SPIEC-EASI method [51], which is robust against compositional data by using clr-transformation to remove the unit-sum constraint of compositional data, as proposed by Aitchison [52]. Each network was constructed with a lambda ratio of 1×10^-4^, with 80 lambda values over 999 cross-validations, while neighborhood selection was performed using the Meinshausen and Bühlmann method (‘mb’). Models were selected using the Stability Approach to Regularization Selection (StARS) algorithm with a stability threshold of 0.05, which was assessed for each network construction. The networks were visualized using Gephi [53] and their topological properties were calculated using the igraph package in R [54]. To allow comparison between networks with different numbers of nodes, nodes degree was normalized by dividing their degree by n-1, where n is the total number of nodes in each network. Betweenness centrality was normalized by its theoretical maximum in each graph using the igraph formula 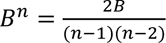 where *B^n^* is the normalized betweenness, B is the absolute betweenness and n is the number of nodes.

In order to assess if inoculated strains preferentially interacted with each other or with non-inoculated strains, assortativity coefficients were calculated following Newman 2003 [55]. This coefficient ranges from − 1 (disassortative mixing, where nodes preferentially connect to dissimilar nodes) and 1 (assortative mixing, where nodes preferentially connect to similar nodes). The assortativity coefficient was based on the inoculation status category of nodes (inoculated or not). We also calculated the proportion of connection each inoculated strain had with other inoculated strains to observe this interaction at node level. Seemingly, domain-specific interaction preferences were assessed by calculating, for each inoculated strain, the proportion of connections to bacterial *versus* fungal nodes.

### Statistical analysis

Statistical analysis was conducted after assessing data distribution using the Shapiro-Wilk test. None of the investigated data followed normal distribution, resulting in use of the Kruskal–Wallis test [56] of R (BH; α = 0.05). When multiple conditions were tested simultaneously, the non-parametric post-hoc test of Dunn (BH; α = 0.05 ; [57]) was conducted.

## Results

### 1) Alginate encapsulation of synthetic microbial communities enhances seed colonization and outcompetes native microbiota

To ensure a sufficient propagule size for effective SynCom inoculation, we targeted a concentration of 10^5^ microorganisms per seed for both bacterial and fungal taxa. This concentration was chosen to ensure a higher microbial biomass than non-inoculated seeds as well as its feasibility for fungal spore collection as part of inoculum preparation. Four distinct inoculation methods were evaluated on *Brassica napus* seeds: soaking in water, 10 mM MgCl_2_, 5% glycerol, and coating with 2.4% sodium alginate. Among these, alginate coating consistently resulted in the highest microbial loads across all taxa (Fig. 2A). Bacterial load increased by approximately 2 logs (∼85-fold), achieving 1.26×10^6^ bacteria per seed compared to 1.26×10^4^ bacteria per seed with water. Yeast abundance increased by almost 3 logs (∼284-fold), from 4.0×10^2^ yeast per seed in water to 1.14 × 10^5^ in alginate. The most pronounced effect was observed for filamentous fungi, with an increase of approximately 5-log (∼74,475-fold), reaching 2.17 × 10^7^ conidia per seed compared to 2.92 × 10^2^ with water. In the absence of microbial inoculation, both alginate coating and glycerol treatments led to a slight but statistically significant increase in germination rate compared to the control (Fig. 2B; Dunn’s test with Benjamini-Hochberg correction, p < 0.05). Dissolution of the alginate matrix enabled recovery of viable microorganisms, confirming its suitability for downstream assessment of seed colonization. Collectively, these results identify alginate encapsulation as an efficient and robust method for SynCom delivery onto seeds.

**Fig. 2:**
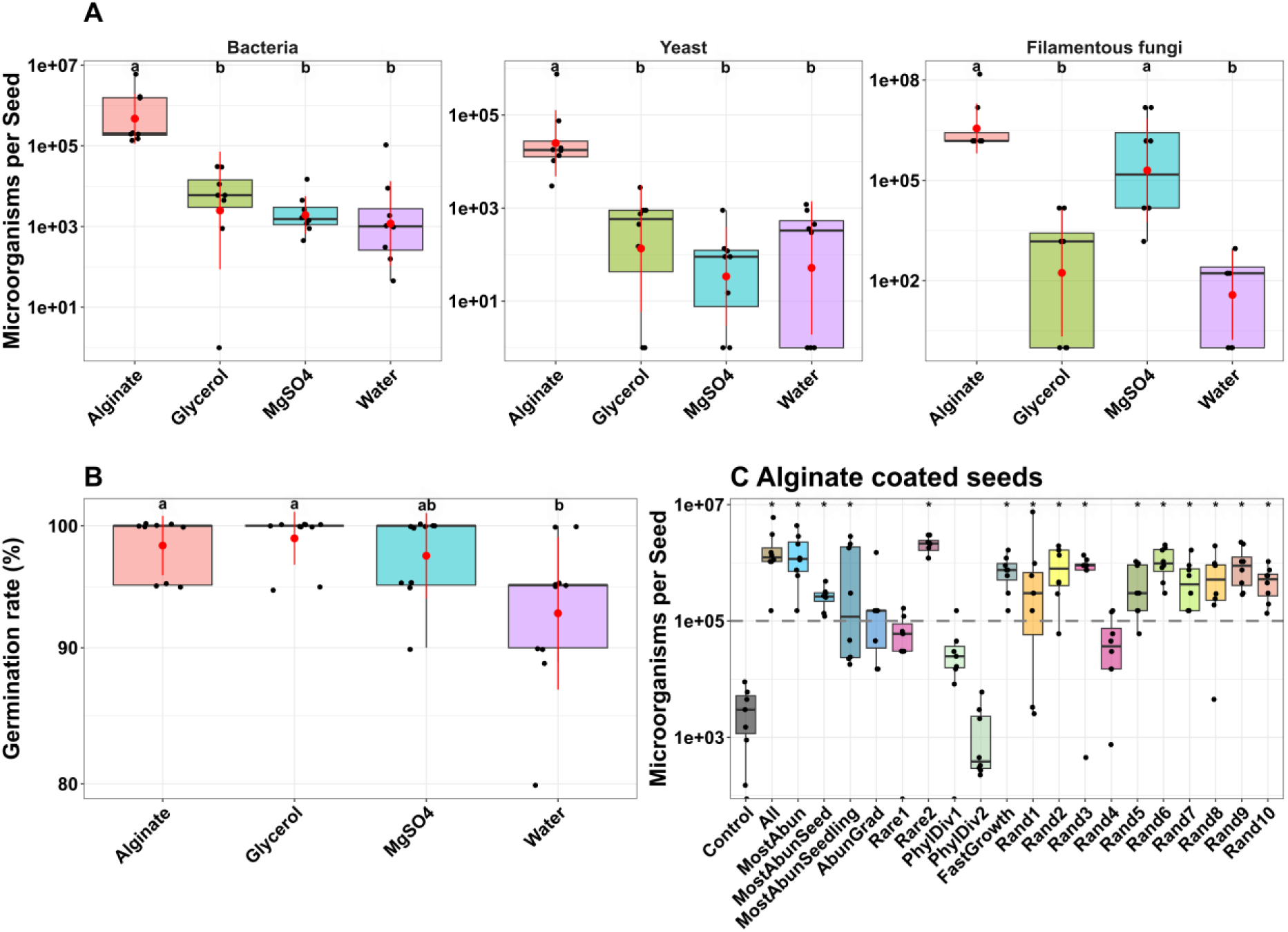
Evaluation of seed soaking and coating strategies and subsequent synthetic community (SynCom) colonization efficiency on *Brassica napus*. (A) Impact of four inoculation methods on seed microbial population sizes using representative single bacterial, yeast, or filamentous fungal strains. (B) Influence of inoculation matrices on seed germination rates in the absence of microbial strain inoculation. For both (A) and (B), distinct letters indicate statistically significant differences between treatment groups (Dunn’s post-hoc test with Benjamini-Hochberg correction, p < 0.05). (C) Total microbial population sizes recovered from individual seeds inoculated *via* sodium alginate encapsulation across 20 distinct SynCom design. Asterisks (*) indicate statistically significant differences compared to the non-inoculated control group (Pairwise comparisons using Dunn’s many- to-one test with Benjamini-Hochberg correction, p < 0.05).

Following optimization of the inoculation method, a total of 20 distinct multi-kingdom SynComs were designed (Fig. 1A) and inoculated with alginate coating on seeds. Compositional distance calculations were used to assess the overall differences in SynCom composition (Fig. 1B). SynCom with expected similar composition (MostAbun, MostAbunSeed and MostAbunSeedling) clustered close to each other while randomly designed SynCom were disseminated along both axis. Post-inoculation, the total microbial load was quantified across eight individual seeds per condition to verify whether the target density of 10^5^ cells per seed was reached (Fig. 2C). Non-inoculated control seeds (alginate only) exhibited a mean concentration of 3.13 × 10^3^ microorganisms per seed. In comparison, 15 of the 20 SynComs reached significantly higher microbial load per single seed relative to non-inoculated controls, with mean concentrations ranging from 2.66 × 10^5^ (MostAbunSeed) to 2.09 × 10^6^ microorganisms per seed (Rare2). Conversely, the five SynComs PhylDiv1, PhylDiv2, Rare1, Rand4, and AbunGrad, yielded densities ranging from 1.58 × 10^3^ to 2.72 × 10^5^ microorganisms per seed, and did not differ from controls. Overall, only four SynComs (PhylDiv1, PhylDiv2, Rare1, and Rand4) failed to reach the target inoculation threshold on seeds.

To determine whether the observed microbial richness reflected the initial inoculum or the persistence of native microbiota, we evaluated the number of ASVs across treatments (Observed richness). Because individual non-inoculated control seeds did not yield sufficient microbial DNA for robust metabarcoding amplification, we analyzed four replicates of 30 pooled seeds each to characterize the native microbiota. In these control pools, mean richness reached 63.25 bacterial ASVs and 33.50 fungal unique ASVs. For bacterial richness (Supplementary Fig. 3A), nine SynComs (All, Rare1 and Rare2, PhylDiv2, FastGrowth, Rand4, Rand5, Rand7, and Rand10) did not differ significantly from control conditions (Pairwise comparisons using Dunn’s many-to-one test with BH correction, p > 0.05). Interestingly, although the “All” SynCom formulation included 24 bacterial strains in the initial inoculum, only 19 ASVs were detected on seeds after inoculation. In contrast, SynCom that significantly differed from the control pools exhibited reduced bacterial richness, with values ranging from 6.75 to 13 ASVs. Contrary to bacteria, fungal richness remained stable across all inoculated conditions (Supplementary Fig. 3B) and did not significantly differ from control pools (Pairwise comparisons using Dunn’s many-to-one test with BH correction, p > 0.05). The highest mean fungal richness was observed in the SynCom All with 51.5 unique ASVs. Together, these results indicated that SynCom inoculation impacted bacterial richness of seeds but did not completely overtake native microbiota while fungal richness remained largely unaffected. The comparable fungal richness between inoculated seeds and pooled controls suggests that native fungal taxa persist following inoculation. This may be explained by the absence of seed surface sterilization prior to inoculation, which could allow native fungal communities, potentially more resistant to water rinsing than bacteria, to be maintained.

To determine whether SynCom inoculation altered seed microbial composition and structure and to assess its similarity with the initial inoculum profiles, we evaluated beta diversity using Bray-Curtis dissimilarity matrices. For bacterial communities (Fig. 3A), principal coordinate analysis (PCoA) revealed clear clustering by SynCom composition, with tight co-localization between each specific inoculum and its corresponding inoculated seeds. PERMANOVA confirmed that SynCom composition was the primary driver of bacterial beta diversity, accounting for 79% of the total community variance (PERMANOVA, 999 permutations, R^2^ = 0.79, p < 0.001). For fungal communities (Fig. 3B), the initial inocula and the resulting inoculated seeds were separated along the first axis of the ordination. Nevertheless, SynCom treatment remained a significant factor, accounting for 35% of the variance (PERMANOVA, 999 permutations, R^2^ = 0.35, p < 0.001). These results indicated that SynCom inoculation strongly modulated bacterial communities on seeds, whereas its effect on fungal communities is more moderate. Furthermore, partial overlap in fungal strains compositions across SynCom formulations may contribute to the reduced clustering observed between distinct SynCom treatments.

**Fig. 3:**
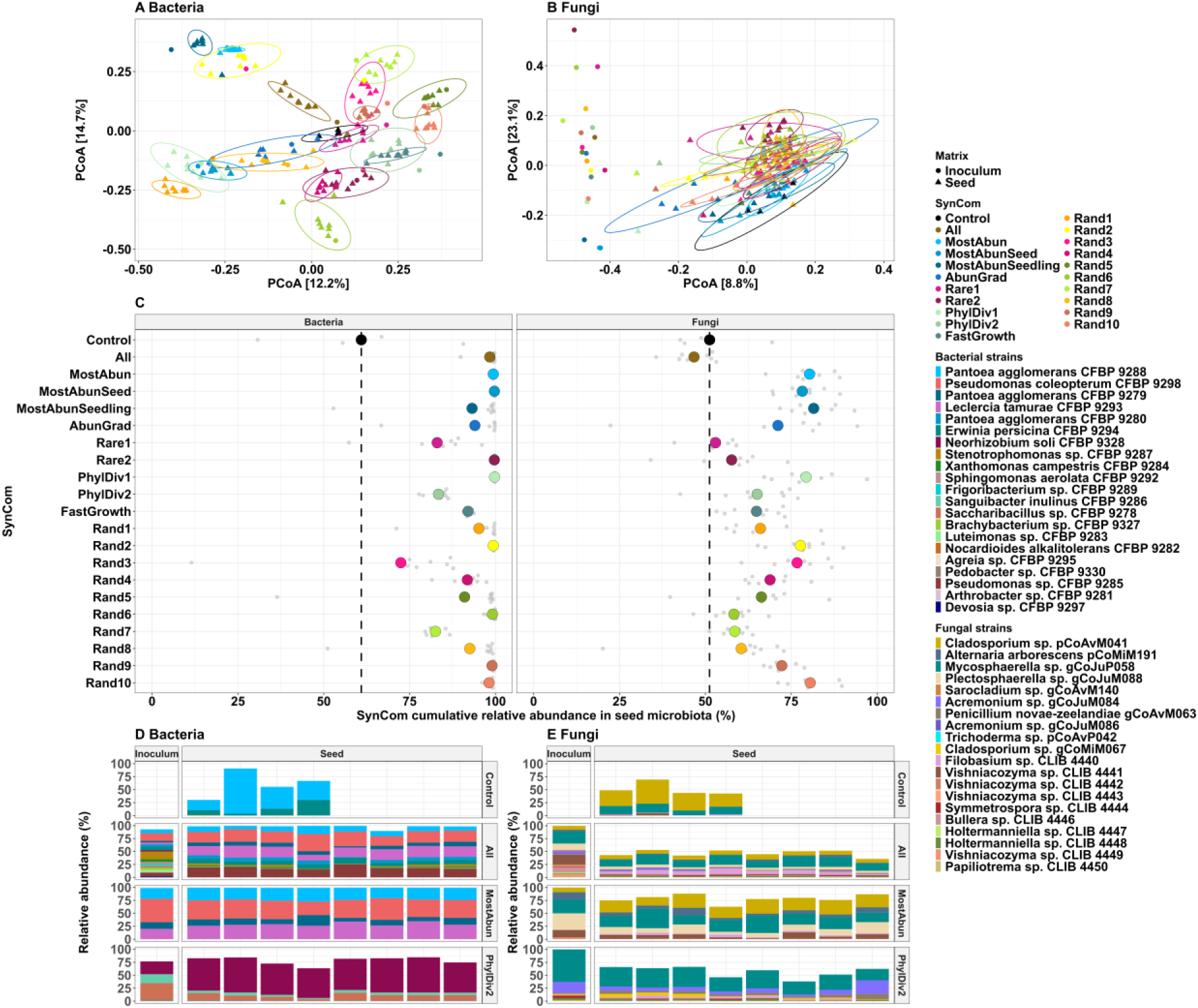
Beta diversity, community composition, and relative contribution of introduced SynComs to the *Brassica napus* seed microbiota. (A–B) Principal Coordinate Analysis (PCoA) based on Bray-Curtis dissimilarity matrices (log(x+1) transformed) evaluating (A) bacterial and (B) fungal community structures across control and SynCom- inoculated seeds and their respective initial inocula. For SynCom treatments and inocula, individual points represent single seeds or independent inoculum samples, whereas control points represent pooled batches of 30 non-inoculated seeds. (C) Relative contribution (%) of inoculated SynCom strains to the total bacterial and fungal seed microbiota, calculated by summing the relative abundance of tracked inoculum-specific ASVs. (D–E) Taxonomic profiling of successfully tracked (D) bacterial and (E) fungal inoculum strains, illustrating relative abundances within the initial inocula and the resulting seed microbiota across the Control, All, MostAbun, and PhylDiv2 SynCom designs as example, all SynCom profiles are presented in Sup. Fig.3. Only inoculated strains are represented, thus blank space represent remaining non inoculated microbiota (i.e native taxa).

To quantify the extent to which the introduced SynComs successfully colonized the seed matrix, we tracked the specific ASVs assigned to the inoculated strains. Across all datasets, only two inoculated strains were not detected after inoculation: *Bacillus pumilus* CFBP 9299 and *Holtermanniella* sp. CLIB 4445, for which no sequence reads were recovered. The cumulative relative abundance of detected inoculum-derived ASVs was used to estimate the overall SynCom contribution to the seed microbiota (Fig. 3C). Non-inoculated control seeds already exhibited a high baseline abundance of SynCom-associated taxa, representing 60.9% of the bacterial communities and 51.2% of the fungal communities. This finding is biologically consistent, as the strains selected for SynCom construction were originally isolated from this exact *B. napus* seed and seedling lot. Following inoculation, the average SynCom contribution increased to 93.7% for bacteria and 68.2% for fungi. This is further supported by taxonomic composition analyses of inoculum and seeds. In control seeds, bacterial communities were dominated by *Pantoea agglomerans* CFBP 9288 and *Erwinia persicina* CFBP 9294, with only minor contributions of other taxa such as *Pseudomonas coleopterum* CFBP 9298, *Neorhizobium soli* CFBP 9328, *Stenotrophomonas* sp. CFBP 9287, *Xanthomonas campestris* CFBP 9284, *Sphingomonas aerolata* CFBP 9292 and *Sphingomonas* sp. CFBP 9329. Similarly, fungal communities were largely dominated by *Cladosporium* sp. pCoAvM041 and *Mycosphaerella* sp. gCoJuP058, alongside low-abundance taxa. In contrast, inoculated seeds showed a pronounced shift in taxonomic composition. For example, seeds inoculated with the SynCom “All” harbored 21 different bacterial and 18 fungal inoculated strains, with taxonomic profile closely mirroring those of corresponding inocula. Same patterns were observed for MostAbun and PhylDiv2, for both bacterial and fungal communities (Fig. 3D and Fig. 3E). Overall, seed taxonomic profiles matched their respective inoculum, a more pronounced effect for bacterial communities (Supplementary Fig. 3 C and E) than for fungal communities (Supplementary Fig. 3D and Fig. 3E). This substantial enrichment demonstrates that the SynCom successfully outcompeted and displaced the remaining native microbiota. This effect is particularly pronounced for bacterial communities, whereas fungal communities, although influenced, retain a greater contribution from native taxa

### 2) Seedling colonization is driven by SynCom composition

Following 15 days of growth in non-sterile potting soil, eight phenotypically normal seedlings seeds per condition (inoculated and control) were harvested to characterize the seedling microbiota. Concurrently, phenotypic effects of SynCom inoculation were evaluated across all remaining seedlings (n = 76) by monitoring germination rate, hypocotyl length, and the proportion of morphologically normal seedlings. Seed germination was not significantly altered by SynCom inoculation (Supplementary Fig. 4A; Kruskal-Wallis rank-sum test, p = 0.85). However, hypocotyl length (Supplementary Fig. 4B) was significantly reduced by four SynComs (Rare2, PhylDiv1, PhylDiv2, and FastGrowth) and significantly increased by five SynComs (MostAbun, Rand5, Rand7, Rand9, and Rand10; Pairwise comparisons using Dunn’s many-to-one test with Benjamini-Hochberg [BH] correction, p < 0.05). The proportion of normal seedlings (Supplementary Fig. 4C) was significantly reduced in five SynComs (All, MostAbun, Rand6, Rand7 and Rand8; Pairwise comparisons using Dunn’s many-to-one test with Benjamini-Hochberg [BH] correction, p < 0.05). Thus, the SynCom inoculation does not affect seed germination but significantly influences seedling phenotype, with both beneficial and detrimental effects depending on community composition.

The colonization of seedlings by inoculated strains was monitored by quantifying the cumulative relative abundance of the SynCom to the microbiota of emerged seedlings (Fig. 4A). In non-inoculated control seedlings, the baseline relative abundance of SynCom-associated taxa was low for both bacteria (6.9%) and fungi (2.6%), indicating limited natural transmission of these strains from seed to seedlings. Bacterial colonization of seedlings varied markedly depending on SynCom composition. Among the 20 SynComs tested, 14 exhibited higher colonization than control seedlings, including five *a priori* designed and nine randomly assembled SynComs. The highest colonization efficiencies were observed for SynCom Rand8 (45.7%) and Rand9 (42.0%), followed by the SynCom All (38.4%) and the three SynComs based on high natural abundance (MostAbun, MostAbunSeed and MostAbunSeedling) (Fig. 4A). Conversely, fungal colonization showed overall lower relative abundance compared to bacteria. However, all SynComs resulted in a higher fungal ASVs abundance than in control seedlings. The most effective SynComs for fungal colonization were Rand3 (36.6%) and Rand9 (26.3%), followed by PhylDiv1 (22.5%) and Rand1 (21.0%). A weak but statistically significant positive correlation was observed between bacterial and fungal colonization success within the same individual seedlings (R² = 0.11, p =2.8×10^-5^; Supplementary Fig. 5A), suggesting limited co-dependence between kingdoms during colonization. To evaluate the impact of SynCom design strategies, we compared colonization efficiency between *a priori* and randomly assembled communities (Fig. 4B). For bacterial colonization, no significant difference was observed between the *a priori* (18.6%) and random (20.6%) design strategies (Wilcoxon-Mann-Whitney test, p = 0.25). In contrast, fungal colonization was significantly higher in randomly assembled SynCom (15.9%) compared to the *a priori* designed SynComs (11.6%) (Wilcoxon-Mann-Whitney test, p < 0.05). Overall, SynComs establishment remained partial for both bacterial and fungal communities. Nevertheless, colonization success was strongly influenced by SynCom composition, and notably, assembly based on high natural abundance outperformed the other designs.

**Fig. 4:**
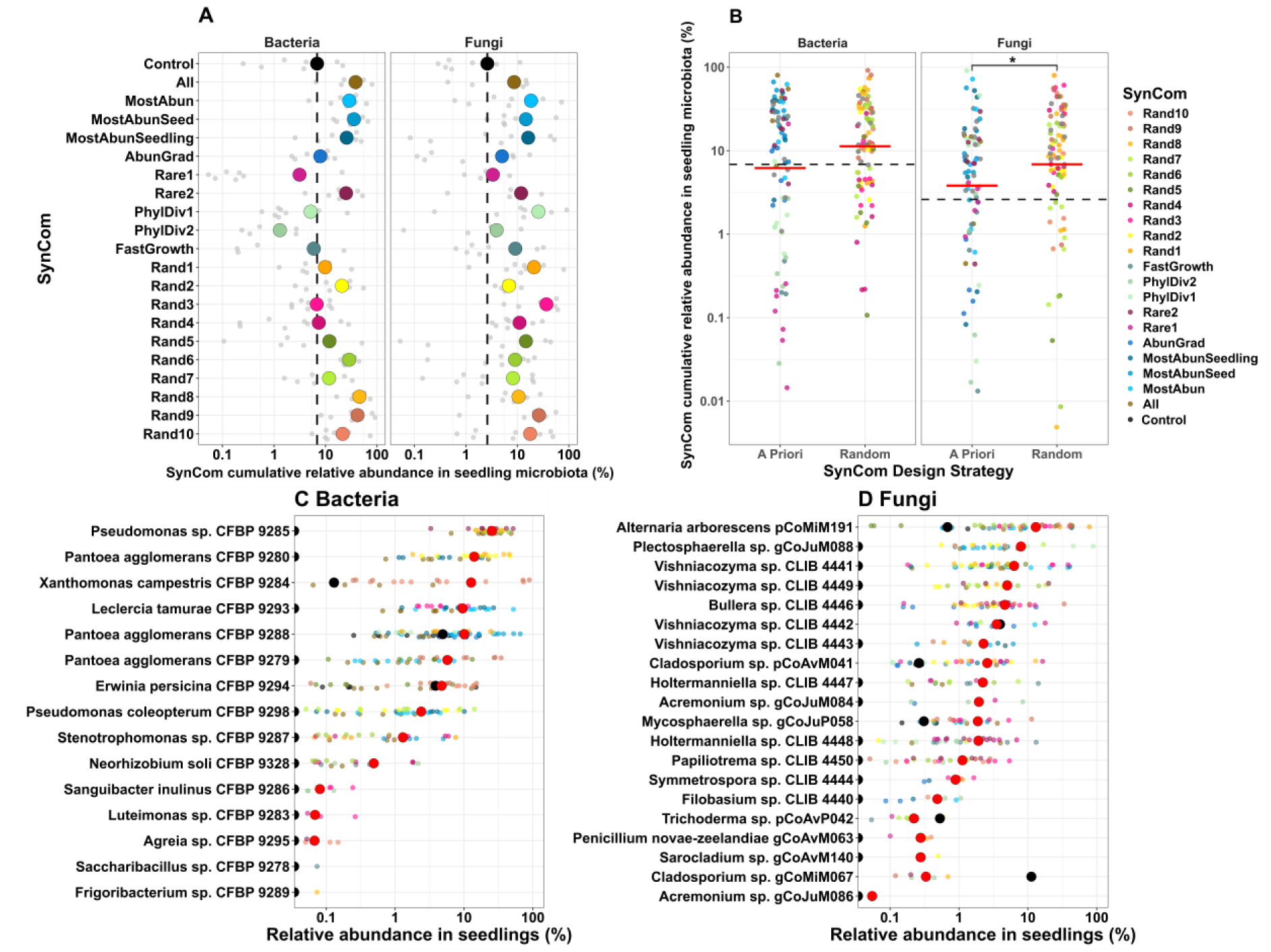
Characterization of SynCom colonization efficiency and strain-level relative abundances in the *Brassica napus* seedling microbiota. (A) Cumulative relative abundance (%) of inoculated SynCom members to the bacterial and fungal communities of emerged seedlings. Light gray points indicate individual seedling replicates (n = 8), colored points designate treatment means, and the horizontal black dashed line indicates baseline recovery in the non-inoculated control group. (B) Comparative analysis of cumulative SynCom colonization between the *a priori* and random community-design frameworks. Solid red horizontal lines indicate group means, and black dashed lines indicate the mean baseline value of the control group. Asterisks (*) denote statistically significant differences between design strategies (Wilcoxon-Mann-Whitney test, p < 0.05). (C–D) Relative abundance profiles of successfully tracked (C) bacterial, (D) filamentous fungal and yeast strains within the seedling microbiota. Smaller colored points represent individual seedling values for each specific SynCom treatment. Red points indicate the mean relative abundance across all replicates for a given strain, while black points indicate its mean baseline abundance across eight non-inoculated control seedlings.

To evaluate the colonization potential of each inoculated strain, we quantified their relative abundance in emerged seedling. Among the bacterial strains (Fig. 4C), 16 out of 24 were successfully detected in seedling microbiota. Eight strains (CFBP 9292, CFBP 9327, CFBP 9282, CFBP 9299, CFBP 9330, CFBP 9296, CFBP 9281 and CFBP 9297) were not detected. Only three bacterial strains were found in control seedlings: CFBP 9284, CFBP 9288 CFPB 9294, explaining the low natural colonization of control seedlings. Strains affiliated to genera *Pseudomonas*, *Pantoea*, *Xanthomonas*, *Leclercia* and *Erwinia* exhibited the highest colonization success. In particular, *Pseudomonas* sp. CFBP 9285 showed the greatest colonization, reaching a mean relative abundance of 21.4%. The remaining detected bacterial strains persisted at low abundances, ranging from 0.03% to 1.2%. For fungal strains (Fig. 4D), all filamentous fungi were successfully recovered from the seedlings, along 10 out of 11 yeast strains with six (five filamentous fungi and 1 yeast) found in control condition. Fungal colonization was primarily driven by filamentous taxa, notably *Alternaria arborescens* pCoMiM191 (10.6%) and *Plectosphaerella* sp. gCoJuM088 (7.5%), followed by the yeast *Vishniacozyma* sp. CLIB 4441 (5.7%). Tracking individual strain performance across multiple, compositionally distinct SynComs revealed that seed to seedling transmission success is strongly strain-dependent. A subset of strains consistently behaved as dominant colonizers across contexts, whereas others failed to establish or persisted only at marginal abundances.

### 3) Strain Traits Drive Seedling Colonization Success

To elucidate the relationships between specific microbial traits and seedling colonization efficiency, we used Generalized Additive Models (GAMs) to create a fungal and a bacterial multivariate model using eight predictive traits as fixed effects and strain identity as a random effect. The bacterial GAM explained 92.10% of the total deviance in log-transformed strain abundance in seedlings. The random effect of strain identity accounted for 43.67% of the model variance, indicating strong strain-specific differences in colonization, although it was not significant (p = 0.085), likely reflecting limited statistical power. Among the eight fixed predictors, three variables were significant. Strain abundance on inoculated seed (Fig. 5A, p = 0.029) and assembled genome size (Fig. 5B, p < 0.001) showed significant linear relationships with seedling colonization (edf = 1), contributing respectively to 20.08% and 14.88% of the explained deviance. Lag phase duration, exhibited a significant non-linear relationship with seedling establishment (Fig. 5C, edf = 4.37, p < 0.001) accounting for 5.42% of the total deviance in the model. The apparent offset between partial effect curves and raw data points reflects the conditional nature of prediction derived from multivariate GAMs, where predictors act simultaneously rather than independently. Overall, these results indicate that bacterial seedling colonization is primarily driven by high initial abundance on seeds and larger genome size, with shorter lag phase providing an additional but more limited advantage.

**Fig. 5:**
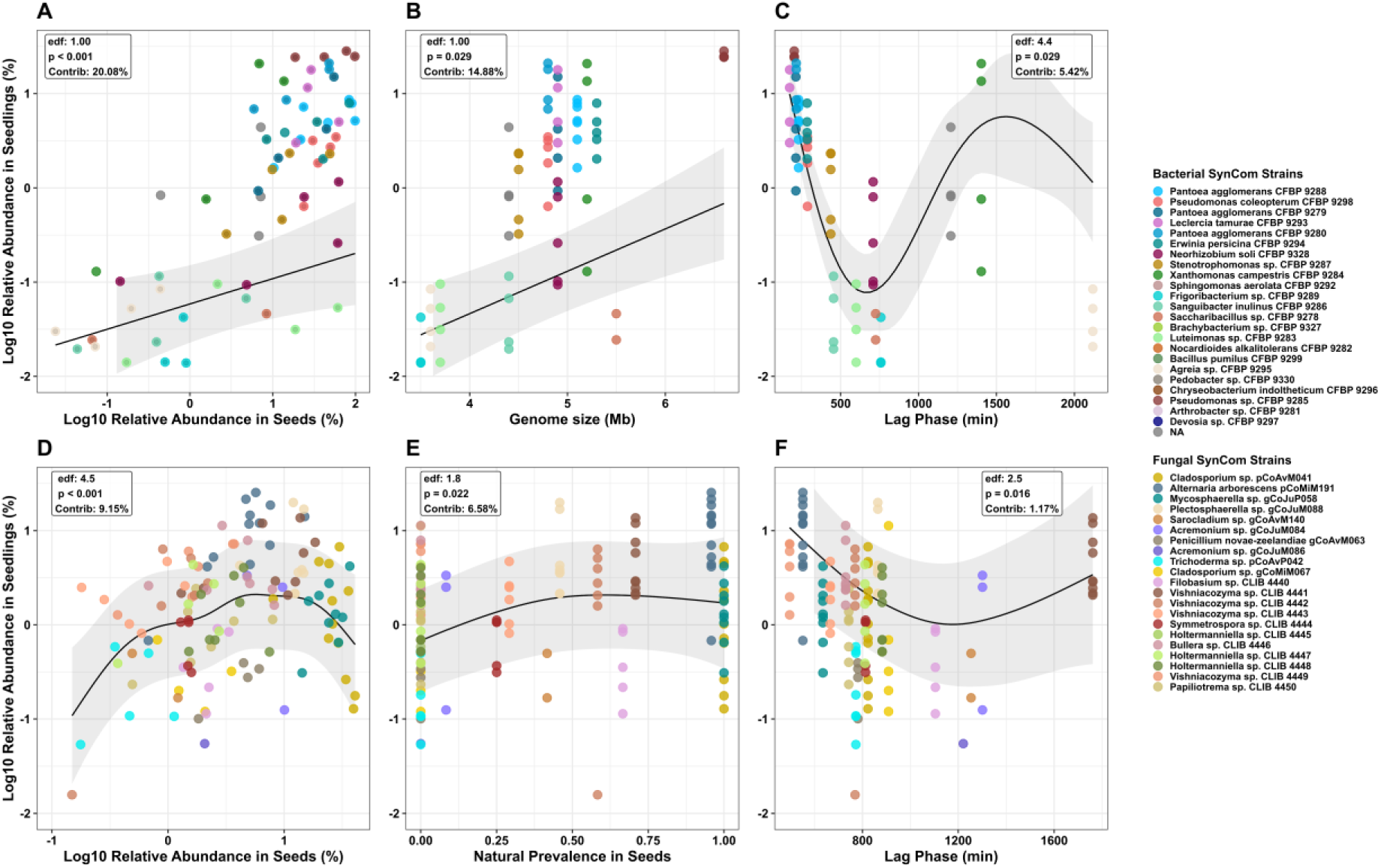
Generalized Additive Model (GAM) partial effects of significant phenotypic and ecological traits driving strain-level seedling colonization. Predicted partial effects (solid blue lines) and corresponding 95% confidence intervals (shaded gray regions) of significant microbial traits predicting the log10-transformed relative abundance of inoculated strains in *Brassica napus* seedlings. For each parameter, the estimated degrees of freedom (edf), p, and relative deviance contribution (%) to the multivariate model are displayed. (A–C) Bacterial models predicting seedling log-abundance based on (A) log10 relative abundance on inoculated seeds, (B) assembled genome size, and (C) *in vitro* lag phase duration. (D–F) Fungal models predicting seedling log- abundance based on (D) log10 relative abundance on inoculated seeds, (E) native seed prevalence, and (F) *in vitro* lag phase duration.

The fungal GAM explained 69.30% of the total deviance. In contrast to bacteria, the random effect of strain identity was statistically significant (p = 0.027) and accounted for the majority of the explained variance (59.77%). Three fixed effects significantly influenced fungal colonization. Relative abundance on inoculated seeds showed a strong non-linear relationship (Fig. 5D, p < 0.001, edf = 4.5) explaining 9.15% of the model deviance. Natural ASV prevalence on seeds also displayed a significant non-linear effect (Fig. 5E, p = 0.022, edf = 1.8), contributing 6.58%. Lag phase duration had a weaker but significant non-linear effect (Fig. 5F, p = 0.016, edf = 2.5), explaining 1.17% of the deviance. Together, these results show that fungal colonization success is primarily associated with high abundance and prevalence on seeds, both following nonlinear relationships, while a shorter lag phase conferring a moderate additional benefit.

### 4) SynCom inoculation modulates seedling microbiota assembly and community structure

To compare the microbial communities of the seedling between conditions, we analyzed beta diversity using Bray-Curtis dissimilarity matrices across seedling and bulk soil samples. Bacterial community composition was significantly driven by SynCom treatment (Fig. 6A), which explained 33% of the total variance (PERMANOVA, 999 permutations, R^2^ = 0.33, p < 0.001), resulting in a clear clustering along the primary axis of the ordination. To objectively define the number and structure of these bacterial assemblages, k-means clustering was applied on the first two axes of the Principal Coordinate Analysis (PCoA). The optimal number of clusters was determined using the NbClust package in R by evaluating 26 distinct diagnostic indices. A majority consensus of 10 indices supported k = 2 as the optimal partition (Table S12). This clustering significantly explained the variation observed in the bacterial community (PERMANOVA, 999 permutations, R^2^ = 0.17, p < 0.001). The clustering approach was also tested on fungal communities (Supplementary Fig. 6) and revealed 3 overlapping clusters with a distribution of the same SynCom in multiple clusters. For these reasons, only bacterial community clusters were investigated.

**Fig. 6:**
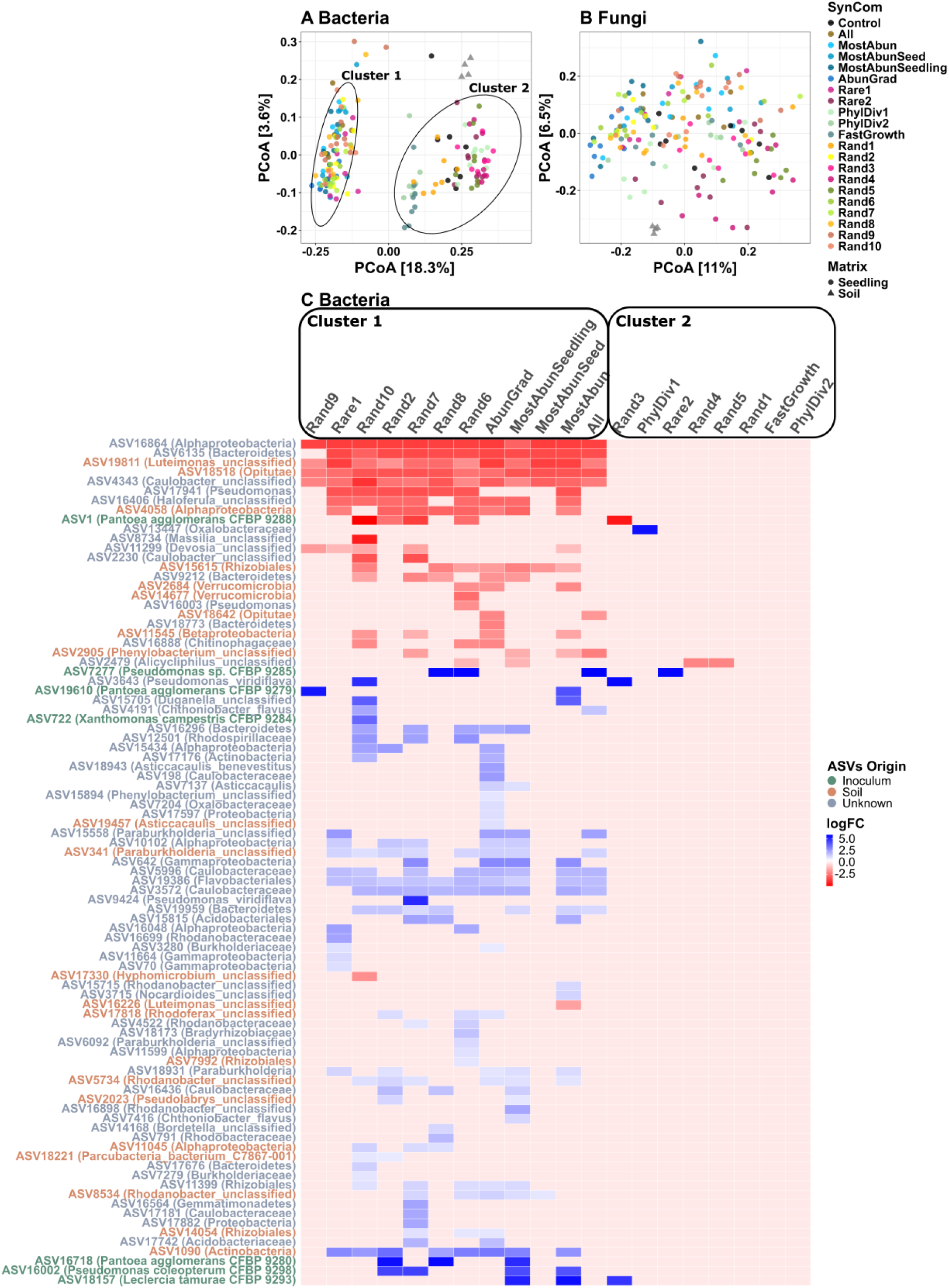
Community-level beta diversity and differential bacterial ASV recruitment in the *Brassica napus* seedling microbiota. (A–B) Principal Coordinate Analysis (PCoA) based on Bray-Curtis dissimilarity matrices (log(x+1) transformed) for (A) bacterial and (B) fungal communities across bulk soil, non-inoculated control seedlings, and SynCom-inoculated seedlings. Cluster overlays are derived from k-means clustering partitioning using the NbClust package in R based on Euclidean distances. (C) Hierarchical clustering heatmap of differentially abundant bacterial ASVs in inoculated seedlings relative to the non-inoculated control group, identified via a consensus of at least three out of four distinct statistical algorithms (LIMMA-ARL, EdgeR exact-TMM, log-transformed t-test, and permutation tests). Row annotations indicate the environmental or experimental origin of each ASV.

For bacteria, Cluster 1 was clearly separated from non-inoculated control and included 12 SynComs: six randomly assembled communities (Rand2, Rand6, Rand7, Rand8, Rand9 and Rand10), and six *a priori* designs, including those based on high natural abundance (MostAbunSeedling, MostAbunSeed, MostAbun), the high richness community (All), the abundance gradient (AbunGrad), and one rare taxa community (Rare1). Conversely, Cluster 2 was positioned closer to the bulk soil samples and included the non-inoculated control, four random SynComs (Rand1, Rand3, Rand4, Rand5), the second rare SynCom (Rare2), both phylogenetic diversity based communities (PhylDiv1 and 2), and the fast-growing community (FastGrowth). Fungal community structure displayed weaker clustering (Fig. 6B), although SynCom treatment remained a significant driver of fungal community composition, explaining 22% of the variance (PERMANOVA, 999 permutations, R^2^ = 0.22, p < 0.001). Bulk soil samples clustered independently and remained separated from all seedling samples. Pairwise comparisons revealed that only three SynComs significantly altered fungal community composition relative to the non-inoculated controls (Pairwise PERMANOVA, false discovery rate adjusted p < 0.005). Notably, all three belonged to the *a priori* design based on high native abundance: Most Abun (R^2^ = 0.13), MostAbunSeedling (R^2^ = 0.16), and AbunGrad (R^2^ = 0.22). Taken together, these findings indicate that SynCom inoculation strongly reshapes bacterial community assembly, whereas fungal community assembly is less responsive to SynCom inoculation.

To identify which taxa were responsible for the community shift in seedlings, we used a differential abundance analysis between each SynCom treatment and the non-inoculated control group with four independent statistical methods, retaining ASVs being detected in at least 3 methods. No fungal ASV met the threshold, confirming that the seedling fungal microbiota remained resistant to specific SynCom-induced shifts. Conversely, 89 bacterial ASVs were identified as differentially abundant across 17 of the 20 SynCom treatments (Fig. 6C). Among these, seven corresponded inoculated strains: *Pantoea agglomerans* CFBP 9288*, Pseudomonas* sp. CFBP 9285, *Pantoea agglomerans* CFBP 9279, *Xanthomonas campestris* CFBP 9284*, Pantoea agglomerans* CFBP 9279*, Pseudomonas coleopterum* CFBP 9298, and *Leclercia tamurae* CFBP 9293. Most of these strains showed significant enrichment in the treatments in which they were introduced. In contrast, *Pantoea agglomerans* CFBP 9288 exhibited a unique pattern, being consistently depleted in treatments where it was not inoculated, likely reflecting its high natural abundance in control seedlings. The remaining 82 ASVs corresponded to non-inoculated native taxa. Taxonomic assignment revealed a strong dominance of *Pseudomonadota*, particularly within *Alphaproteobacteria* (35.4%; n= 29/82), *Gammaproteobacteria* (18.3%; n = 15/82), and *Betaproteobacteria* (17.1%; n = 14/82). Tracking their environmental origin indicated that only a minority of differentially abundant ASVs originated from bulk soil matrix (24.7%; n = 22) and non were identified a originating from native seed microbiota, whereas the majority could not be traced (67.4%; n = 60), suggesting recruitment from either unsampled source or rare seed or soil-derived taxa. Regression analyses revealed no significant correlation between the cumulative colonization efficiency of the inoculated strains and the number of differentially abundant native taxa, for either the bacteria (Supplementary Fig.5B, R^2^ = 0.102, p = 0.171) or fungi (Supplementary Fig. 5C, R^2^ = 0.063, p = 0.286). However, differential abundance profiles closely matched the k-means clusters established in Fig. 6A. This concordance indicates that community-level restructuring is not solely driven by the dominance of inoculated strains, but also depends on broader, SynCom-specific ecological interactions.

### 5) Interkingdom Co-occurrence Networks Reveal Structural Drivers of Community Clustering

To elucidate the ecological mechanisms underlying the distinct structural partitions observed between Cluster 1 (community shift) and Cluster 2 (no community shift), we constructed three multi-kingdom co-occurrence networks: (i) A global network including all bacterial and fungal ASVs detected across all seedling samples (Fig. 7A), (ii) a Cluster 1-specific (Fig. 7B), and (iii) a Cluster 2-specific network (Fig. 7C).

**Fig. 7:**
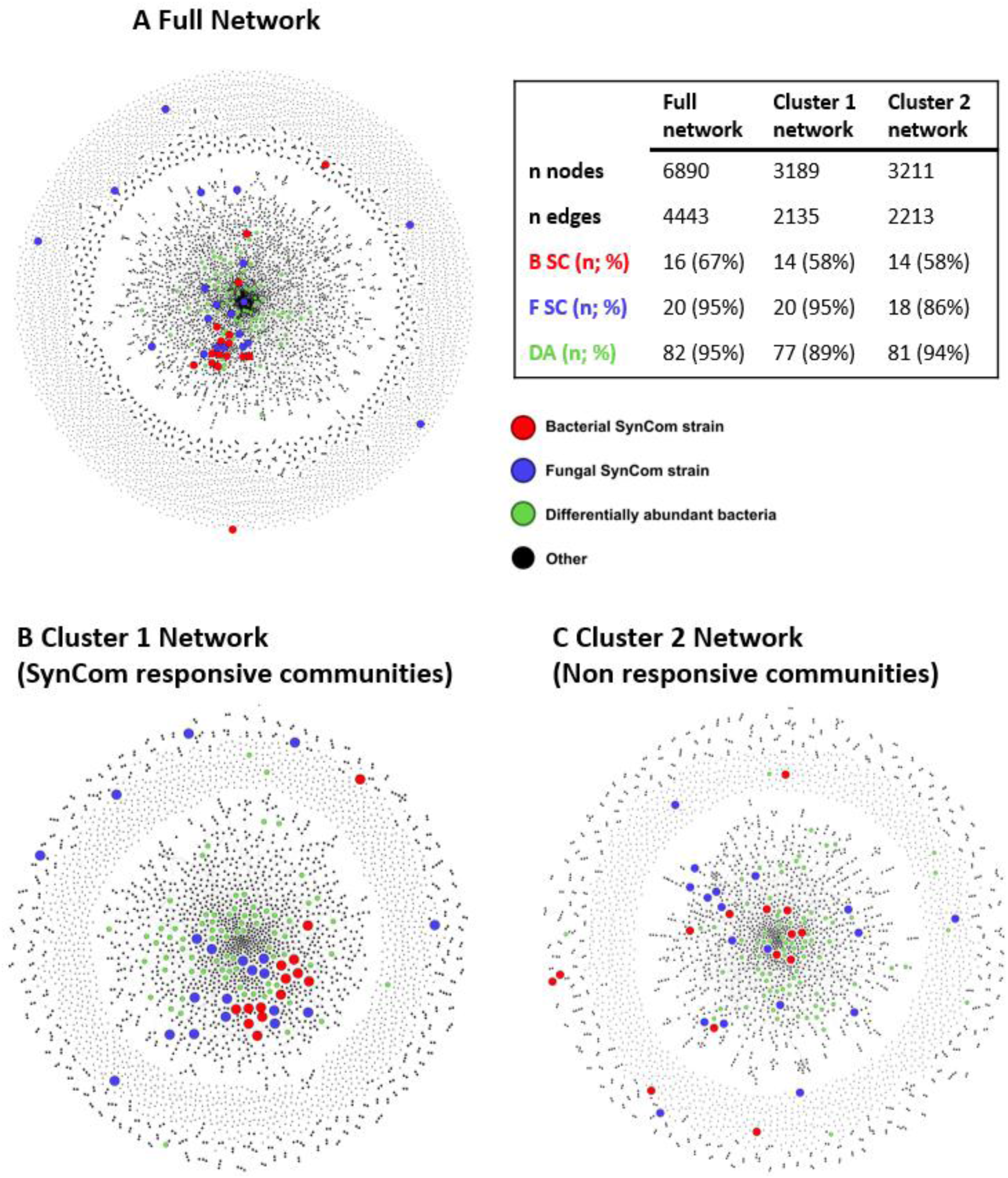
Topology of interkingdom co-occurrence networks in the *Brassica napus* seedling microbiota. Visualizations represent microbial networks constructed across (A) the full dataset including all seedling samples, (B) seedling samples within Cluster 1, and (C) seedling samples within Cluster 2. Nodes represent unique bacterial or fungal ASVs, while edges denote statistically significant co-occurrence relationships. Node coloring distinguishes introduced SynCom bacterial (red) and fungal (blue) strains, differentially abundant native ASVs identified in consensus profiling (green), and background native microbiota (black). The table encompasses the number of nodes and edges as well as the number and proportion of bacterial SynCom strain (B SC), fungal SynCom strain (F SC) and differentially abundant native ASVs (DA) across each network.

Although clustering was primarily defined by bacterial differential abundance profiles, the full fungal dataset was incorporated into each network to capture interkingdom associations, with particular emphasis on inoculated strains and differentially abundant bacteria.

Even though the networks differed in size (n = 6890, 3189 and 3211 nodes for full, Cluster 1 and Cluster 2 networks), they showed similar density (1.87 × 10^-4^, 4.2 × 10^-4^, and 4.29 × 10^-4^ for full, Cluster 1 and Cluster 2 networks) and similar normalized numbers of connections among taxa (0.64, 0.67 and 0.69 normalized degree for full, Cluster 1 and Cluster 2 networks). This pattern was consistent when considering the full networks as well as their largest connected component (giant component) and peripheral nodes (Table S13).

We then investigated the position of SynCom strains within each network. The giant component of the Cluster 1 network included a higher proportion of inoculated strains than Cluster 2 (79%, n = 27 strains, versus 72%, n = 23 strains; Supplementary Fig. 7A), with 20 strains shared between both clusters. Network centrality metrics revealed marked differences in SynCom strain integration between the two clusters (Fig. 8A). SynCom strains exhibited significantly higher normalized degree in the Cluster 1 network (normalized node degree = 1.23 × 10-3) compared to both the global network (normalized node degree = 6.25 × 10-4) and Cluster 2 (normalized node degree = 6.81 × 10-4 Dunn’s test with Benjamini-Hochberg correction, p < 0.05). This pattern was consistent with the analysis of the 100 most connected ASVs in each network (Supplementary Fig. 7C–E), where Cluster 1 network contained the highest number of SynCom strains (n = 10) and was the only network in which introduced fungal strains ranked among the most connected taxa. To assess whether inoculated strains acted as important connectors within the co-occurrence network, we calculated normalized betweenness centrality, which reflects how often a node lies on the shortest paths connecting other nodes (Fig. 8B). SynCom strains showed higher mean betweenness centrality in Cluster 1 (mean betweenness = 1.03 × 10-3) than in Cluster 2 (mean betweenness = 2.45 × 10-4; Dunn’s test with BH correction, p < 0.05), while not differing significantly from the global network (mean betweenness = 3.35 × 10-4). These results suggest that SynCom strains are more integrated and play a greater structural role in Cluster 1 network than in Cluster 2. To further characterize SynCom integration in seedling microbiota, we examined assortativity coefficients to test whether inoculated strains preferentially connected with each other rather than with native taxa. The Cluster 1 network showed the highest assortativity (r = 0.36), followed by the global (r = 0.28) and Cluster 2 (r = 0.25) networks, indicating a general tendency for SynCom strains to co-associate. However, node-level analysis (Fig. 8C) revealed that SynCom strains shared a lower proportion of their edges with other introduced strains in Cluster 1 (percentage of connections between inoculated strains = 7.3%) and in the global network (9.3%) than in Cluster 2 (28%; Dunn’s test with BH correction, p < 0.05). Across all networks, SynCom strains had a higher number of connections with native bacteria than with fungi (Fig. 8D). Notably, connections with fungi were significantly less frequent in Cluster 2 compared to the other networks (Dunn’s test with BH correction, p < 0.05).

**Fig. 8:**
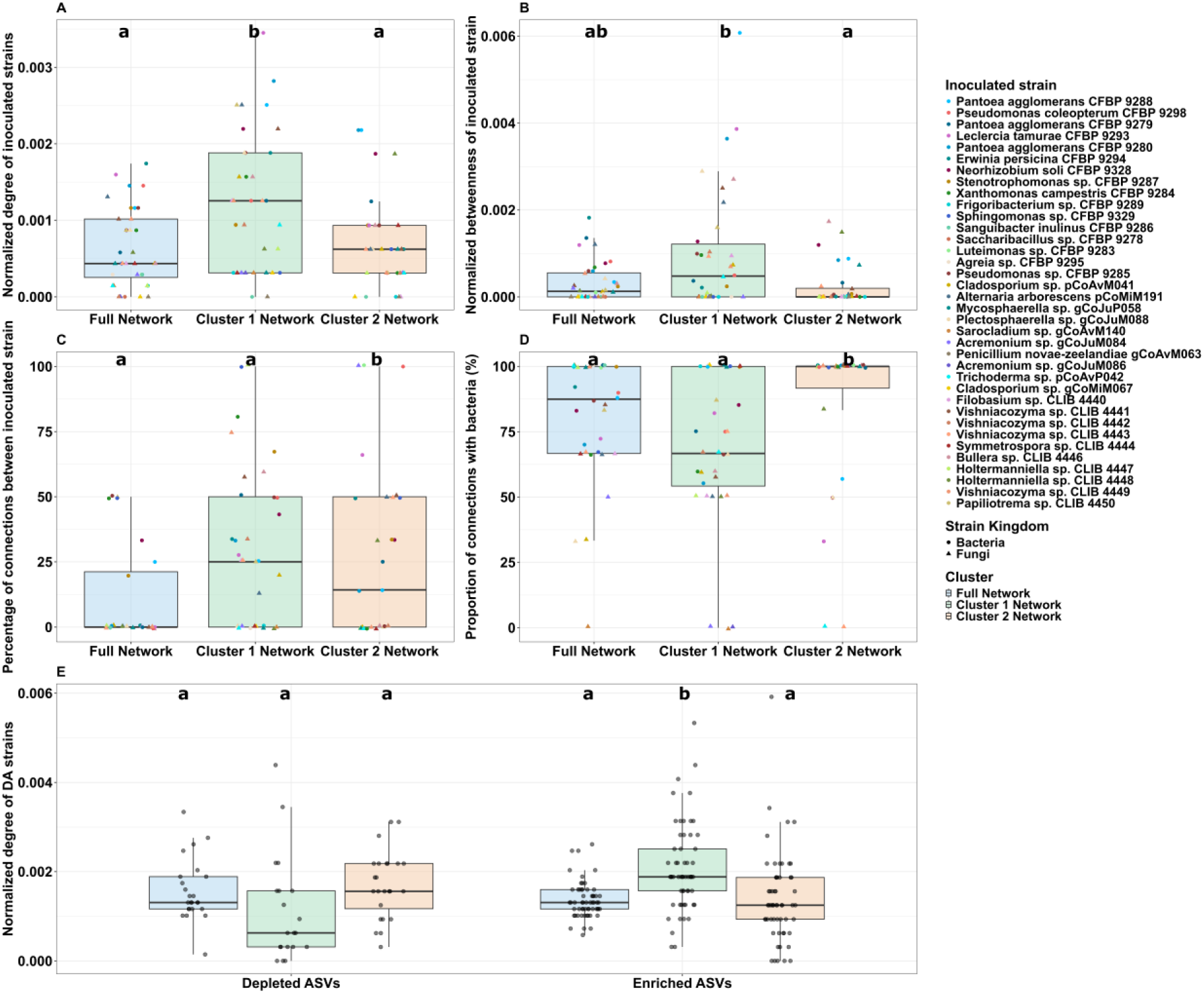
Network topological integration of introduced SynCom strains and differentially abundant native taxa. (A–B) Characterization of SynCom strain centrality metrics using (A) normalized degree centrality (Edges/(Nodes - 1)) and (B) normalized betweenness centrality across the full, Cluster 1, and Cluster 2 network frameworks. (C) Proportion (%) of assortative interactions, defined as the percentage of topological edges linking an inoculated SynCom strain directly to another introduced strain. (D) Proportion of connections of introduced strains with bacterial ASVs. (E) Normalized degree centrality of responding native taxa, partitioned by their differential abundance profiles (depleted vs. enriched ASVs) across the three network configurations. Different lowercase letters denote statistically significant differences among network types within each abundance group (Dunn’s test with Benjamini-Hochberg correction, p < 0.05).

Since SynComs strains integration modified the abundance of specific taxa, we investigated how the 82 identified differentially abundant (DA) native bacterial ASVs were behaving in these networks. These taxa were previously classified as enriched (n = 58) or depleted (n = 24, Fig. 6C). Within the global network, all 82 DA bacterial ASVs were exclusively located within the giant component. Within the cluster-specific networks, 77 DA ASVs were detected in Cluster 1, and 81 in Cluster 2, with the majority (79.30%, n = 65 ASVs) shared between the giant components of all three networks. The normalized degree (Fig. 8E) of Cluster 1 network DA strains (2.08 × 10^-3^) was higher than in the Cluster 2 (1.39 × 10^-3^) and global (1.36 × 10^-3^) networks (Dunn’s test with BH correction, p < 0.05). Conversely, depleted ASVs showed reduced connectivity in Cluster 1 (1.13 × 10^-3^) relative to Cluster 2 (1.70 × 10^-3^; Dunn’s test with BH correction, p = 0.062), while remaining comparable to the global network (1.57×10^-3^). To test whether these connectivity shifts were driven by interactions with inoculated SynCom members, we quantified the proportion of edges linking SynCom strain to DA strains. This proportion was consistent across all three networks, with no significant differences (Kruskal-Wallis test, p = 0.590), indicating that global connectivity changes were not directly attributable to SynCom-DA interactions. However, strain-level analysis revealed a notable exception: *Pantoea agglomerans* CFBP 9280. In the Cluster 2 network, 85.7% of its interactions (6 out of 7 edges) involved DA native taxa, evenly distributed between enriched and depleted ASVs. In contrast, in the Cluster 1 network, only 11.1% of its interactions (1 out of 9 edges) targeted DA taxa. Overall, DA bacterial strains displayed network-specific connectivity patterns, with increased connectivity correlated to enriched ASVs and decreased connectivity correlated with depleted ASVs. These modifications were not explained by overall interaction frequency with SynCom strains, although the distinct connections of *Pantoea agglomerans* CFBP 9280 suggests the presence of strain-specific interaction dynamics that may contribute to community restructuring.

The main characteristics of SynComs impact on seedlings are summarized in Fig. 9.

**Fig. 9:**
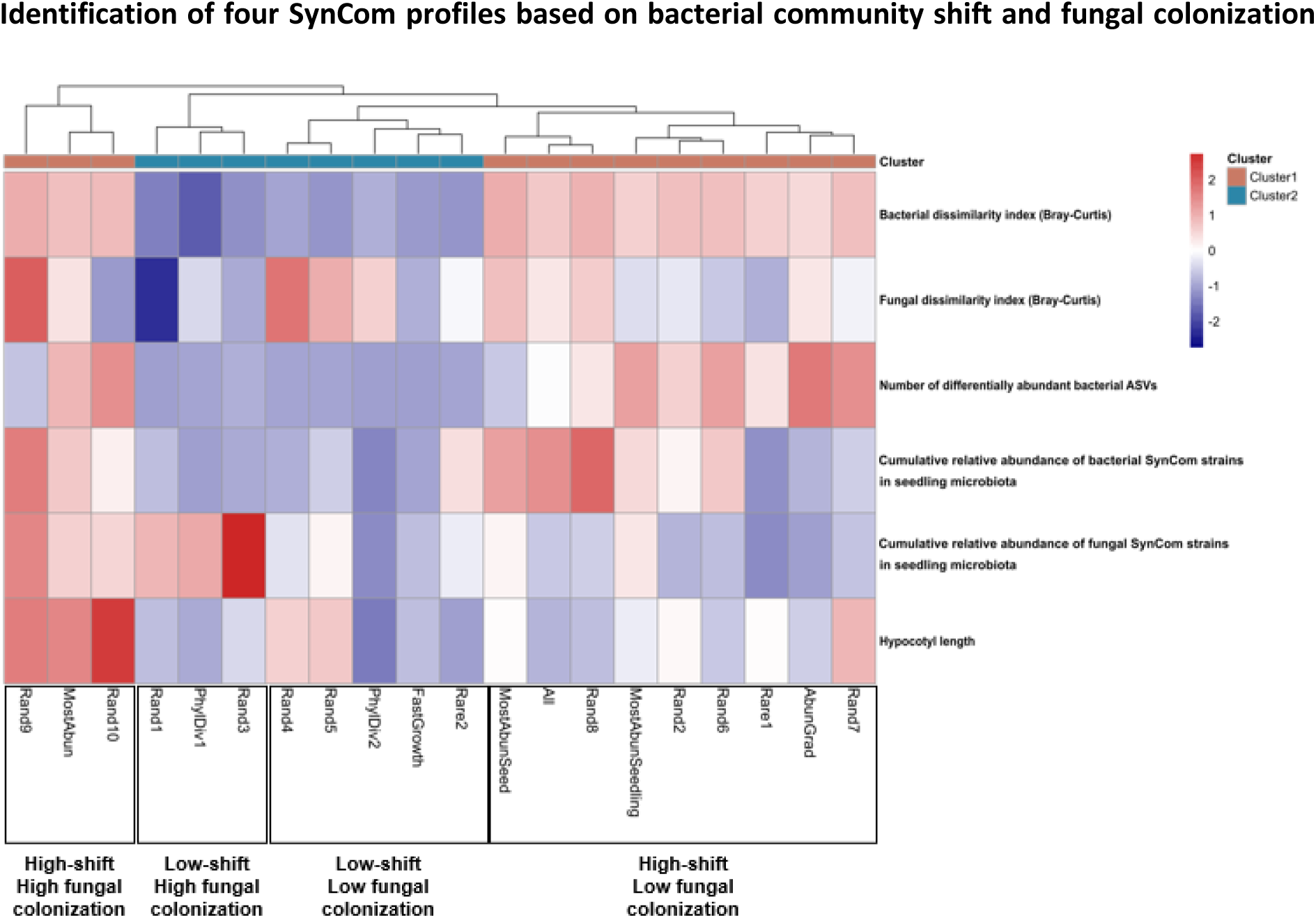
Main characteristics of SynComs’ impacts. Hierarchical clustering heatmap summarizing the SynCom effects on seedling microbiota and phenotype: Bacterial and fungal Bray-Curtis dissimilarity index compared to control condition, number of differentially abundant bacterial taxa, Cumulative relative abundance of bacterial and fungal SynCom strains and hypocotyl length. All displayed values are Z-score. SynComs belong to either Cluster 1 (Orange) or Cluster 2 (Blue) and are grouped in 4 distinct profiles.

While clustering primarily separated SynComs based on their ability to modulate seedling bacterial microbiota (Cluster 1: community shift, Cluster 2: no community shift), this more integrative approach revealed four distinct SynCom profiles defined by two criteria: bacterial community shift and fungal colonization. 3 SynComs (Rand10, Rand9, and MostAbun) combined bacterial community shift with high fungal colonization and were termed “High shift - High fungal colonization”. SynComs that modulated bacterial communities but had low fungal colonization were termed “High shift - Low fungal colonization.” The remaining SynComs with limited community impact were further split by colonization levels into “Low shift - High fungal colonization” and “Low shift - Low colonization”. The other criteria assessed (fungal community shift, bacterial colonization, hypocotyl length) did not clearly structure these profiles. Still, we can note that the “high shift - high colonization” SynComs displayed greater hypocotyl length than the other profiles. While comparing the four SynCom profiles with their composition (Fig. 1A) and compositional distance (Fig. 1B), no specific strain or SynCom composition matched with these profiles.

## Discussion

### Tailored seed inoculation enables successful seed microbiota modulation

In this study, we provide new insights on the transmission and impact of multiple multi-kingdom SynComs on the seedling microbiota of an important crop species. In line with our first hypothesis (H1), we successfully inoculated *Brassica napus* seeds with 20 distinct multi-kingdom SynComs and demonstrated their partial transmission to the seedlings. By comparing four inoculation methods, we identified alginate coating as a robust method for high microbial load inoculation and transmission of multi-kingdom SynComs. This optimization highlights the importance of tailoring inoculation strategies to the target plant model to achieve a sufficient propagule pressure, a key factor to enhance seedling colonization [15]. Such methodological considerations are particularly critical given the diversity of plant models used in SynCom studies [19], and the fact that relatively few studies directly monitor the fate of introduced communities [6]. With efficient seed inoculation, we show that bacterial communities are more easily modified than fungal ones, likely reflecting the higher resistance and stability of native fungal communities [58]. Despite successful inoculation, only a small fraction of inoculated strains persisted in 15-day old seedlings grown in non-sterile soil. Colonization levels were highly variable for both bacterial and fungal strains, with dominant seedling colonizers largely originating from the soil, in accordance with previous studies [29]. This contrasts with studies on younger seedlings of other plant (e.g., *Phaseolus vulgaris* [15]), and highlights the dynamic nature of microbiota assembly influenced by nutrients availability [59] and the exposure to oxidative stress [60] during early plant development.

### Evaluating SynCom design strategies: Key role of strain selection in colonization performance

We next tested whether SynCom design could influence colonization success (H2), specifically whether *a priori*-designed communities would outperform randomly assembled ones. Overall, only minor differences were observed between the two strategies, with random SynComs showing slightly higher fungal colonization. This suggests that community assembly rules alone are not sufficient to predict colonization outcome. Still, within the *a priori* SynComs, some designs outperformed others at colonizing seedlings. SynComs composed of naturally abundant taxa (*Cladosporium, Alternaria, Pantoea, Pseudomonas*) in the seed or seedling microbiota were the most effective for both bacterial and fungal colonization. This observation is consistent with previous studies [15] and likely reflects stronger ecological adaptation of these strains to plant-associated environments [25]. Importantly, these abundant taxa are not only frequently detected through sequencing but are also commonly isolated in culture collections [31], supporting their naturally high prevalence. In contrast, SynComs composed of rare taxa showed more variable outcomes. While some performed well, this was largely driven by the presence of specific strains such as *Pseudomonas* sp. CFBP 9285 and *Vishniacozyma* sp. CLIB 4449. Although rare taxa can naturally colonize seedlings [29, 62], our results suggest that their efficient transmission under artificial enrichment is strain-dependent and less predictable. A mixed SynCom combining taxa of different abundance levels created to mimic the natural community structure of natural seed microbiota [63], and reduce niche overlap [64] did not outperform communities composed solely of highly abundant taxa. This suggests that, in our system, ecological compatibility with the plant environment was a stronger determinant of establishment than niche complementarity. Similarly, SynComs designed to maximize phylogenetic diversity did not enhance colonization, indicating that reducing intra-community competition, or potentially increasing metabolic complementarity [65], is insufficient when strong competition with native microbiota persists [66]. The limited performance of SynComs composed of fast-growing strains further emphasizes that *in vitro* growth traits are poor predictors of *in planta* success, likely due to the mismatch between artificial media and plant-associated environments [67]. Finally, a highly diverse SynCom containing a large number of strains achieved strong colonization of seedling for both bacteria and fungi. This might reflect facilitative processes such as niche expansion [68], cross feeding [69] and even higher order interactions [23]. More broadly, our results suggest that native microbiota exert stronger competitive pressures than interactions within the SynCom itself. Taken together, these findings suggest that strain selection is a more critical determinant of SynCom performance than assembly strategy. Even randomly assembled communities can perform well, as long as they are drawn from a pool of ecologically relevant and well-adapted taxa. However, because randomly assembled SynComs were drawn from an ecologically relevant strain pool and overall colonization remained limited, the role of assembly criteria may be underestimated and likely depends on the environmental context of the inoculation and the desired functional outcomes.

### Identification of shared and kingdom-specific traits involved in seedling colonization

Monitoring individual strains across multiple SynCom contexts revealed strong strain-dependent colonization patterns, supporting our third hypothesis (H3). Strain identity emerged as the most explanatory variable for both bacterial and fungal colonization, indicating that colonization success relies on strain-specific traits that are not easily captured by simple phenotypic descriptors. This phenomenon may emerge from interaction-driven processes, in line with the concept of phenotypic niche extension, whereby microbial strains expand or shift through niche interactions with co-occurring taxa [68]. Nevertheless, several traits were associated with improved colonization of strains. For both bacteria and fungi, higher initial abundance on seed increased colonization of seedling. This relationship was linear for bacteria in accordance with previous research [15] whereas for fungi, it reached a plateau and then decreased, indicating an inoculation threshold beyond which colonization decreased. This threshold could be explained by autoregulatory mechanisms such as *quorum sensing* controlling conidia germination [70]. Lag phase also influenced colonization in both kingdoms, although with a modest and non-linear effect. A biphasic pattern emerged, where rapidly growing strains likely benefited from early niche occupation, whereas strains with longer lag phases appeared favored at later stages (around 21 h). This might be linked to the dynamic environmental modification during oilseed rape seed germination, including exudates released in the soil [71] which could create temporal niches favoring late-establishing strains, and oxidative stress release that induce strain selection [60, 72]. Additional kingdom-specific traits were identified. For bacteria, larger genome size was positively correlated to a better colonization. Since bacterial genomes are highly dense in genes [73], higher genome size could be linked to greater metabolic versatility and adaptive capacity [74]. For fungi, natural prevalence on seed was positively linked to better colonization, suggesting that dominant seed-associated fungi are better adapted to withstand germination-related stresses and persist during early plant development.

### Impact of SynCom inoculation on bacterial native communities revealing distinct profiles

We demonstrated that SynCom inoculation significantly altered the native bacterial microbiota, leading to the emergence of two distinct community profiles: weakly disturbed and strongly modified bacterial communities. In contrast, fungal microbiota remained stable. These two clusters were robustly confirmed by ASV differential abundance analyses, in line with current methodological frameworks [75, 76]. This clustering further enabled the construction of two distinct multi-kingdom co-occurrence networks, corresponding to SynCom-responsive (Cluster 1) and non-responsive (Cluster 2) communities. Differences in network topology between these clusters likely reflect the extent to which inoculated strains successfully integrate into the native microbiota and influence its assembly. A total of 82 native bacterial ASVs were identified as responsive to SynCom inoculation, most of which likely originated from the soil, indicating that SynComs influence community assembly by modulating the recruitment of environmental taxa. Importantly, these shifts were not attributable to any single SynCom composition, as the 14 SynComs associated with modified communities did not share a common set of strains. In responsive communities (Cluster 1), inoculated strains appeared more central and connected within the network, suggesting successful integration into the native microbiota and a greater capacity to reshape microbial interactions, including the recruitment of additional taxa. In contrast, non-responsive communities (Cluster 2) displayed network structures closer to the native state, indicating limited integration of introduced strains. By separating these two community response types, our approach allowed a more accurate identification of SynCom-driven dynamics and highlighted the context dependency of microbial establishment and interaction within plant-associated microbiota. At the strain level, *Pantoea agglomerans* CFBP 9280 displayed a different co-occurrence profile between clusters. This strain’s relative abundance being highly enhanced in seedling of inoculated conditions, its introduction might play a pivotal role in early seedling microbial assembly through interactions with environmental taxa, as it belongs to a seed core microbiota genus with a potential high transmission in a natural context [77]. Interestingly, samples with altered bacterial communities exhibited increased interkingdom connectivity, highlighting the importance of bacteria-fungi interactions in shaping microbiota assembly [20].

Finally, four different SynCom profiles emerged from their impact on seedling, whether they shifted bacterial communities and/or fungal strains colonized the seedlings. The modulation of bacterial communities without or with low colonization of SynComs translates to a potential effect of transient interactions, a pattern already observed in soil only between bacterial inoculant and communities [78]. On the other hand, a combined fungal colonization with bacterial shift can be associated with longer-lasting inter-kingdom interactions between inoculated and native strains mediated by the installation of introduced fungi. Notably, this profile of high colonization and community shift was associated with an increase of hypocotyl length, hinting at specific design that might impact the plant more efficiently [79]. Interestingly, profiles with no bacterial community shift were observed whether SynCom colonized or not seedlings, showing that SynCom transmission is not necessarily associated with broad community modification. Overall, these results highlight the complexity of microbiota modulation, which arises from multiple interactions within and between kingdoms, between inoculated strains, native microbiota, and the plant. Since our observations are limited to a 15-day time point, the potential role of transient or low abundance taxa remains to be investigated, and may exert a lasting effect on community assembly [80].

## Conclusions

Through the inoculation and monitoring of 20 different multi-kingdom SynComs on Brassica napus seeds, we showed that inoculated strains were partially transmitted to seedlings while 14 SynComs significantly reshaped the seedling bacterial microbiota by modifying the recruitment of 82 bacterial taxa. Our design strategy highlighted that strain identity mattered more than the SynCom assembly strategy for seedling colonization. We also demonstrated that the fungal microbiota was more resistant to both seed colonization and community modulation in contrast to bacterial communities. These modulations of the bacterial communities were explained by a better integration of microbial strains in the network community structures. Based on our findings, we distinguish four different SynComs profiles: SynComs that have high colonization and those that do not, as well as between communities that disrupt or preserve seedling bacterial recruitment. These non-disruptive profiles deserve attention given growing concerns about the ecological risks of microbial inoculants [8] by their potential effect on plant health while having reduced impact on native communities. Altogether, these findings provide actionable directions for improving SynCom design, suggesting that leveraging ecological processes such as host adaptation, optimal inoculation density, and network integration could enhance both colonization efficiency and plant phenotypic outcomes.

## Declarations

### Ethics approval and consent to participate

Not applicable

### Consent for publication

Not applicable

### Availability of data and materials

The datasets generated and analyzed during the current study are available in the INRAE forge repository, https://forge.inrae.fr/irhs-emersys/2026_logan_inoseed

### Authors’ contributions

LS produced the data, analyzed, interpreted and was the main contributor in writing the manuscript. CC, NG and MS provided support for experimentation, insight for analysis as well as thorough corrections and rephrasing of the manuscripts. CM helped for bacterial culture and phenotyping as well as for amplicon bank preparation and sequencing. MB assembled bacterial genomes. AH and KH helped with plant culture. KM provided help and expertise for the co-occurrence network analysis, power calculation as well as corrections and rephrasing of the manuscript. MM provided support for fungal strain culture, phenotyping and taxonomic affiliation. All authors read and approved the final manuscript.

## Acknowledgements

We thank the platform PHENOTIC [81] for the plant culture, the ANAN platform for genome and amplicon sequencing, the Himic platform for the individual growth culture assay. We also thank the CIRM-CFBP [82] and CIRM-Levures for the strain conservation. We thank Elise Alix and Bernard Moulin (IGEPP) and the Experimental Unit UE La Motte for the oilseed rape seed production in the field and the CRB BrACYSol for providing the oilseed rape genetic resources. We would also like to thank Alain Sarniguet for his advice and expertise on *B. napus* seed experiments, Justin Colou for her knowledge on fungal genome assembly, Martial Briand for his expertise on bacterial genomes and Philippe Simoneau for his insight on fungal taxonomy. A special thanks for everyone who helped inoculating, sowing and harvesting the seed and seedlings: Louna Colaert-Sentenac, Thomas Chadelaud, Coralie Marais, Clara Centa, Marie-Agnès Jacques, Chrystelle Brin, Martial Briand, Nathan Kavunu, Géraldine Taghouti, Oscar Joubert and Franck Bastide.

## Additional files

**Supplementary Fig. 1:**
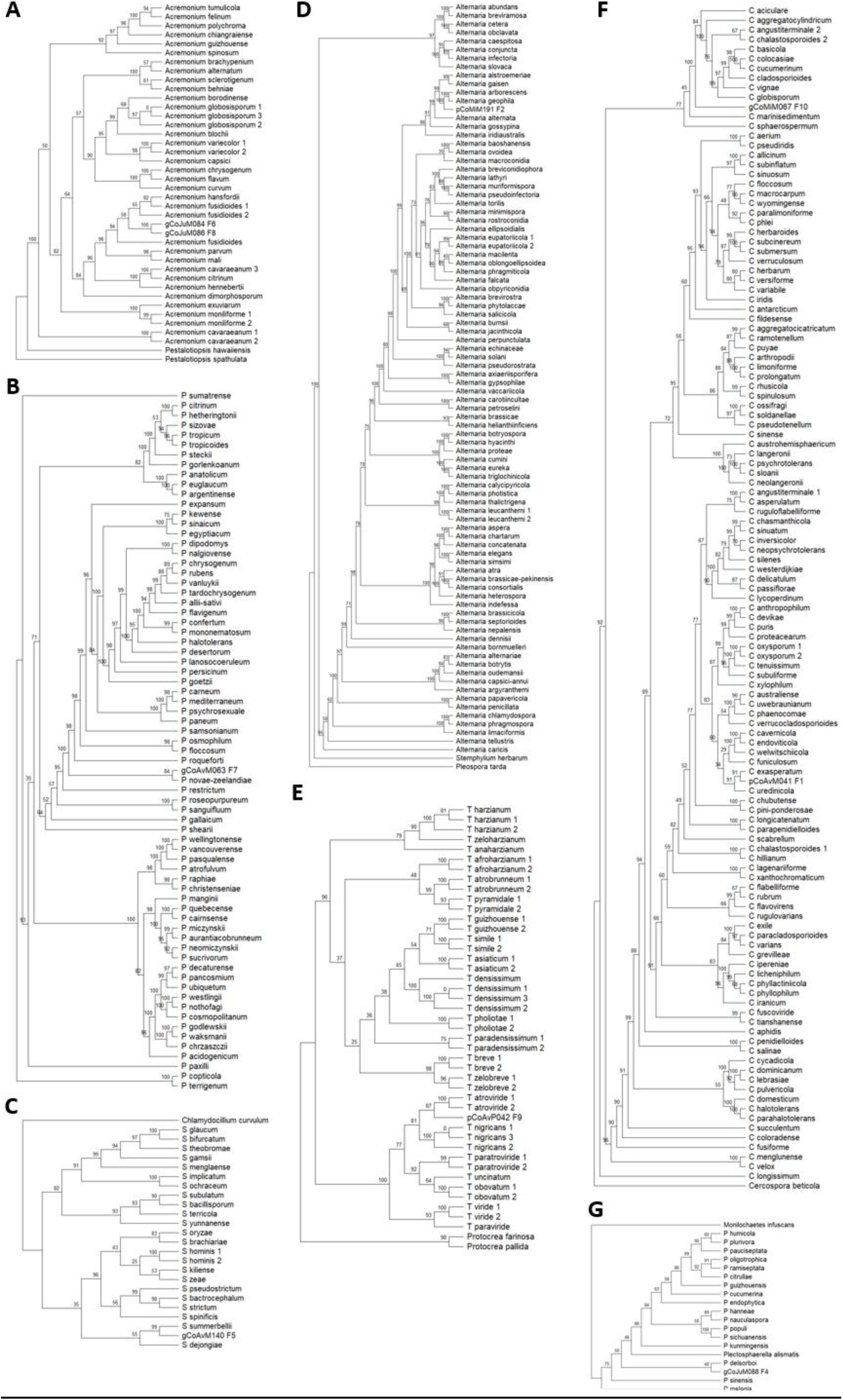
Phylogenetic trees of the genus (A) *Acremonium*, (B) *Penicillium*, (C) *Sarocladium*, (D) *Alternaria*, (E) *Trichoderma*, (F) *Cladosporium* and (G) *Plectosphaerella* using the IQ-TREE Ultrafast bootstrap (1000 iterations)

**Supplementary Fig. 2:**
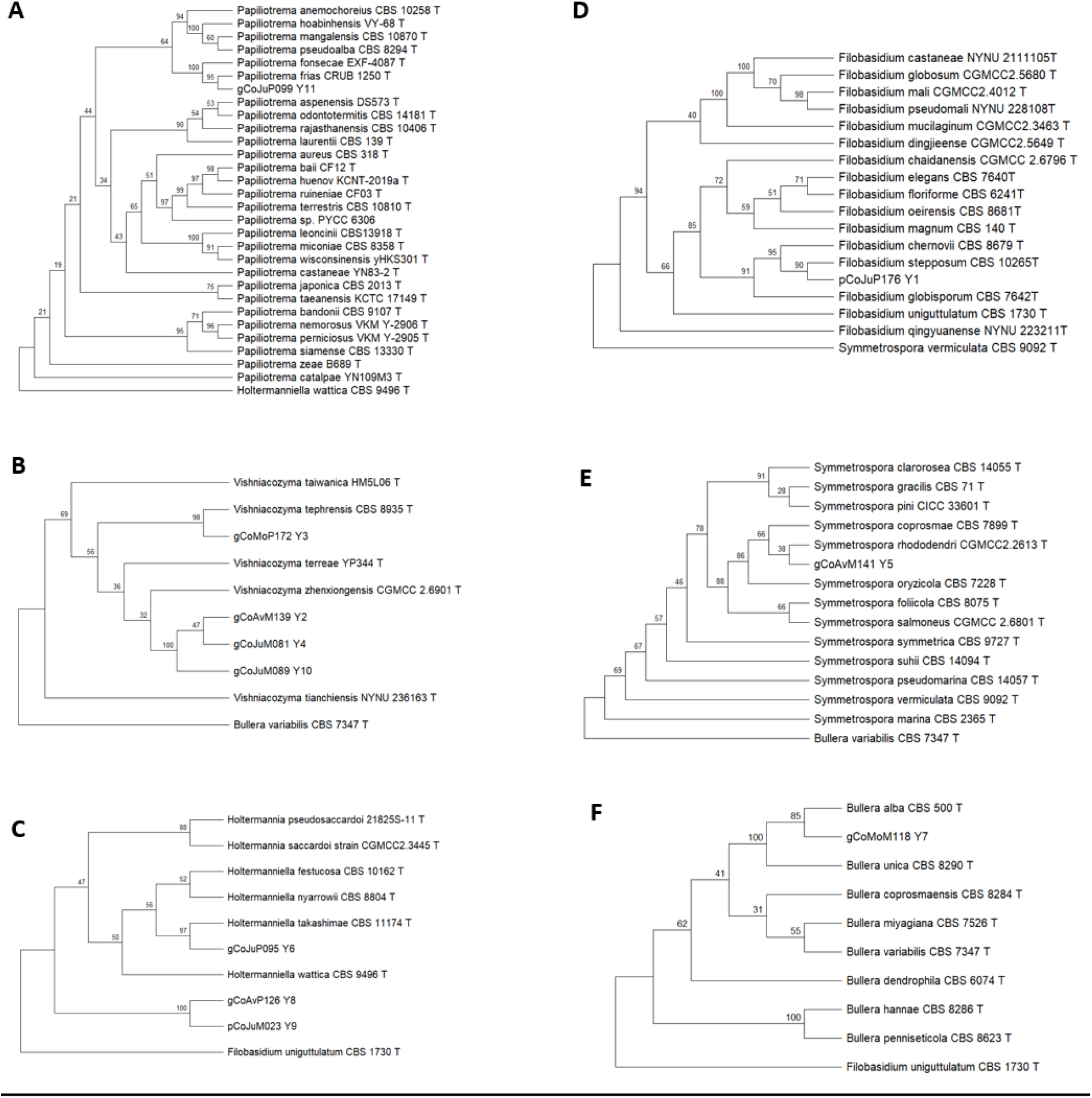
Phylogenetic tree of the genus (A) *Papiliotrema*, (B) *Vishniacozyma*, (C) *Holtermaniella*, (D) *Filobasidium*, (E) *Symmetrospora* and (F) *Bullera* using the IQ-TREE Ultrafast bootstrap (1000 iterations)

**Supplementary Fig. 3:**
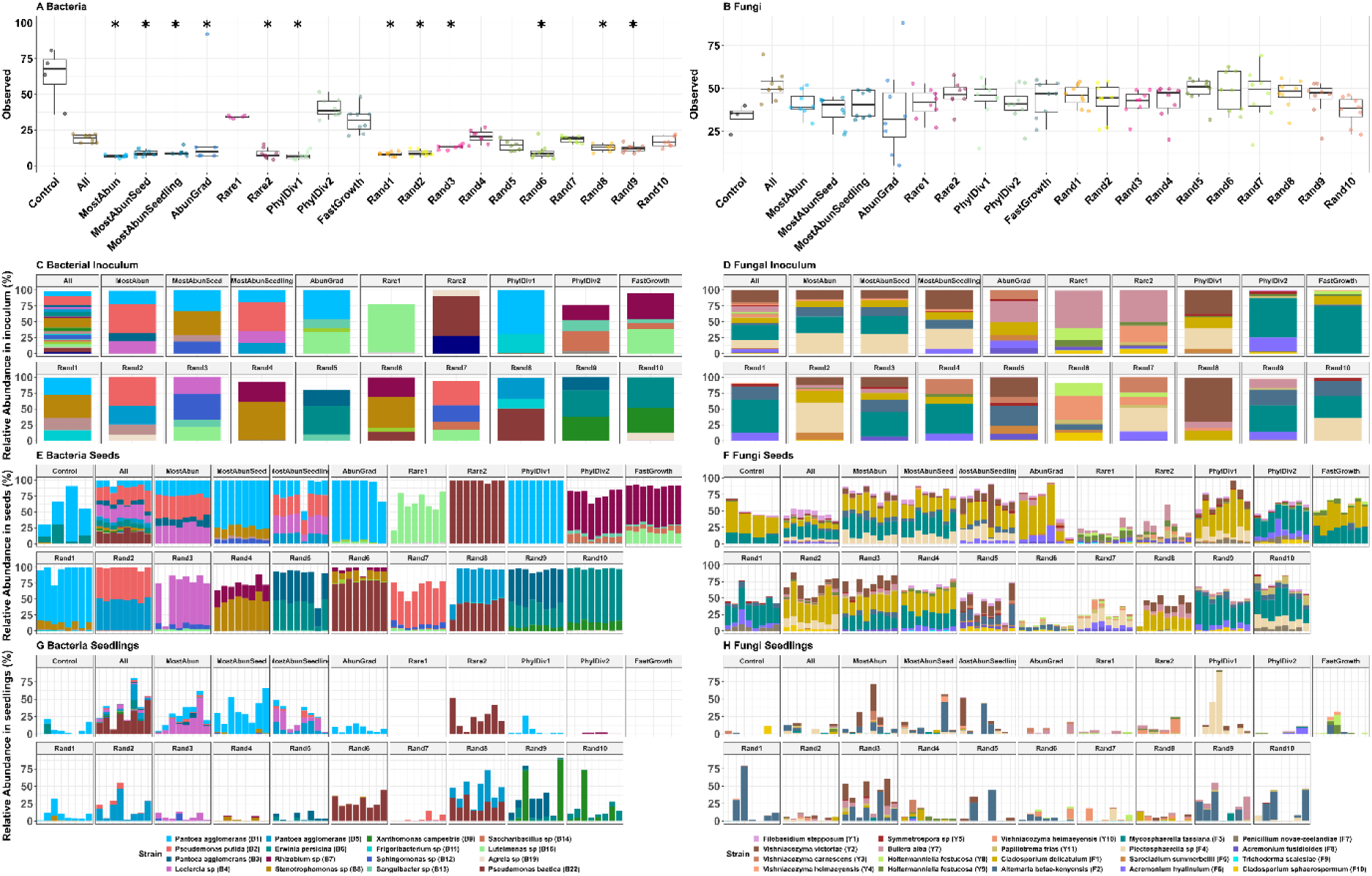
Assessment of microbial alpha diversity and taxonomic composition across seeds and seedlings. (A–B) Observed alpha richness (number of unique ASVs) for (A) bacterial and (B) fungal communities. For the non-inoculated control condition, each replicate point represents a pooled batch of 20 seeds, whereas all other points represent individual inoculated seeds. Asterisks denote significant differences relative to the control group (Pairwise comparisons using Dunn’s many-to-one test with Benjamini-Hochberg correction, p < 0.05). (C–H) Taxonomic profiling of recovered inoculated strains across different experimental stages for bacterial (left panels: C, E, G) and fungal (right panels: D, F, H) communities. Panels represent relative abundances within the initial inoculum (C, D), the mature seed microbiota (E, F), and the emerged seedling microbiota (H, G).

**Supplementary Fig. 4:**
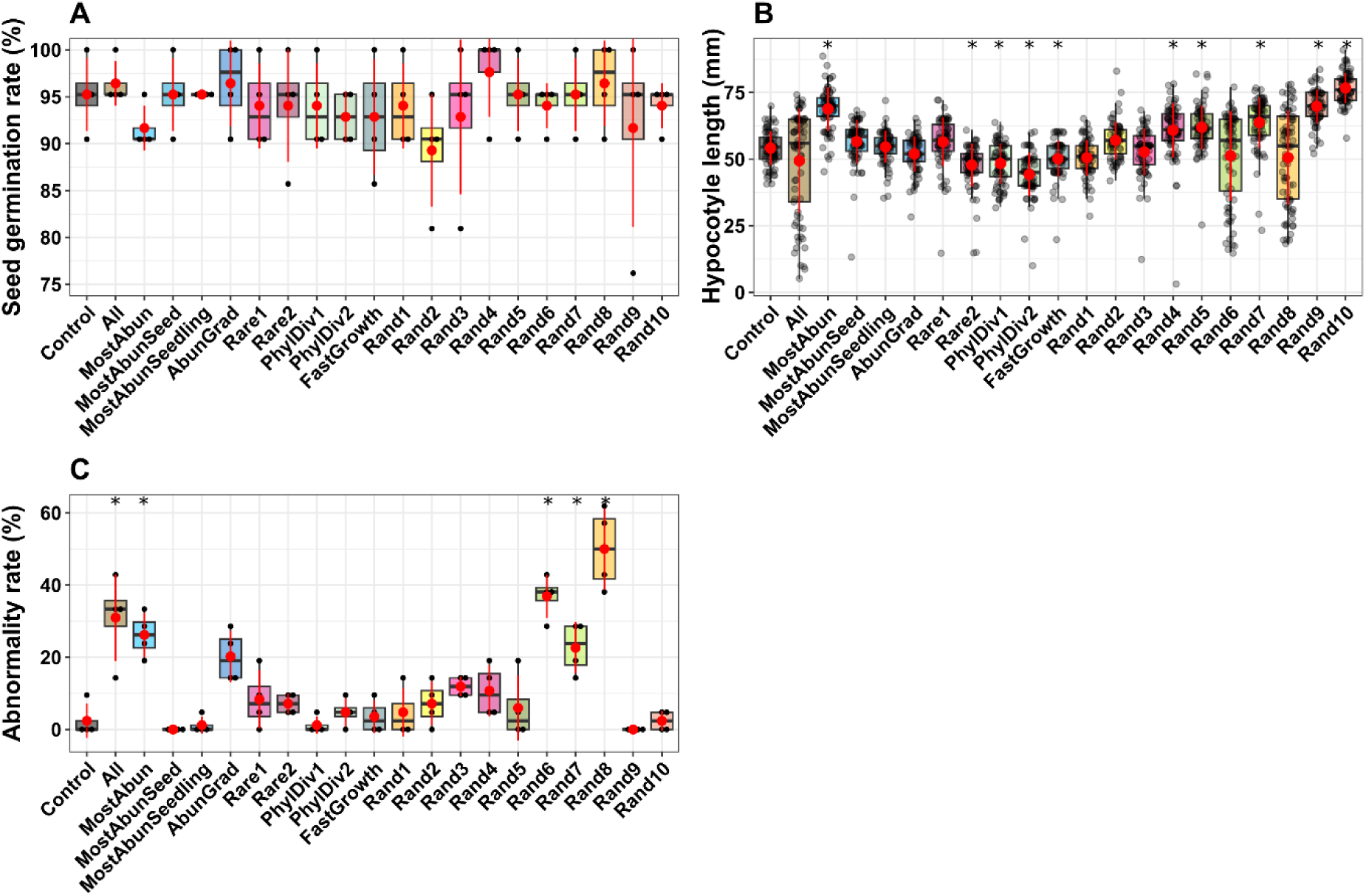
Phenotypic assessment of *Brassica napus* seedlings 15 days post-inoculation with various SynCom formulations. (A) Germination percentages, (B) hypocotyl length, and (C) proportion of morphologically normal seedlings. Asterisks (*) denote statistically significant differences relative to the non-inoculated control group (Pairwise comparisons using Dunn’s many-to-one test with Benjamini-Hochberg correction, p < 0.05).

**Supplementary Fig. 5:**
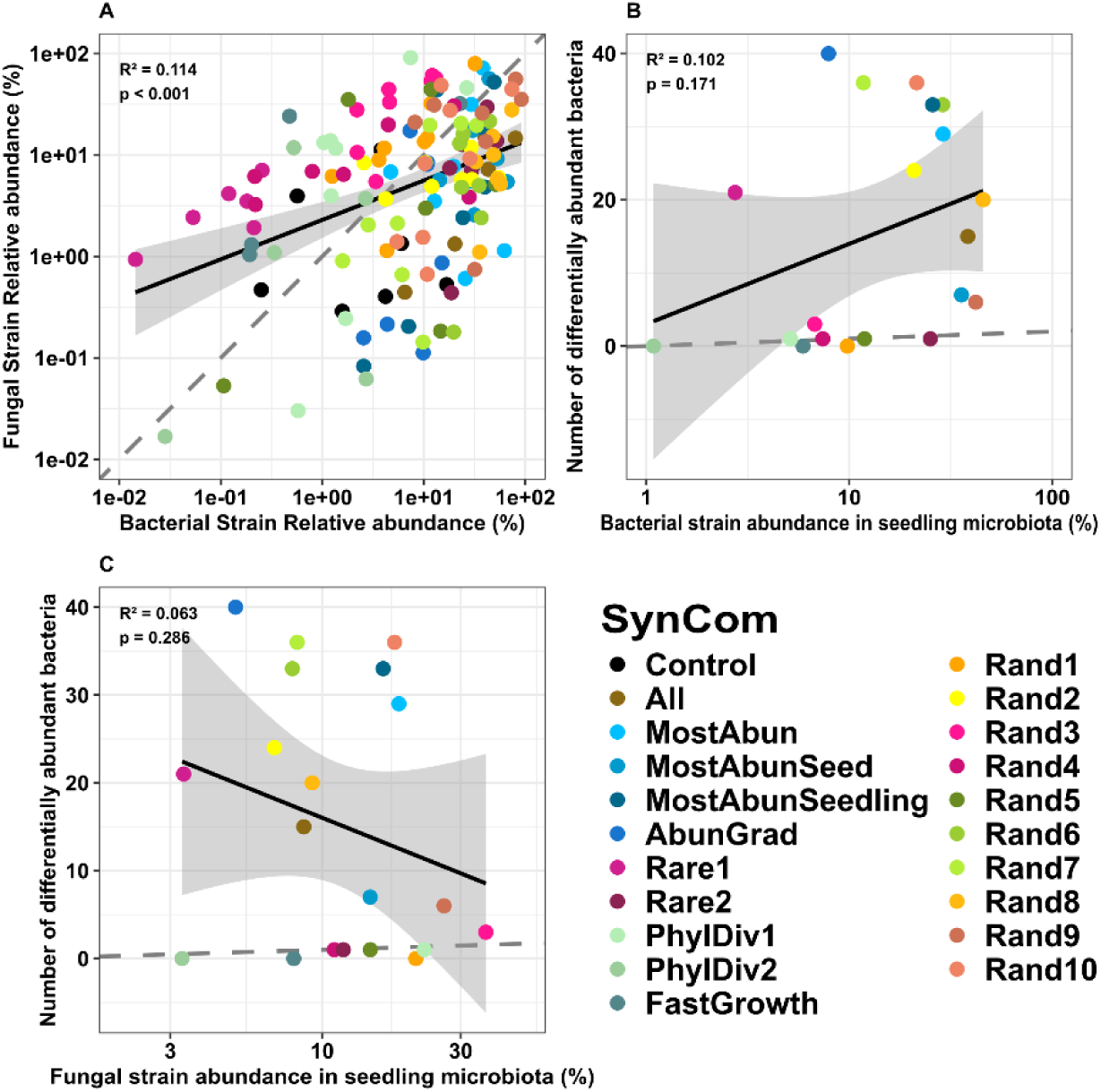
Correlation analyses between colonization dynamics and differential taxonomic abundance in seedlings. (A) Linear regression evaluating the relationship between the cumulative relative abundance of fungal and bacterial SynCom strains within individual seedlings. (B–C) Relationship between the number of differentially abundant host taxa and the cumulative abundance of introduced (B) bacterial or (C) fungal SynCom strains within the seedling matrix.

**Supplementary Fig. 6:**
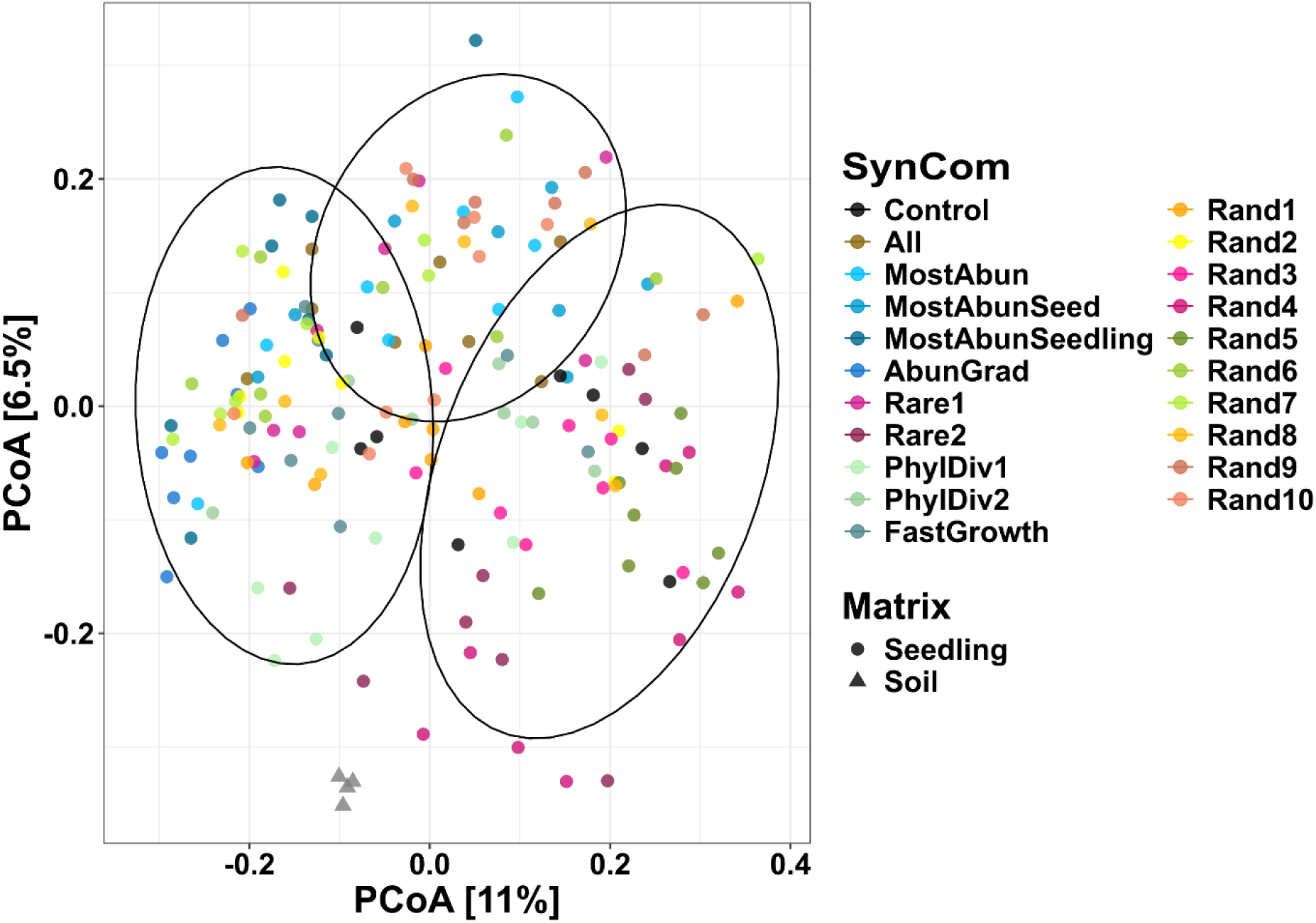
Principal Coordinate Analysis (PCoA) based on Bray-Curtis dissimilarity matrices (log(x+1) transformed) for fungal communities across bulk soil, non-inoculated control seedlings, and SynCom-inoculated seedlings. Cluster overlays are derived from k-means clustering partitioning using the NbClust package in R based on Euclidean distances.

**Supplementary Fig. 7:**
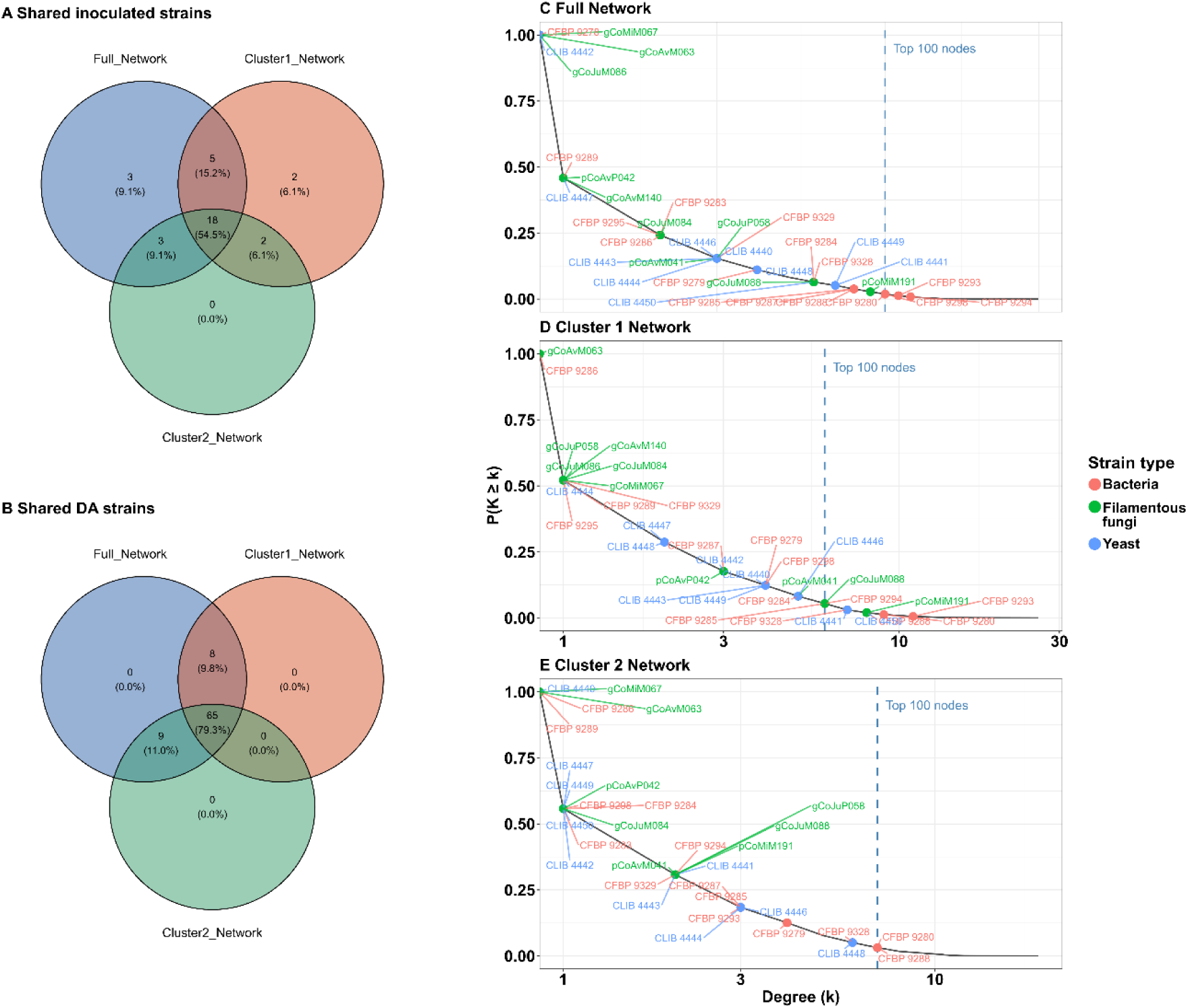
(A–B) Venn diagrams illustrating the overlapping and unique (A) introduced SynCom strains and (B) differentially abundant bacterial ASVs localized within the giant components of the Full, Cluster 1 and Cluster 2 networks. (C–E) Complementary cumulative distribution function (cCDF) curves modeling the topological degree distributions within the (C) full, (D) Cluster 1 and (E) Cluster 2 network models. Individual introduced SynCom strains are highlighted by taxonomic groups: bacteria (green points), filamentous fungi (red points), and yeasts (blue points). The vertical black dashed line marks the threshold designating the top 100 most highly connected nodes within each respective network architecture.

**Table S1:**
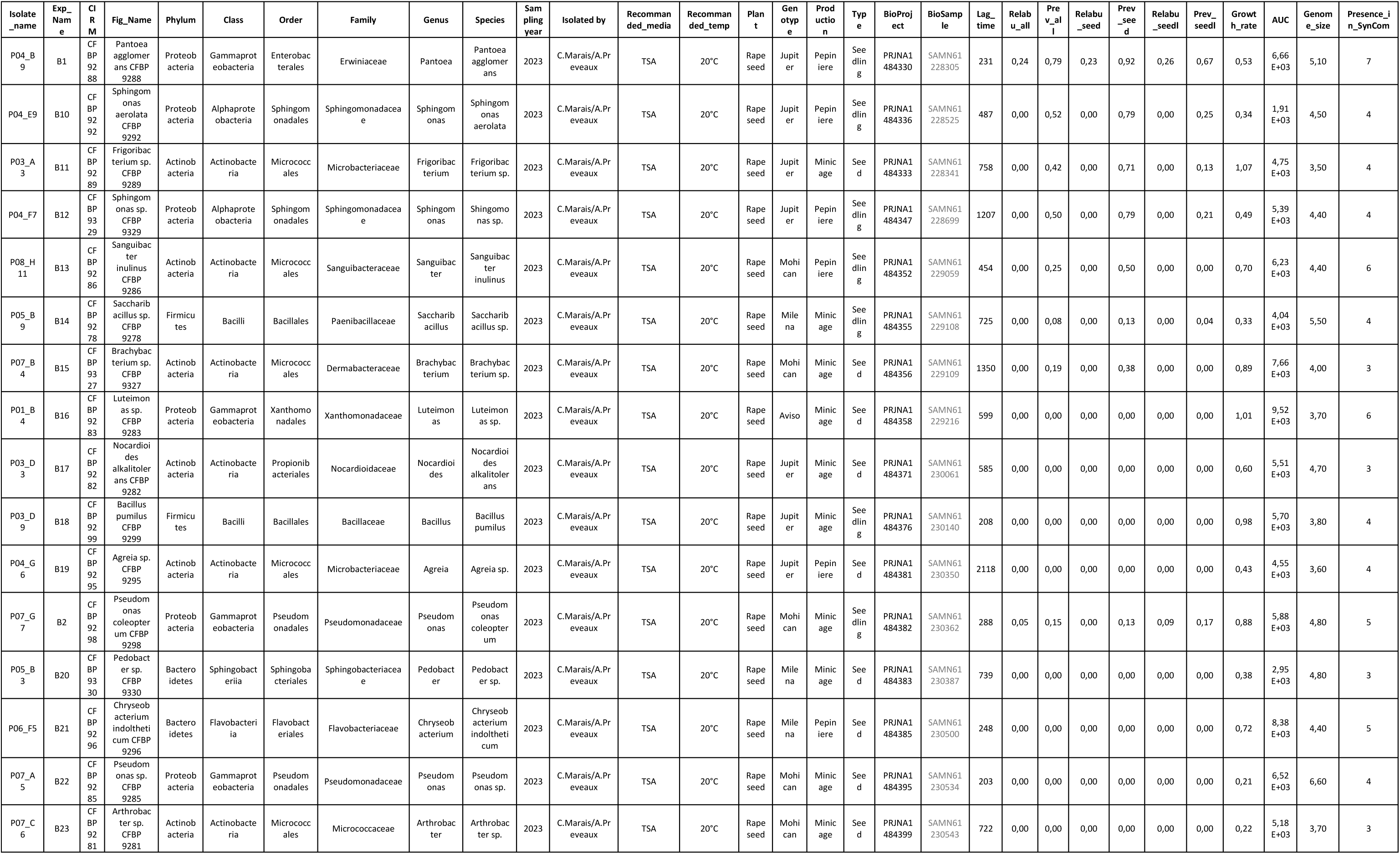
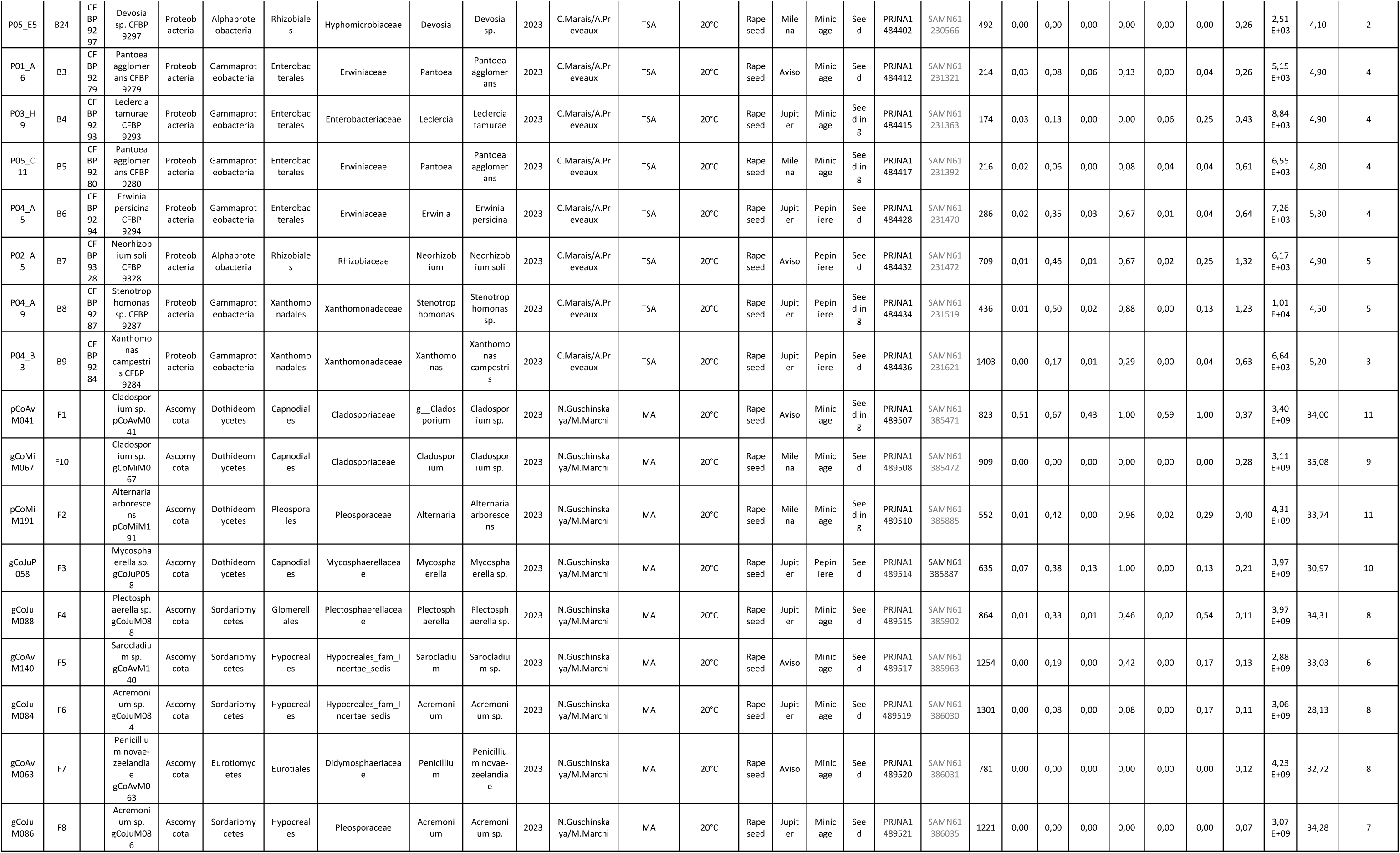
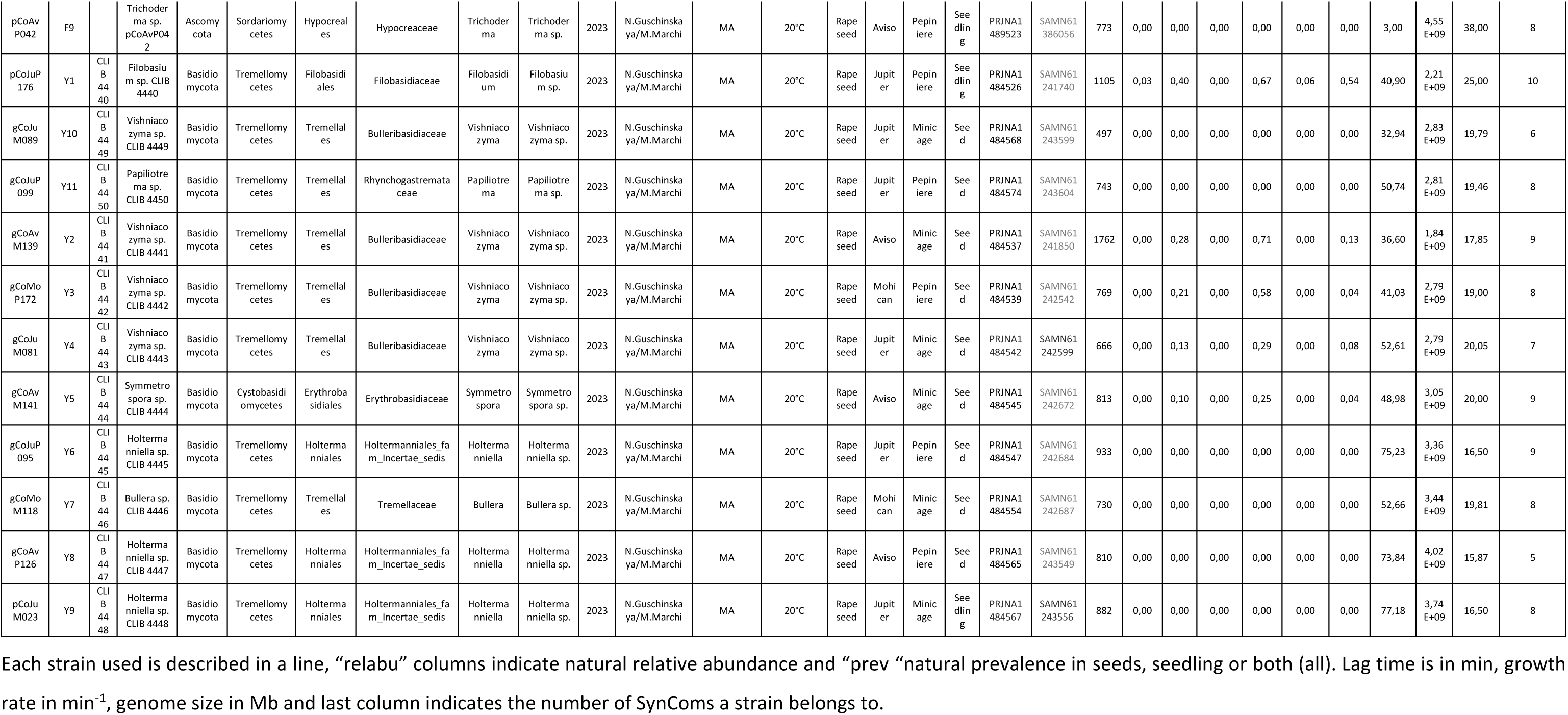
Strain used in SynCom inoculation and their information.

**Table S2:**
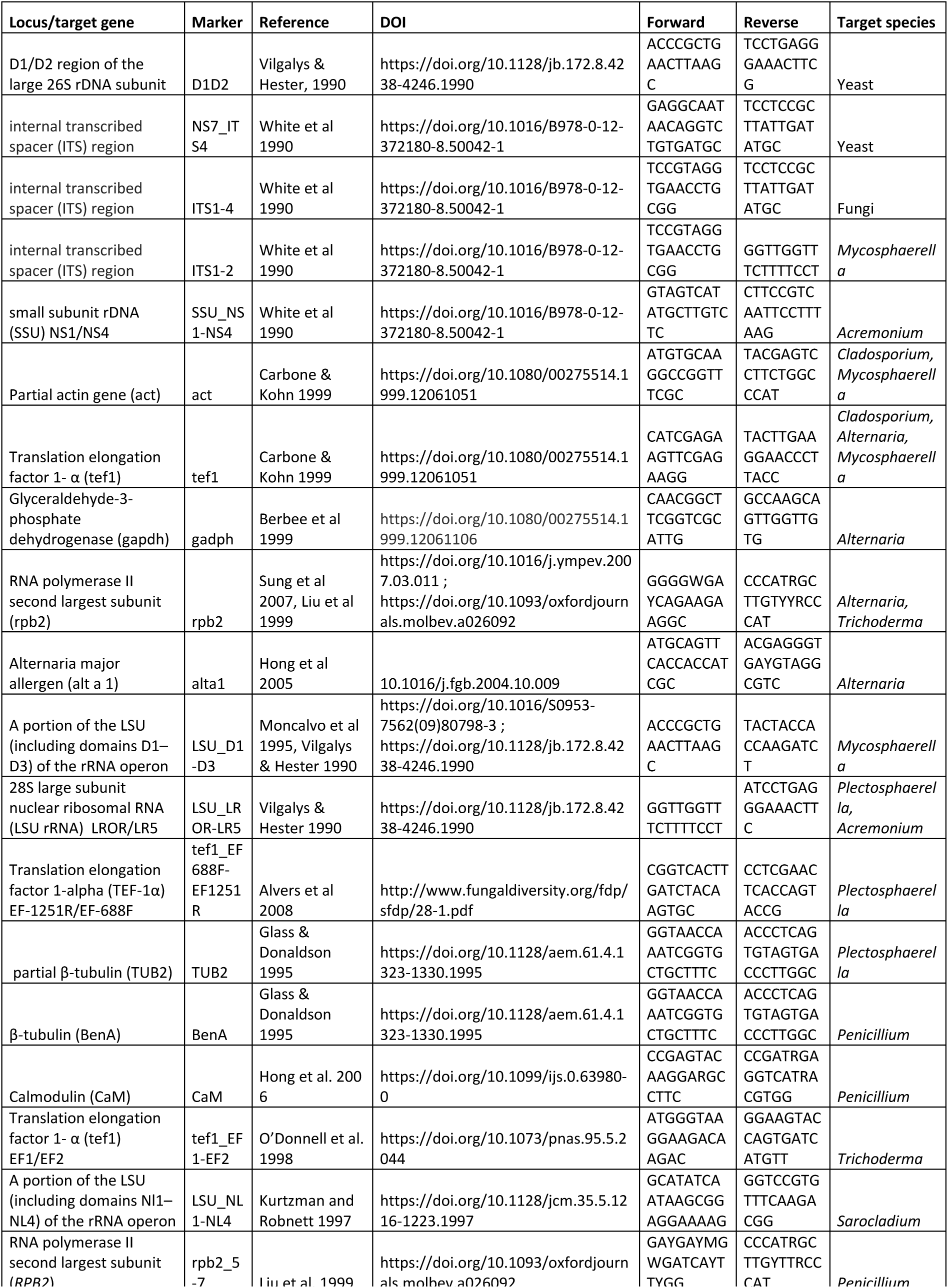
Loci used for taxonomic identification of fungal strains.

**Table S3:**
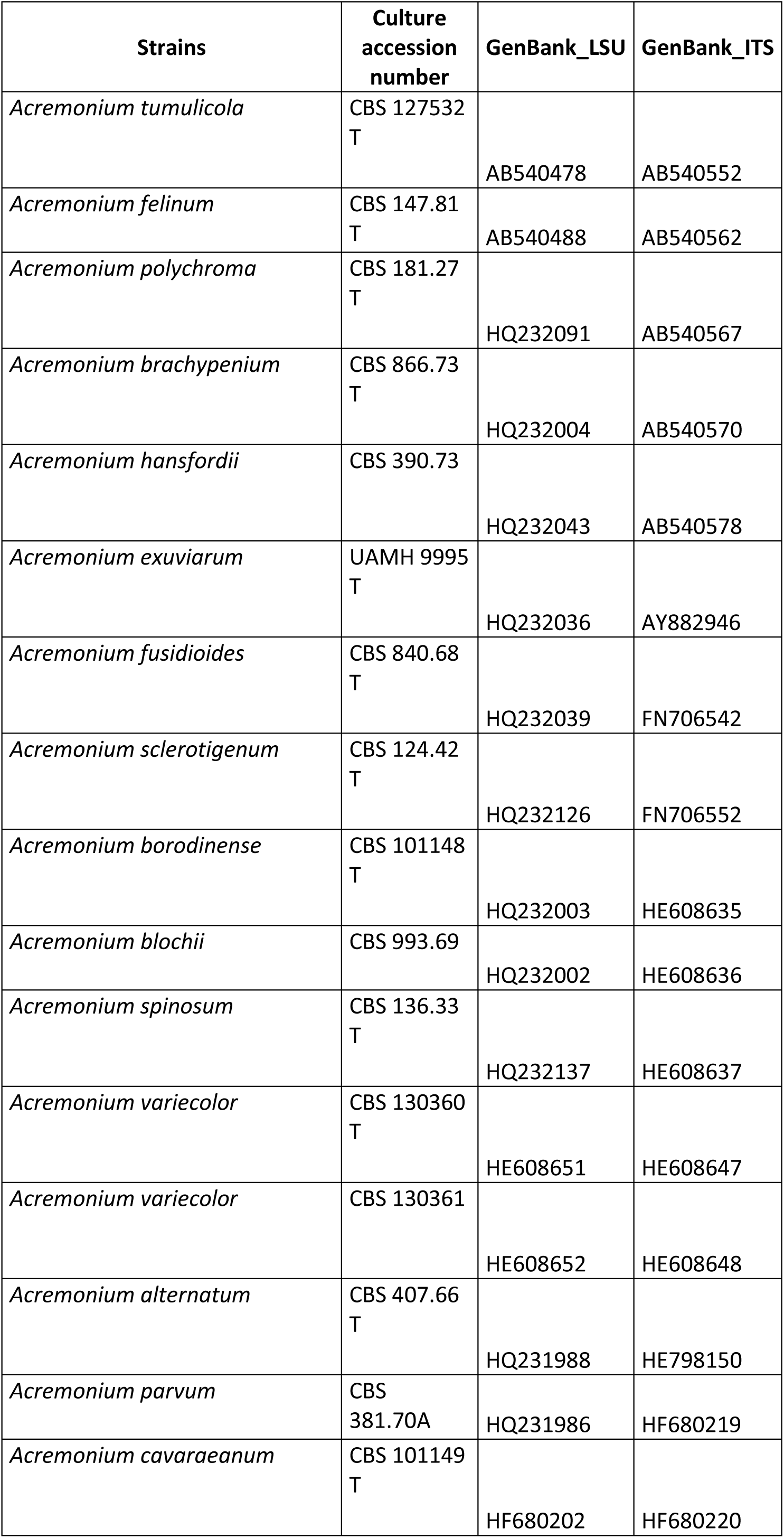
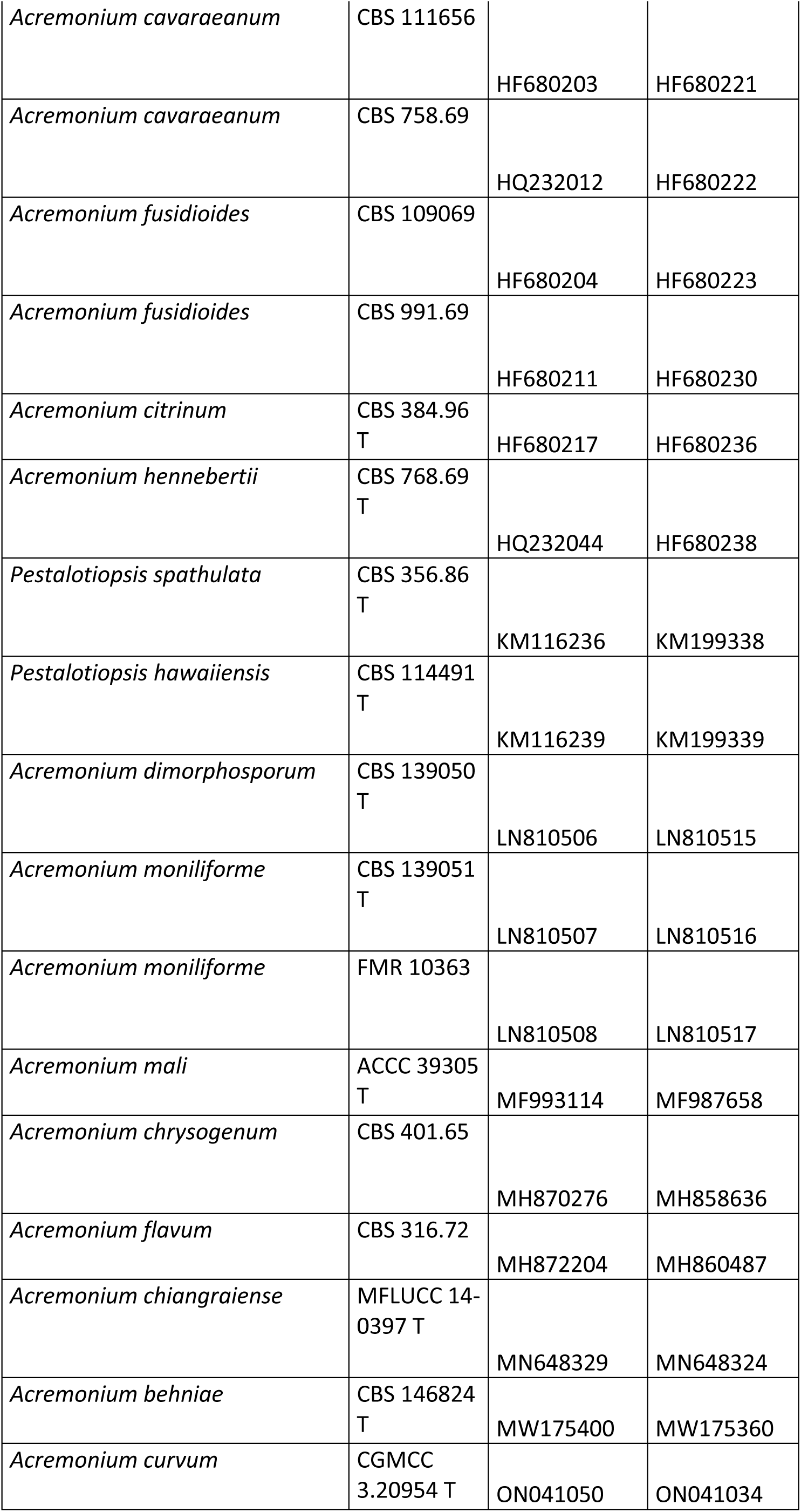
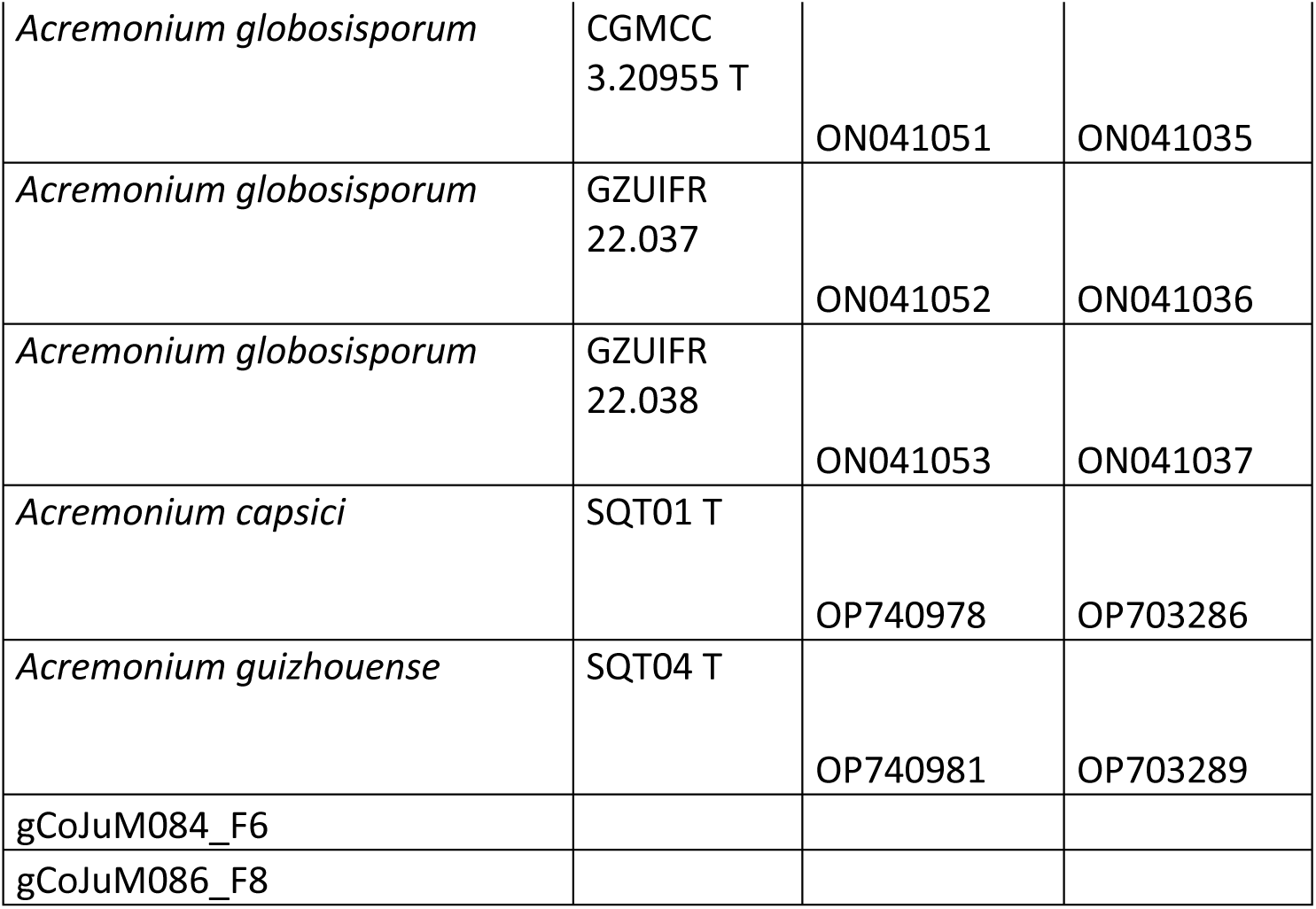
Strains used for the taxonomic identification within the *Acremonium* genus.

**Table S4:**
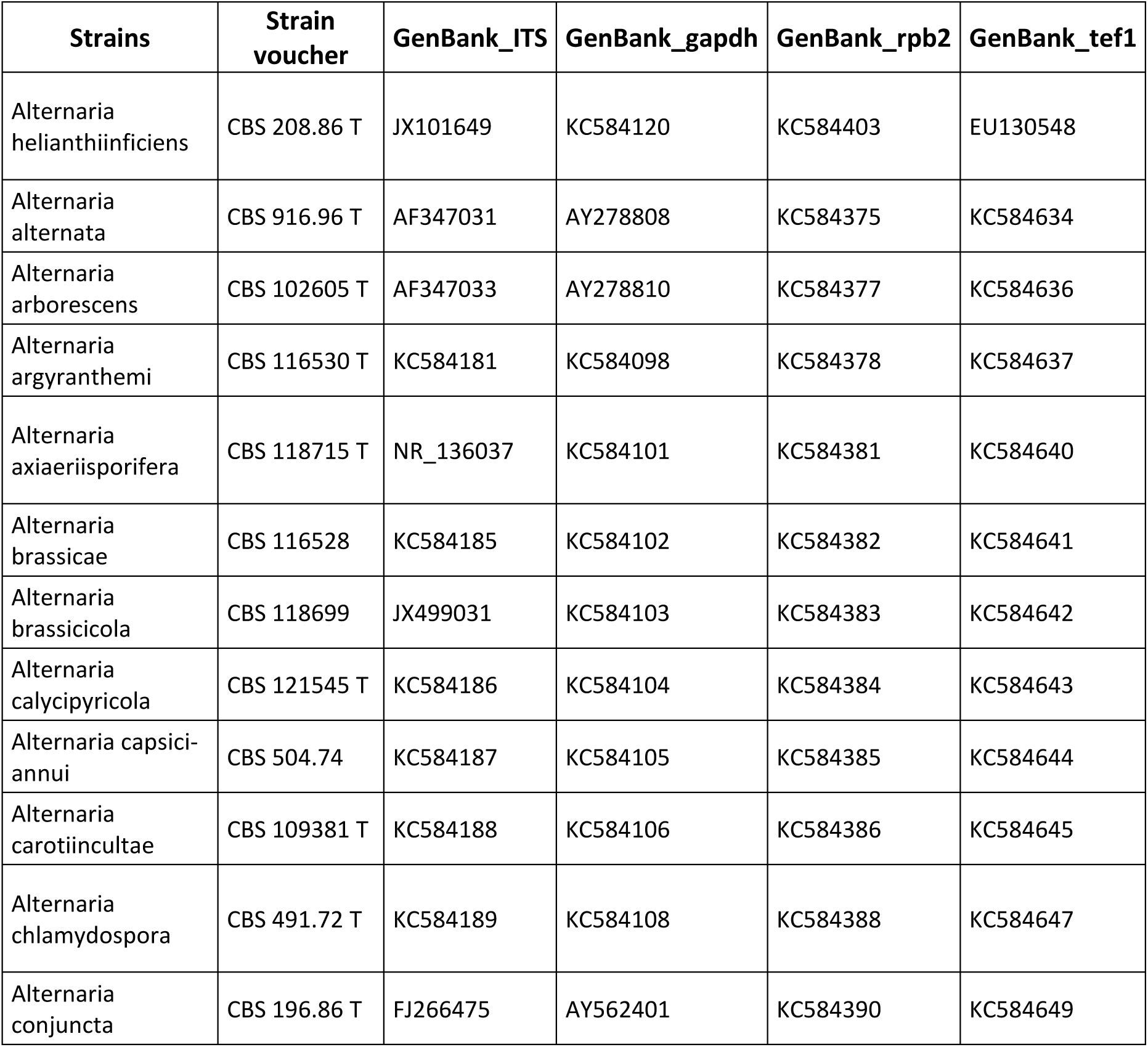
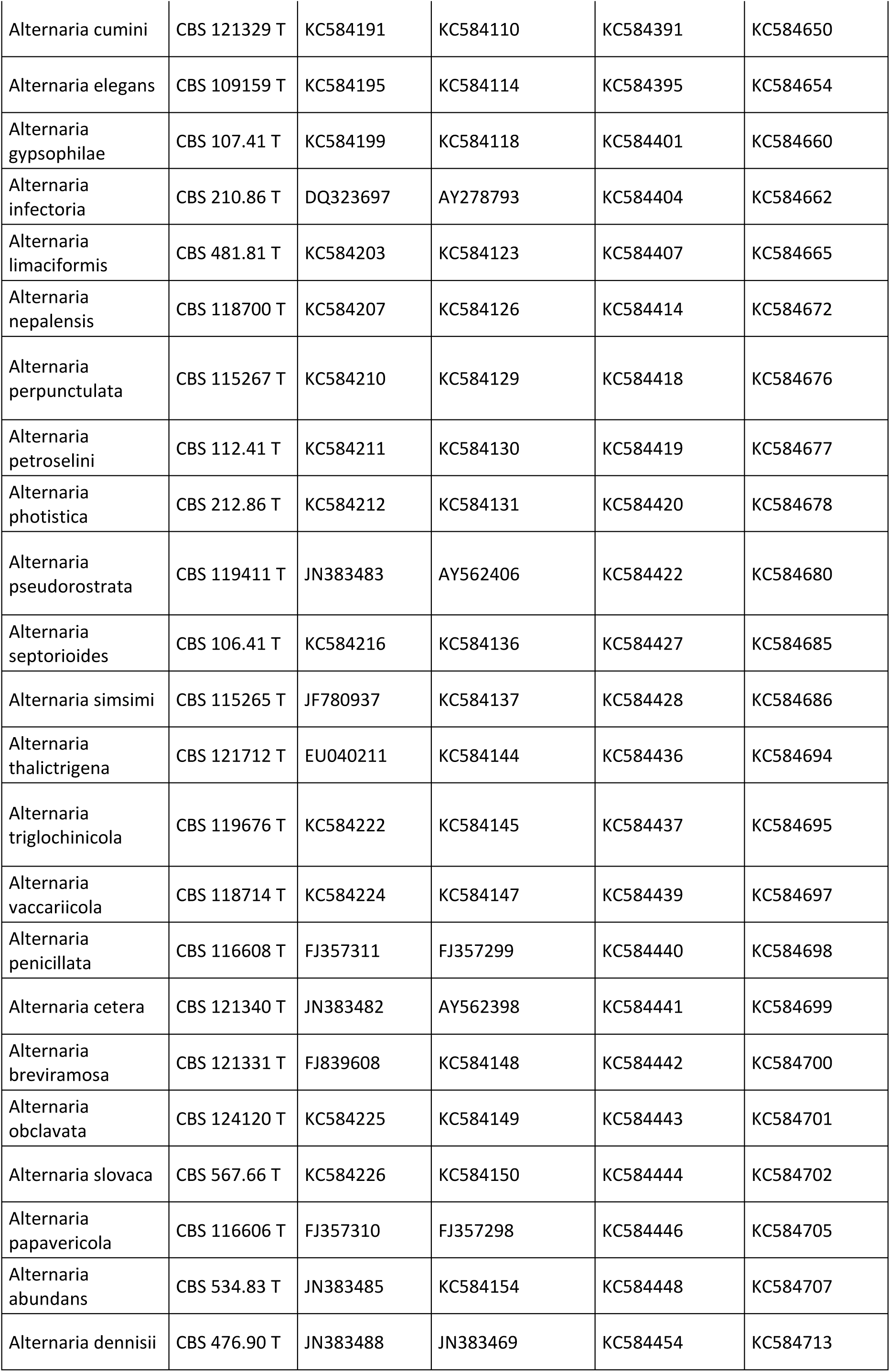
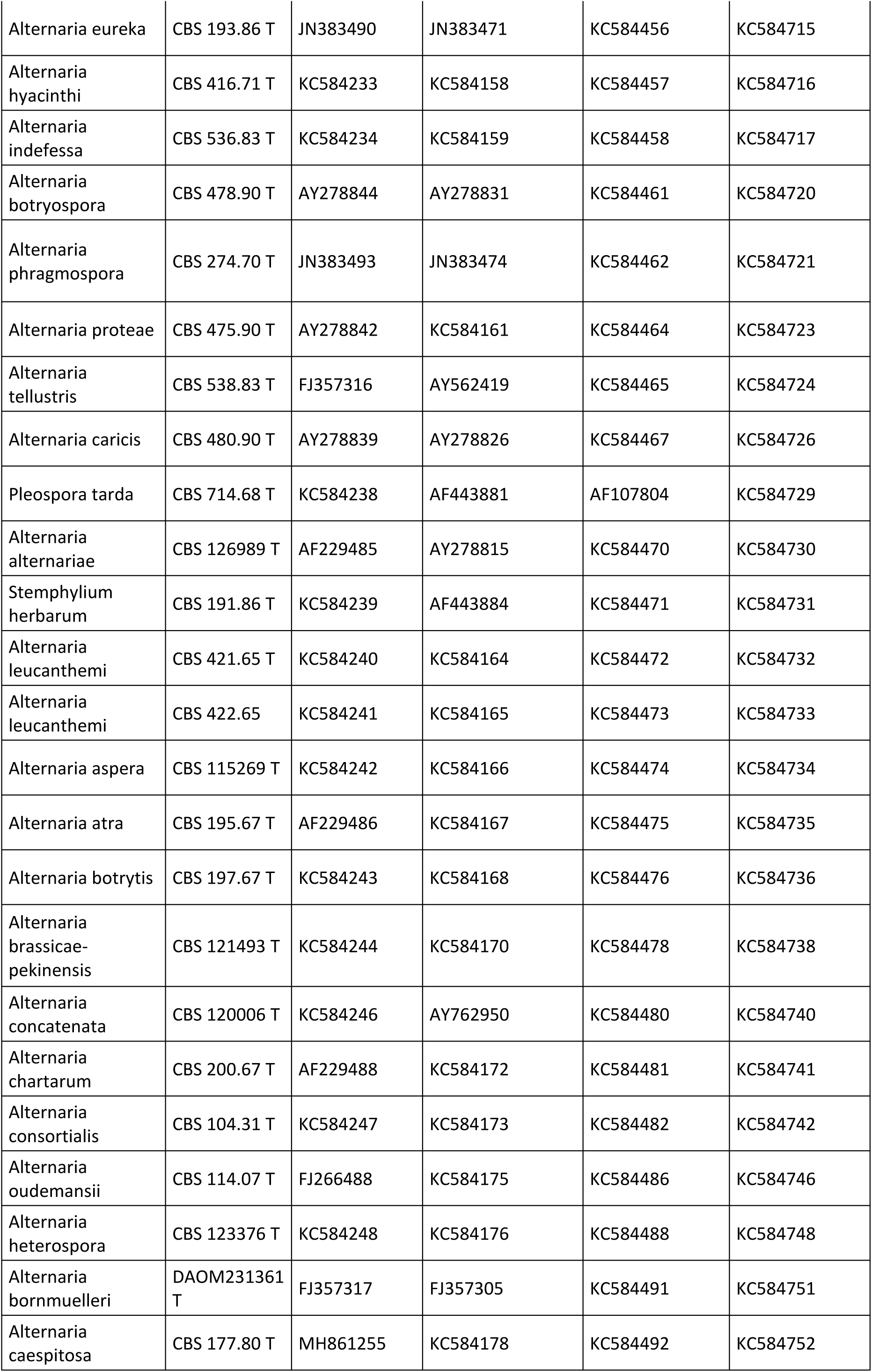
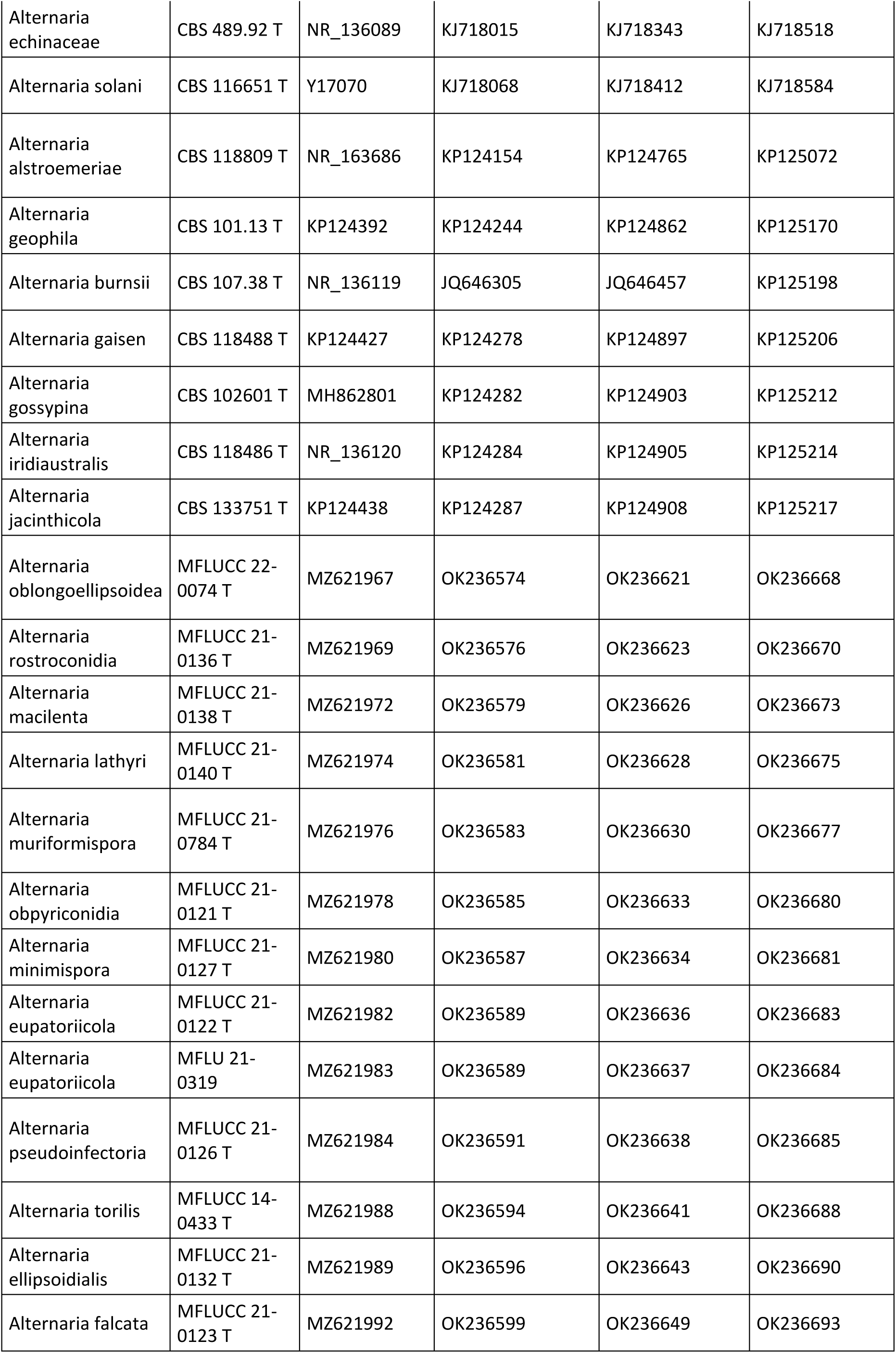
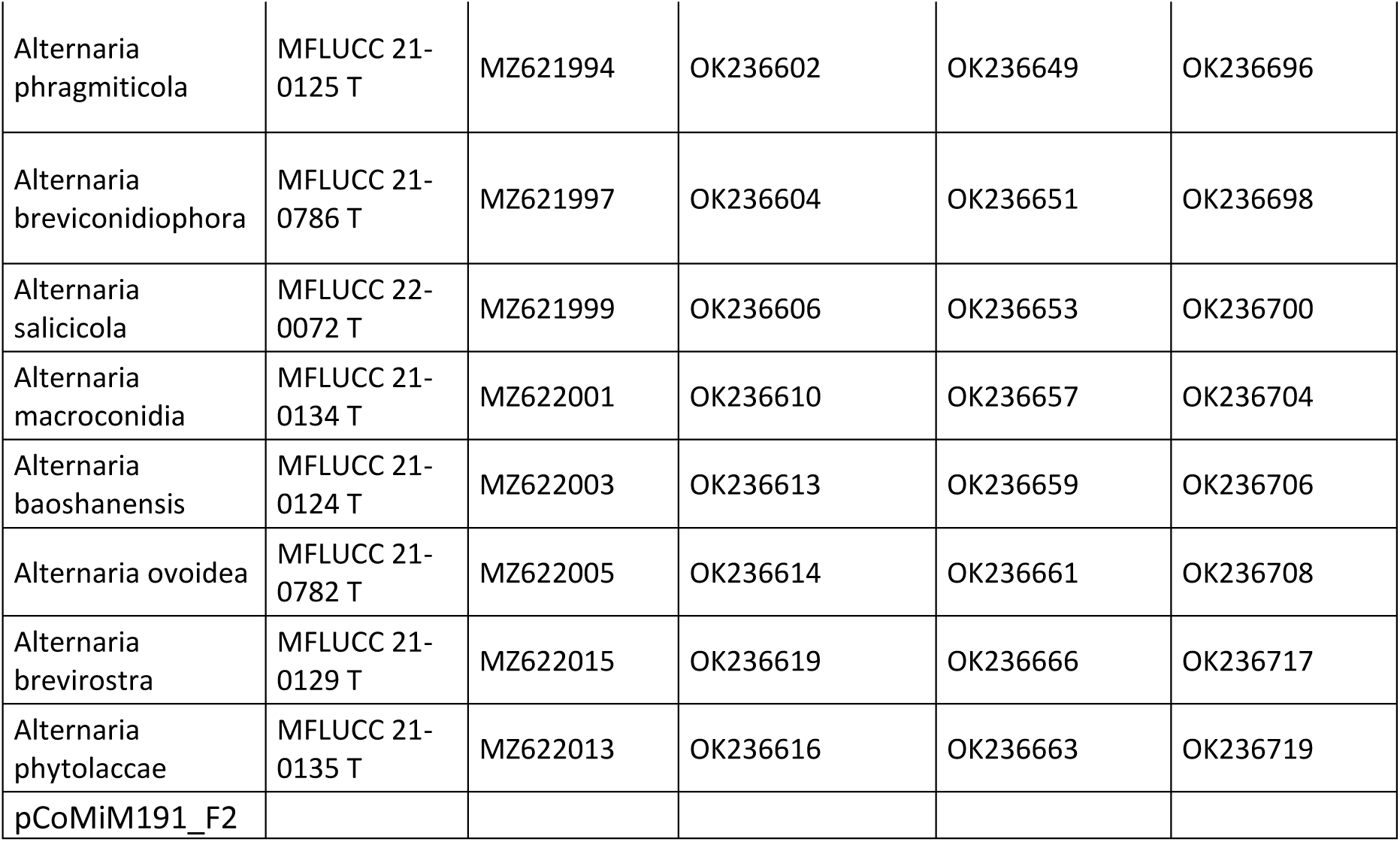
Strains used for the taxonomic identification within the *Alternaria* genus.

**Table S5:**
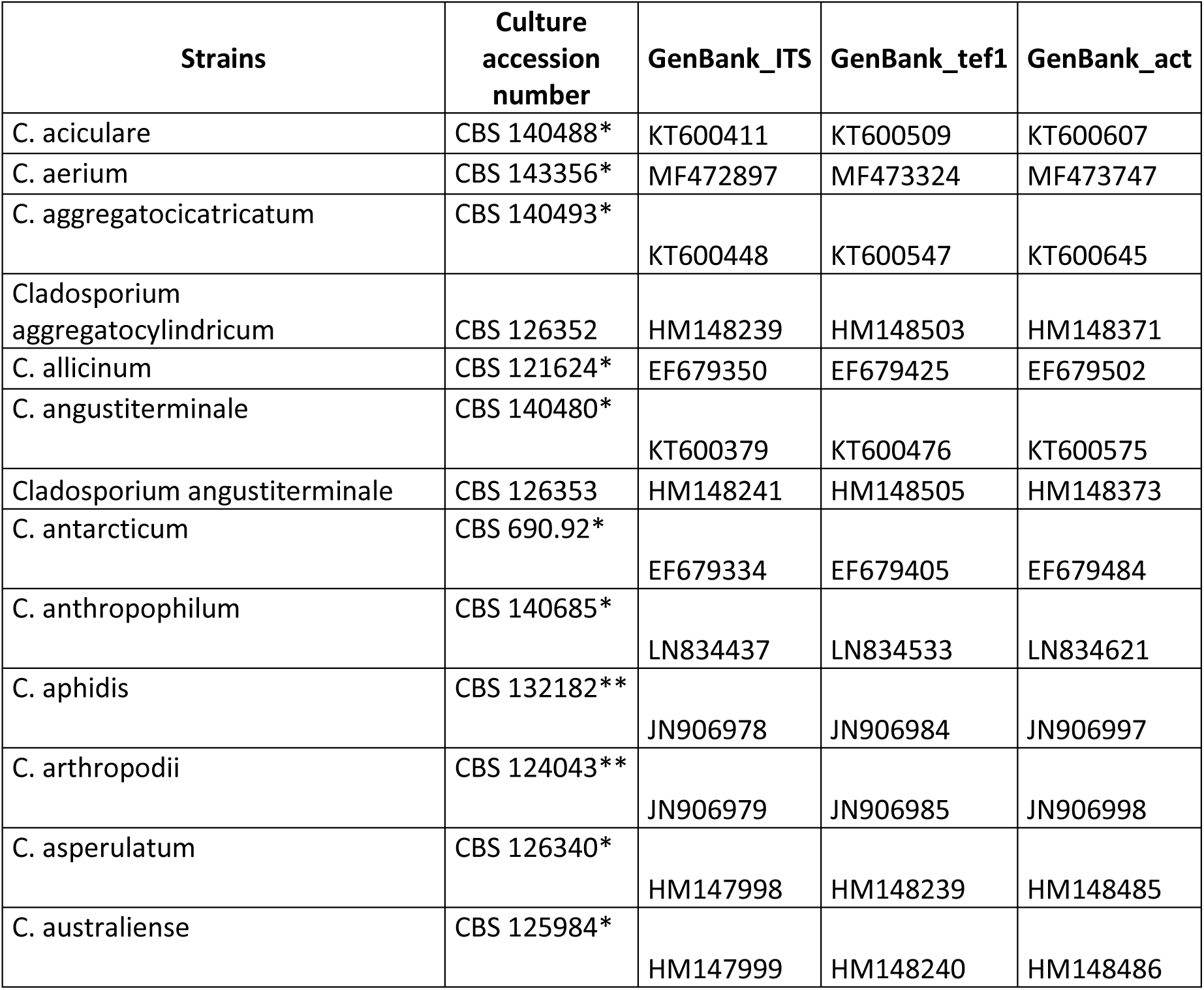
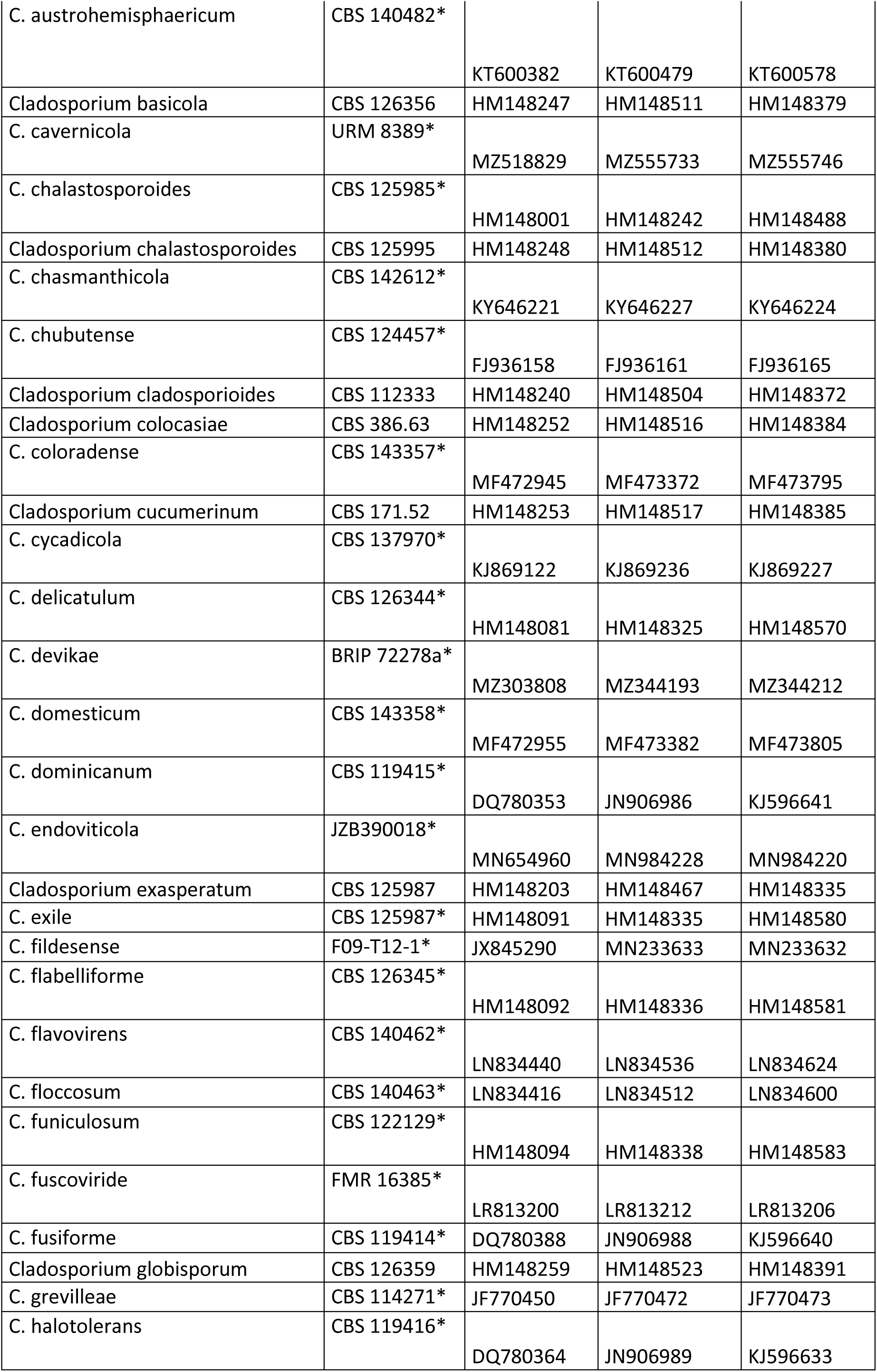
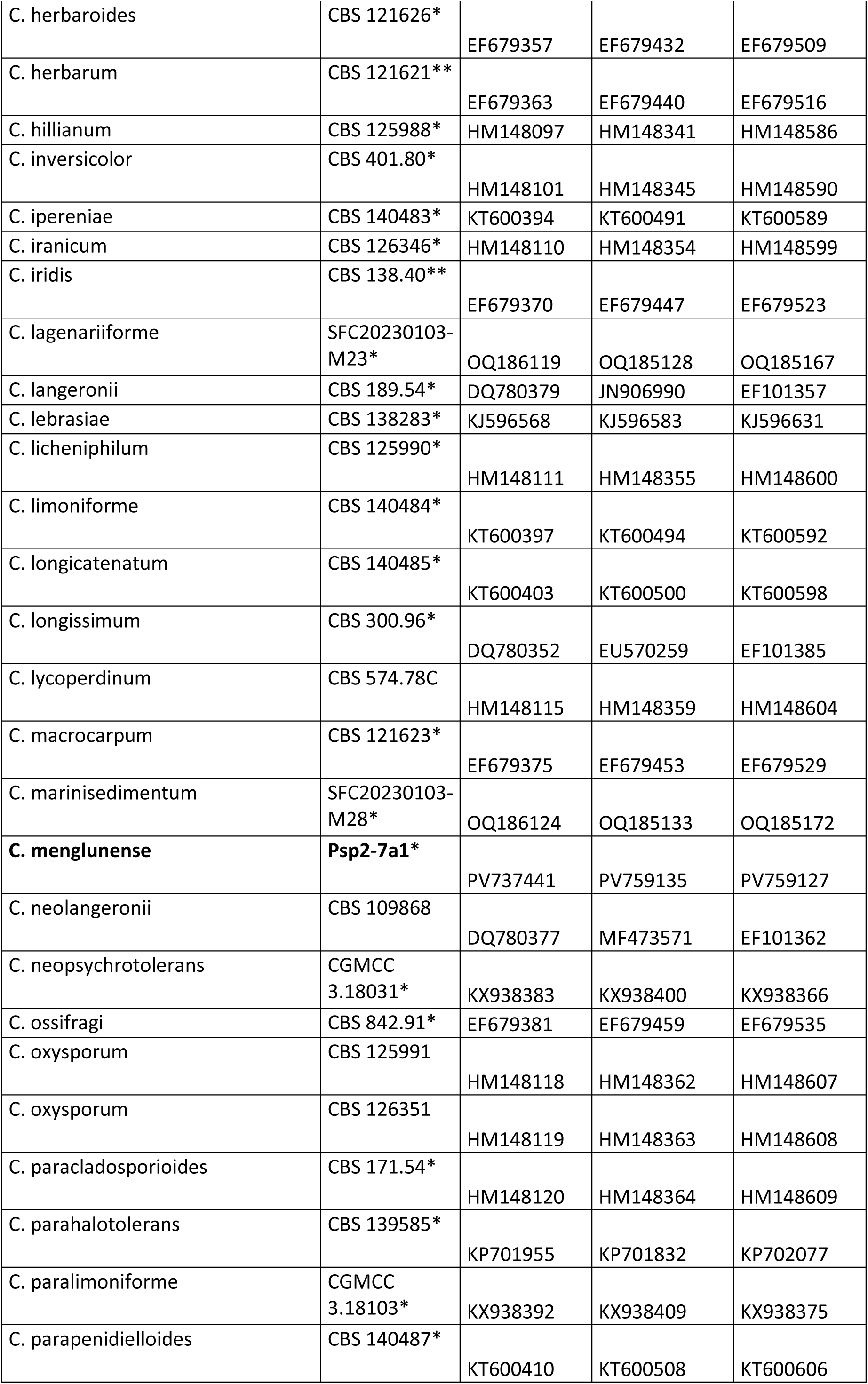
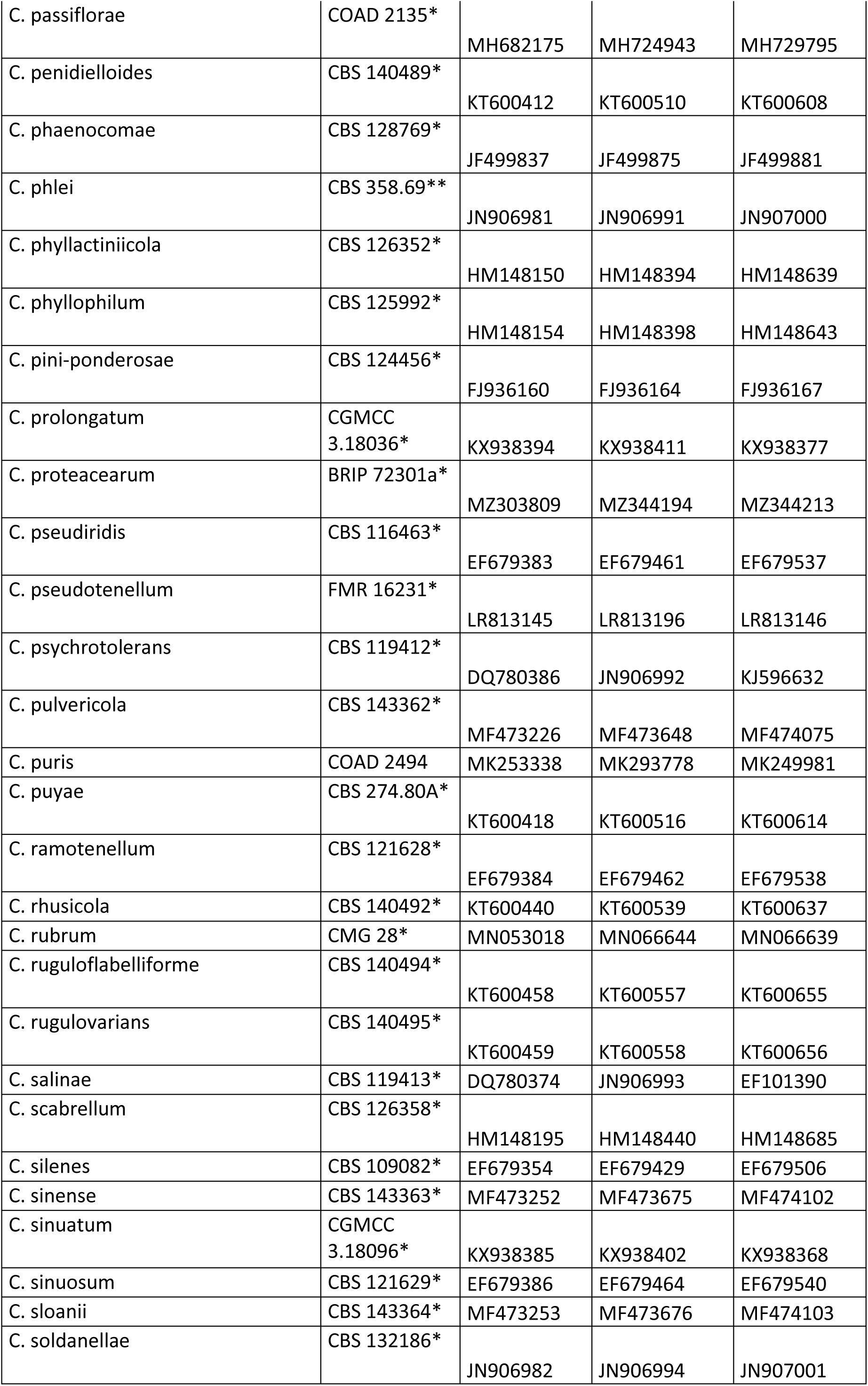
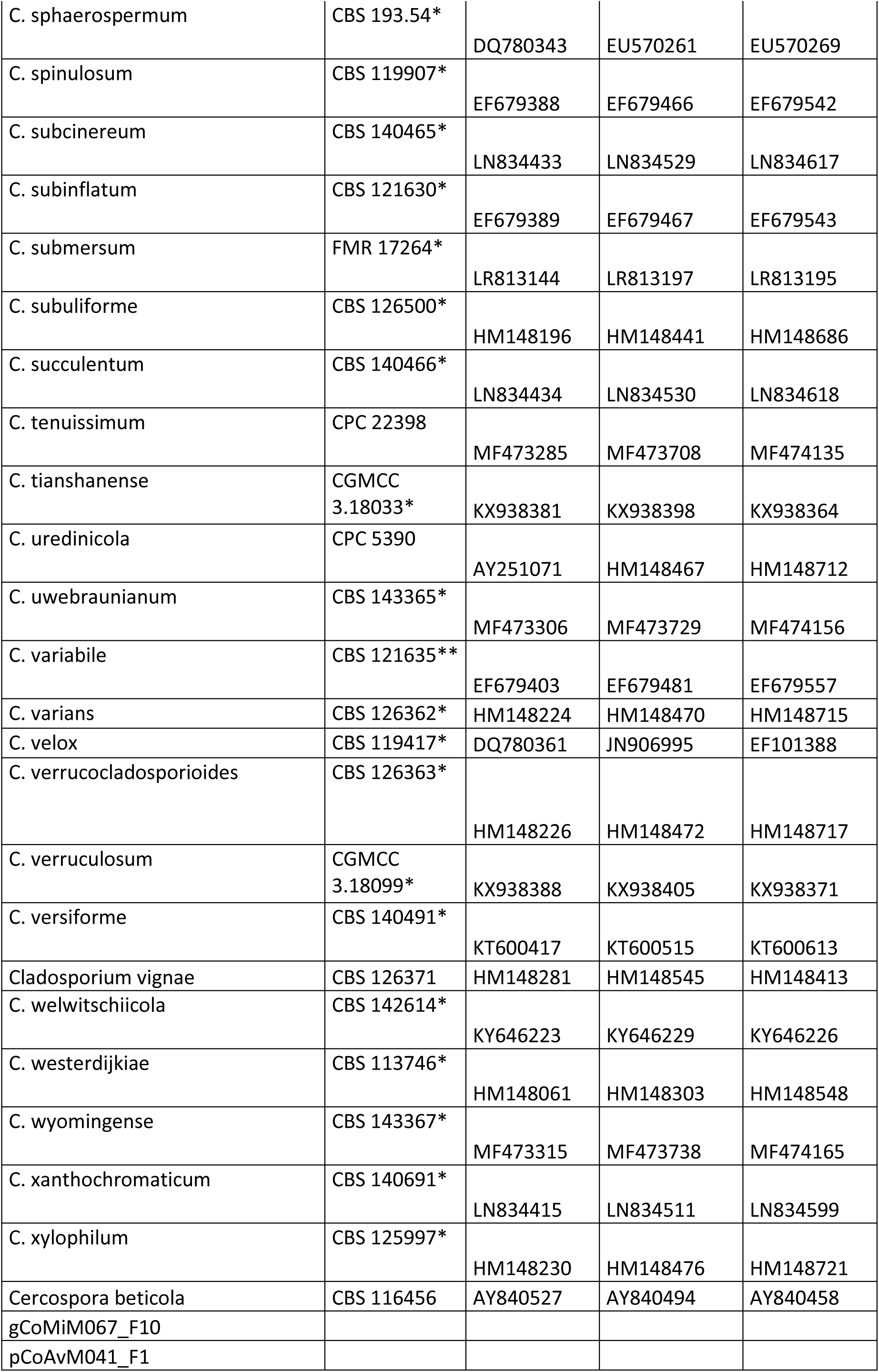
Strains used for the taxonomic identification within the *Cladosporium* genus.

**Table S6:** Strains used for the taxonomic identification within the *Mycosphaerella* genus.

| Strains | Culture accession number | GenBank ITS | GenBank LSU_D1-D3 | GenBank_act |
| --- | --- | --- | --- | --- |
| *M. colombiensis* | CBS 110969 | AY752149 | DQ204744 | DQ147639 |
| *M. crystallina* | CBS 681.95 | AY490757 | DQ204747 | DQ147636 |
| *M. gracilis* | CBS 243.94 | DQ267582 | DQ204750 | DQ147616 |
| *M. heimii* | CBS 110682 | AF309606 | DQ204751 | DQ147638 |
| *M. heimioides* | CBS 111364 | DQ267586 | DQ204752 | DQ147632 |
| *M. irregulariramosa* | CBS 114774 | AF309607 | DQ204754 | DQ147634 |
| *M. madeirae* | CBS 112895 | AY725553 | DQ204756 | DQ147641 |
| *M. lateralis* | CBS 110748 | AF173315 | DQ204768 | DQ147651 |
| *M. aurantia* | CBS 110500 | AY725531 | DQ246256 | DQ147610 |
| *M. africana* | CBS 116155 | DQ267577 | DQ246258 | DQ147608 |
| *M. keniensis* | CBS 111001 | AF309601 | DQ246259 | DQ147611 |
| *M. suberosa* | CBS 436.92 | AY626985 | DQ246235 | DQ147656 |
| *M. ambiphylla* | CBS 110499 | AY725530 | DQ246219 | DQ147669 |
| *M. molleriana* | CBS 111164 | AF309620 | DQ246220 | DQ147671 |
| *M. vespa* | CBS 117924 | DQ267590 | DQ246221 | DQ147668 |
| *M. parva* | CBS 110503 | AY626980 | DQ246243 | DQ147645 |
| *M. nubilosa* | CBS 116005 | AF309618 | DQ246228 | DQ147666 |
| *M. readeriellophora* | CBS 114240 | AY725577 | DQ246238 | DQ147658 |
| *M. cryptica* | CBS 110975 | AF309623 | DQ246222 | DQ147674 |
| *M. toledana* | CBS 113313 | AY725580 | DQ246230 | DQ147672 |
| *M. tasmaniensis* | CBS 111687 | DQ267591 | DQ246233 | DQ147676 |
| *M. mexicana* | CBS 110502 | AY725558 | DQ246237 | DQ147660 |
| *M. flexuosa* | CBS 111012 | AF309603 | DQ246232 | DQ147653 |
| *M. gregaria* | CBS 110501 | DQ267585 | DQ246251 | DQ147650 |
| *M. endophytica* | CBS 111519 | DQ267579 | DQ246255 | DQ147646 |
| *M. intermedia* | CBS 114415 | AY725547 | DQ246248 | DQ147627 |
| *M. marksii* | CBS 682.95 | DQ267587 | DQ246249 | DQ147624 |
| *M. parkii* | CBS 387.92 | AY626979 | DQ246245 | DQ147612 |
| *M. communis* | CBS 114238 | AY725541 | DQ246262 | DQ147655 |
| *Botryosphaeria ribis* | N/A | DQ267604 | DQ246263 | DQ267605 |
| gCoJuP058_F3 |  |  |  |  |

**Table S7:**
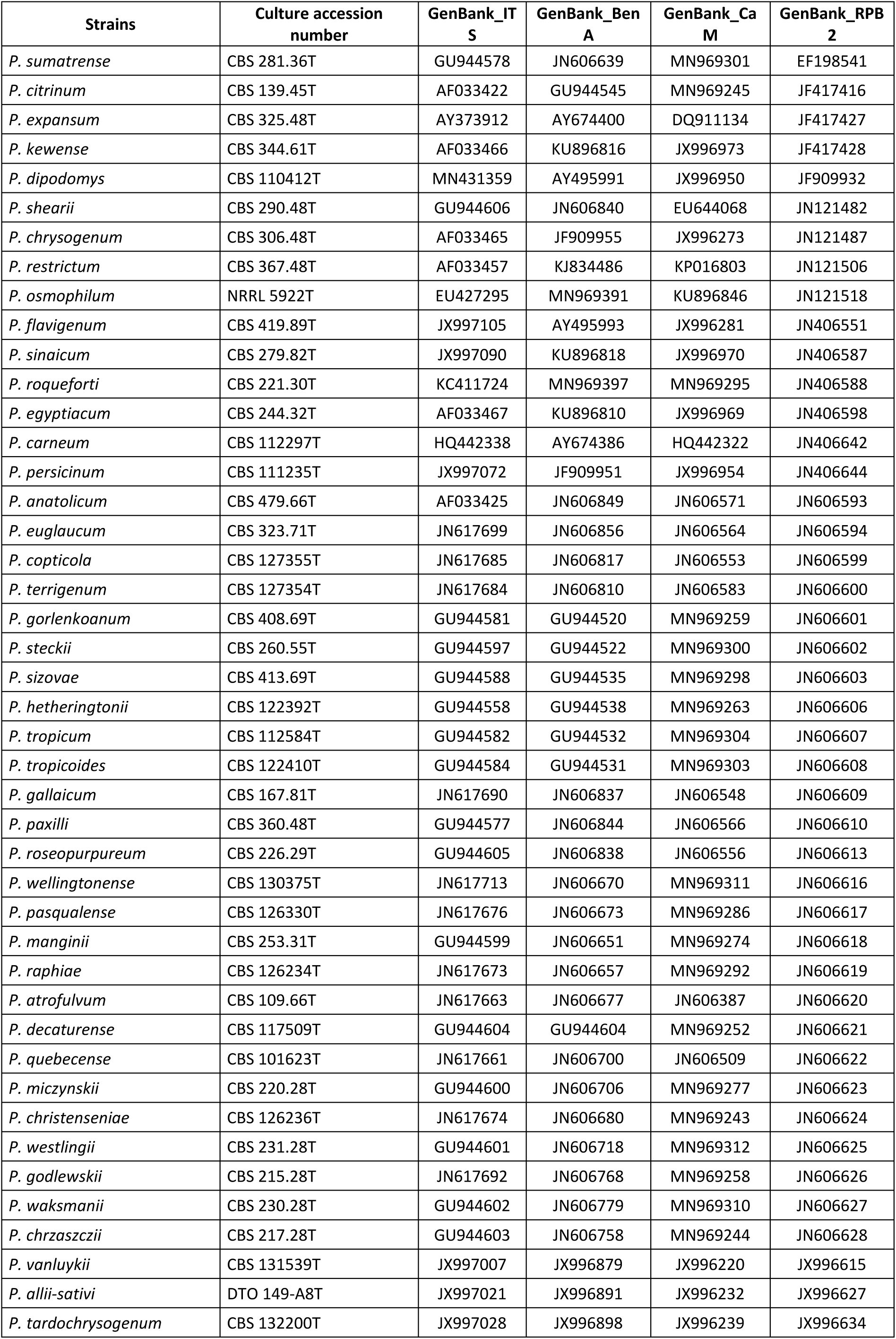
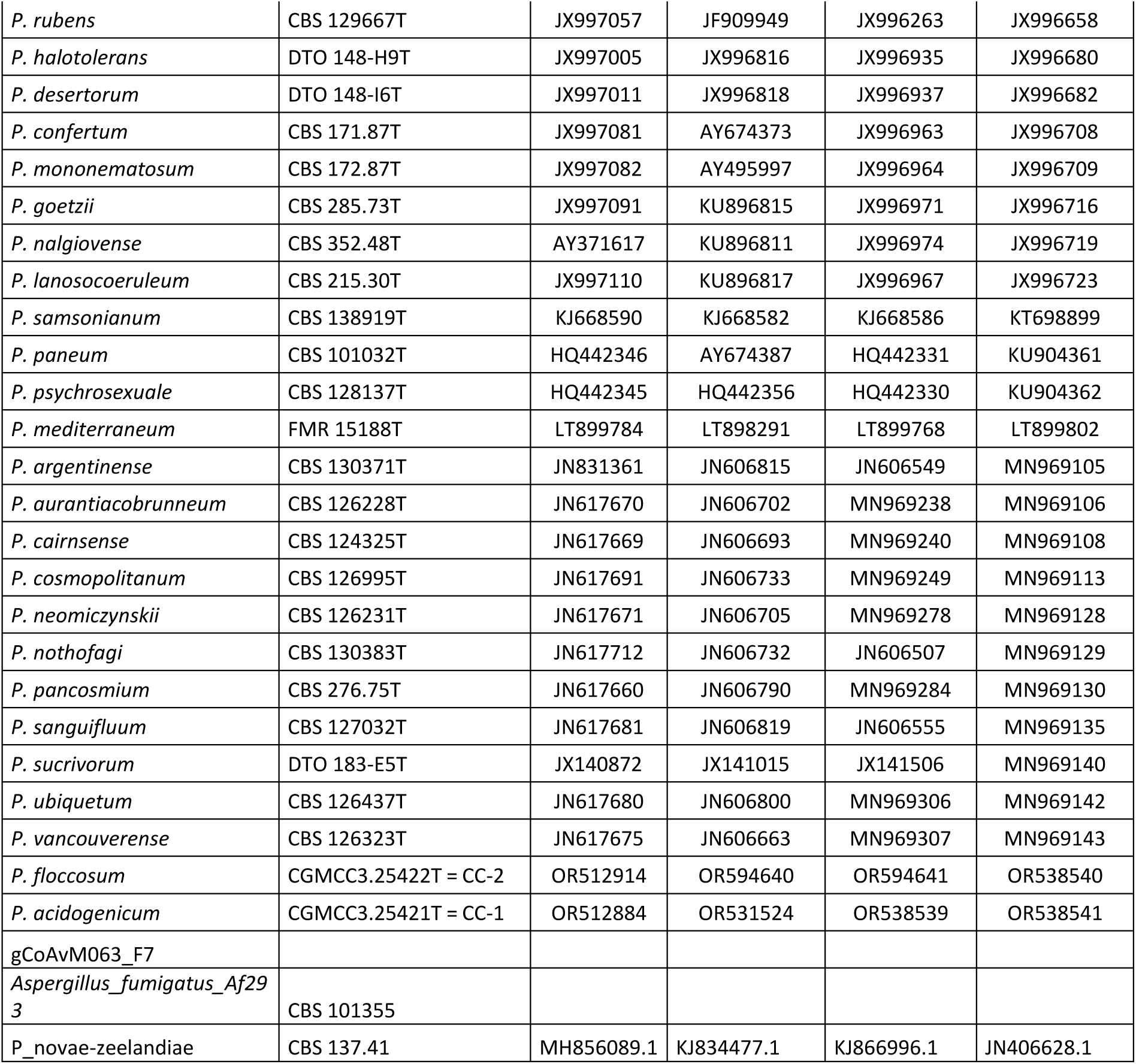
Strains used for the taxonomic identification within the *Penicillium* genus.

**Table S8:** Strains used for the taxonomic identification within the *Plectospaerella* genus.

| Strains | Culture accession number | GenBank ITS | GenBank LSU LROR-LR5 | GenBank TUB2 | GenBank tef1 |
| --- | --- | --- | --- | --- | --- |
| *Monilochaetes infuscans* | CBS 379.77 | LR026764 | GU180645 | — | — |
| *P. humicola* | CBS 423.66<sup>T</sup> | LR026811 | LR025949 | — | — |
| *P. hanneae* | CBS 144925 | LR590201 | LR590378 | — | — |
| *P. nauculaspora* | CGMCC 3.19656<sup>T</sup> | MK880439 | MK880424 | — | — |
| *P. guizhouensis* | CGMCC 3.19658<sup>T</sup> | MK880441 | MK880431 | — | — |
| *P. kunmingensis* | KUMCC 18-0181 | MK993014 | MK993015 | — | — |
| *P. oligotrophica* | CGMCC 3.15078<sup>T</sup> | KY399817 | JX508806 | KY421309 | KY421315 |
| *P. ramiseptata* | CBS 131743 | KY399819 | KY662252 | KY421306 | KY421318 |
| *P. citrullae* | CBS 131741<sup>T</sup> | KY399822 | KY662255 | KY421308 | KY421320 |
| *P. plurivora* | CBS131742<sup>T</sup> | KY399826 | KY662248 | KY421298 | KY421324 |
| *P. sinensis* | ACCC 39144 | KX527889 | KX527892 | KY421313 | KY421326 |
| *Plectosphaerella alismatis* | CBS 113362<sup>T</sup> | KY399815 | KY662261 | KY421304 | KY421328 |
| *P. melonis* | CBS 489.96<sup>T</sup> | KY399813 | KY662260 | KY421295 | KY421329 |
| *P. cucumerina* | 3F17 | KY401602 | KY399831 | KY421300 | KY421331 |
| *P. populi* | CBS139623<sup>T</sup> | KY399828 | KY662257 | KY421311 | KY421336 |
| *P. delsorboi* | CBS 116708<sup>T</sup> | KY499237 | EF543843 | KY499240 | KY499238 |
| *P. endophytica* | YMF 1.04701 | MW024054 | MW024052 | MW029608 | MW029607 |
| *P. sichuanensis* | YMF 1.05081 | MW027539 | MW027535 | MW441780 | MW309880 |
| *P. pauciseptata* | YMF 1.04725 | MW024104 | MW466575 | MW441784 | MW441779 |
| gCoJuM088_F4 |  |  |  |  |  |

**Table S9:** Strains used for the taxonomic identification within the *Sarocladium genus*.

| Strains | Culture accession number | GenBank ITS | GenBank LSU |
| --- | --- | --- | --- |
| *Chlamydocillium curvulum* | CBS 430.66* | HE608638 | HE608656 |
| *S. glaucum* | CBS 796.69* | OQ429841 | HE608657 |
| *S. oryzae* | CBS 180.74* | HG965026 | HG965047 |
| *S. bifurcatum* | UTHSC 05-3311* | HG965009 | HG965057 |
| *S. hominis* | UTHSC 04-1034* | HG965012 | HG965060 |
| *S. hominis* | UTHSC 04-3464 | HG965013 | HG965061 |
| *S. gamsii* | CBS 707.73* | HG965015 | HG965063 |
| *S. implicatum* | CBS 959.72* | HG965023 | HG965072 |
| *S. pseudostrictum* | UTHSC 02-1892* | HG965029 | HG965073 |
| *S. subulatum* | MUCL 9939* | HG965031 | HG965075 |
| *S. summerbellii* | CBS 430.70* | HG965034 | HG965078 |
| *S. bactrocephalum* | CBS 749.69* | HG965006 | HQ231994 |
| *S. kiliense* | CBS 122.29* | AJ621775 | HQ232052 |
| *S. ochraceum* | CBS 428.67* | HG965025 | HQ232070 |
| *S. strictum* | CBS 346.70* | FN691453 | HQ232141 |
| *S. zeae* | CBS 800.69* | FN691451 | HQ232152 |
| *S. spinificis* | Z0106 Ex-type | KF269096 | JQ954463 |
| *S. brachiariae* | CGMCC 2192* | EU880834 | KP715271 |
| *S. terricola* | CBS 243.59* | MH857853 | MH869389 |
| *S. bacillisporum* | CBS 425.67* | HE608639 | MH870718 |
| *S. dejongiae* | CBS 144929* | MK069419 | MK069415 |
| *S. theobromae* | CBS 113440 Ex-type | OQ429856 | OQ430108 |
| ***S. menglaense* sp. nov.** | **KO1-8\*** | PV737445 | PV737452 |
| ***S. yunnanense* sp. nov.** | **WF8-2a1\*** | PV737449 | PV737456 |
| gCoAvM140_F5 |  |  |  |

**Table S10:**
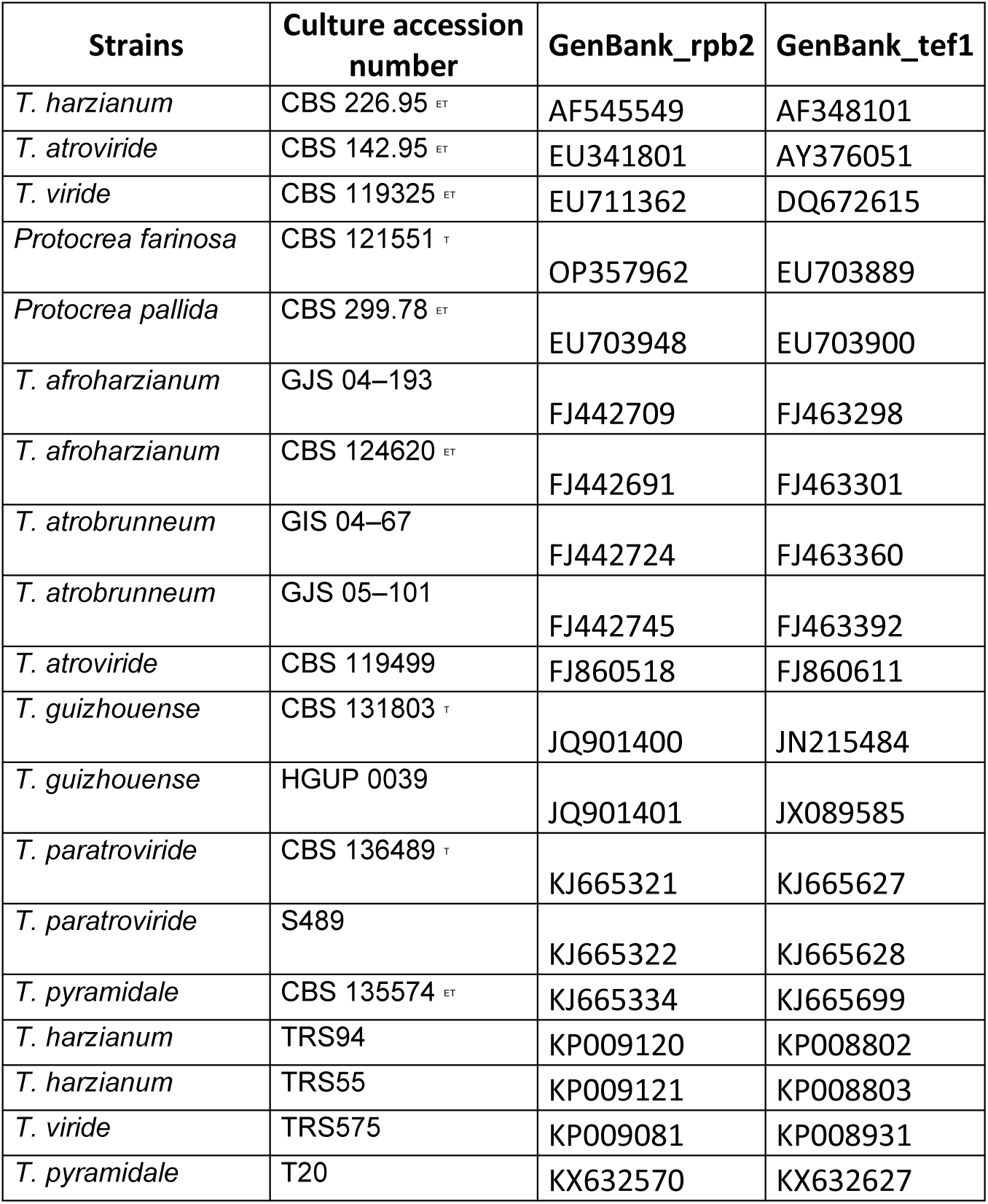
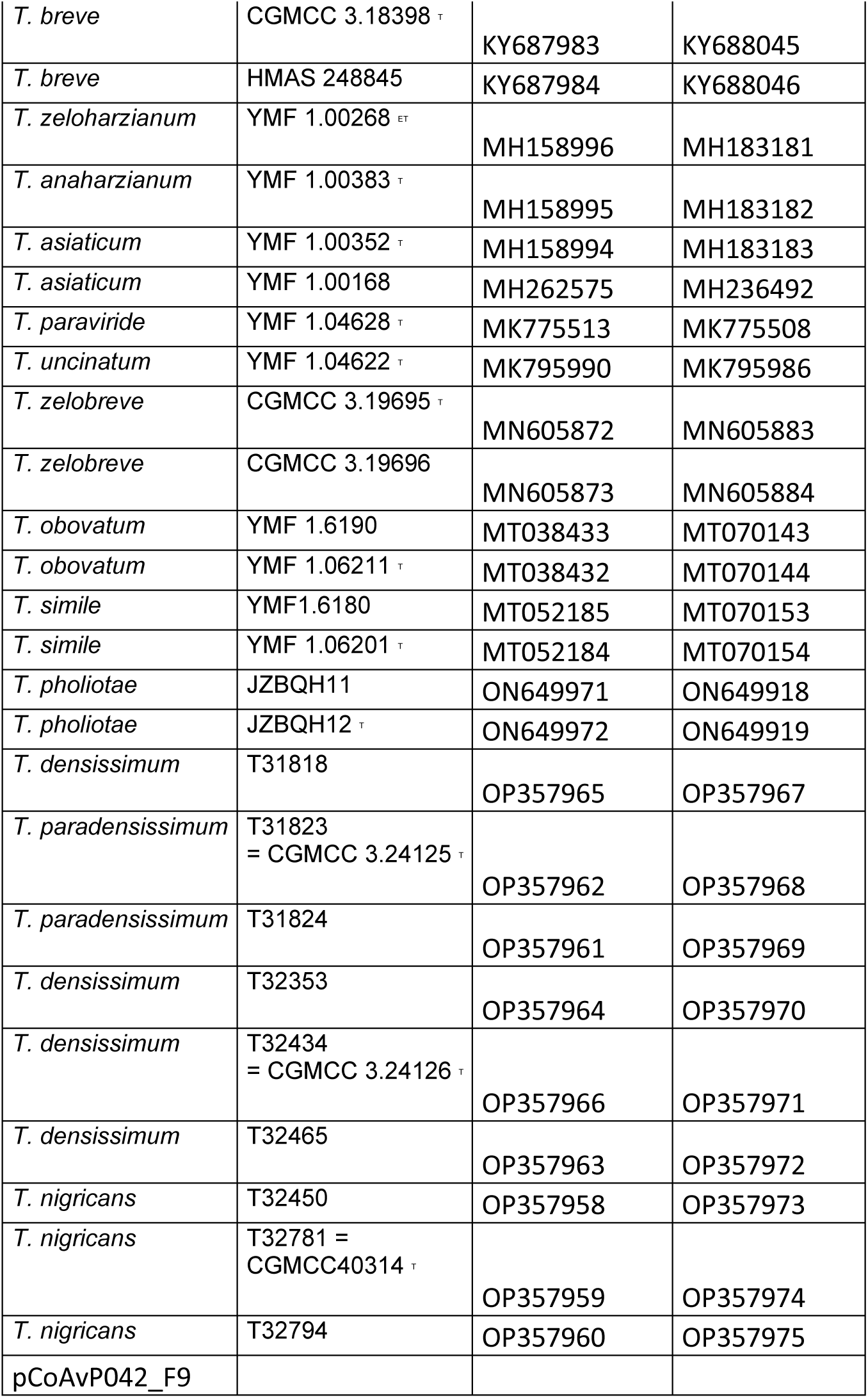
Strains used for the taxonomic identification within the *Trichoderma genus*.

**Table S11:**
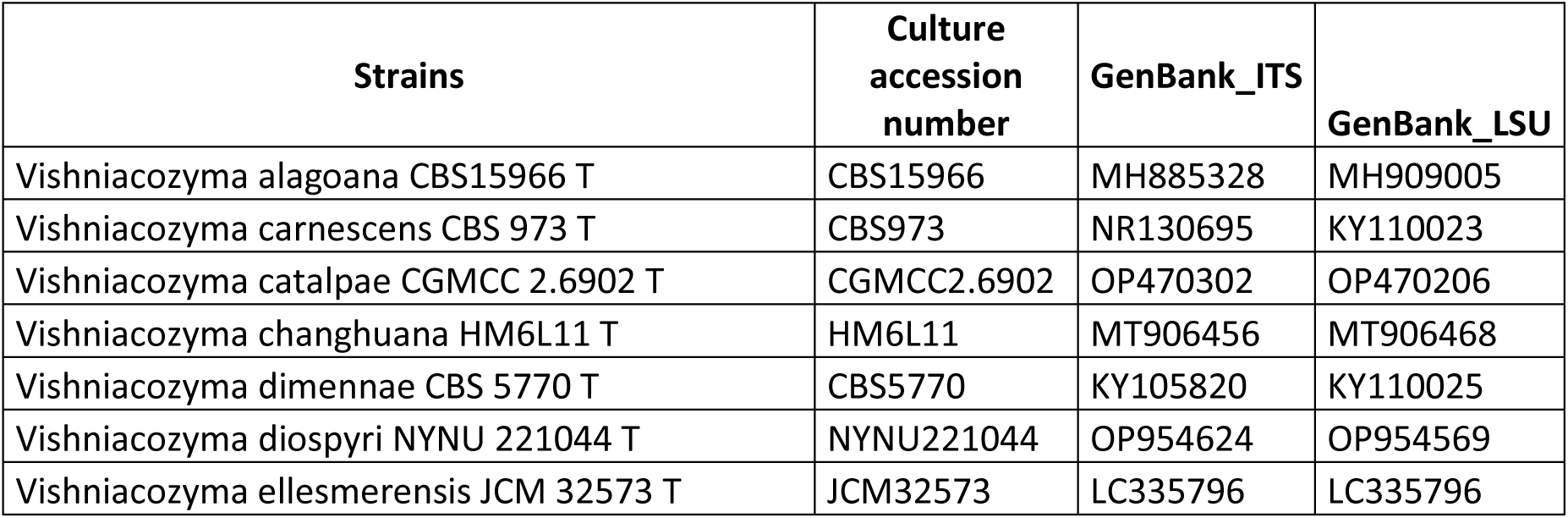
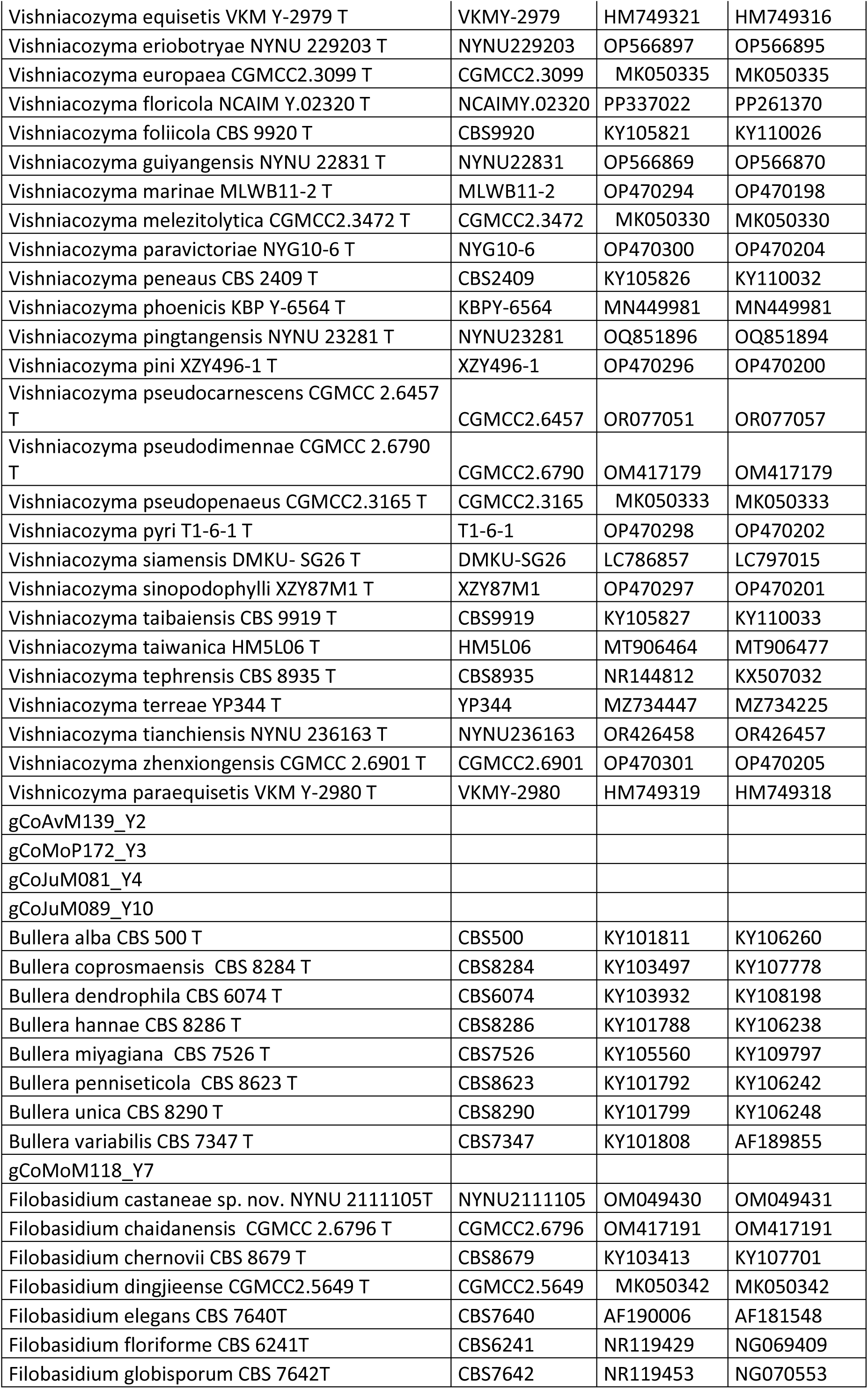
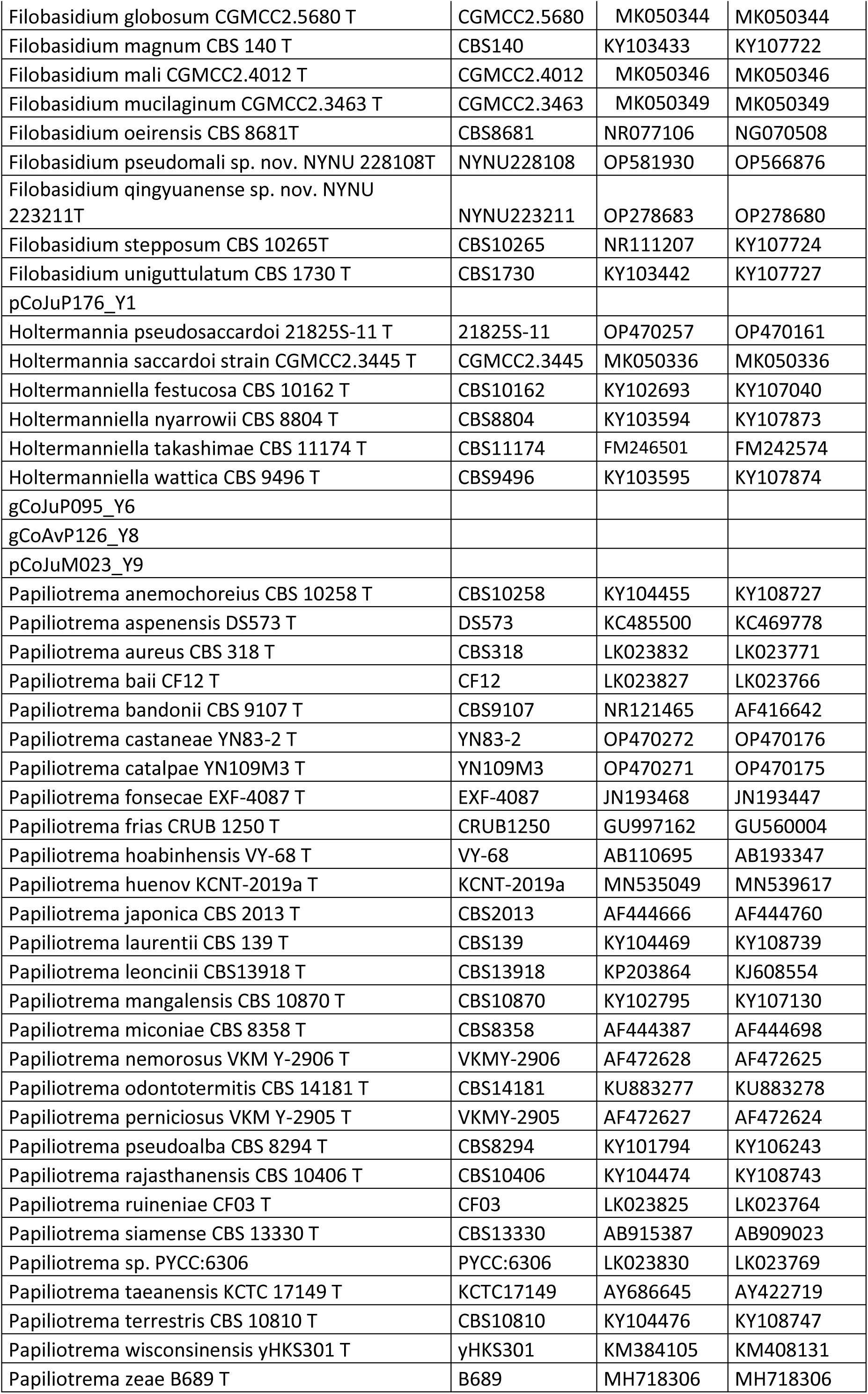
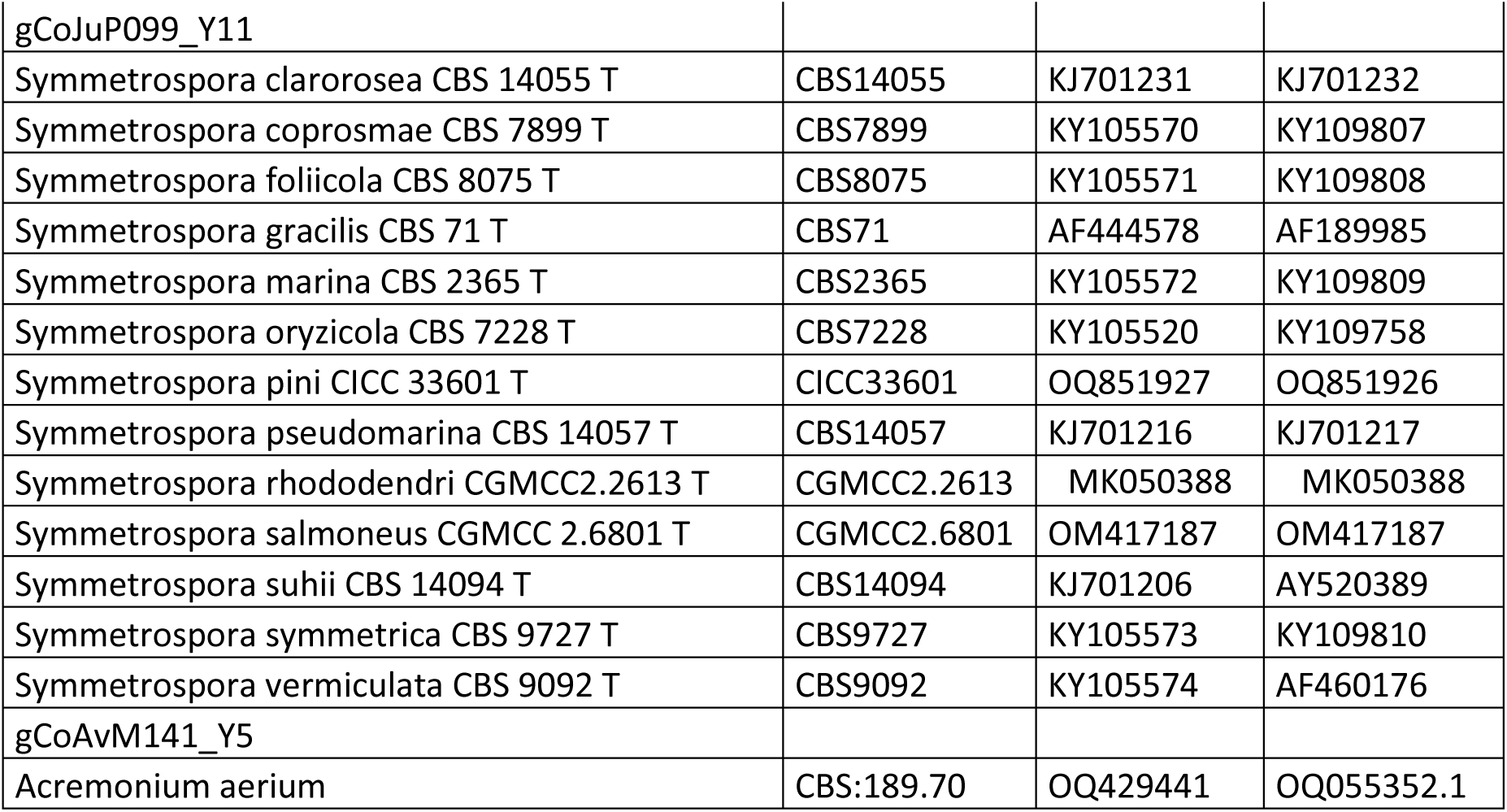
Strains used for the taxonomic identification within the different yeast genus.

**Table S12:**
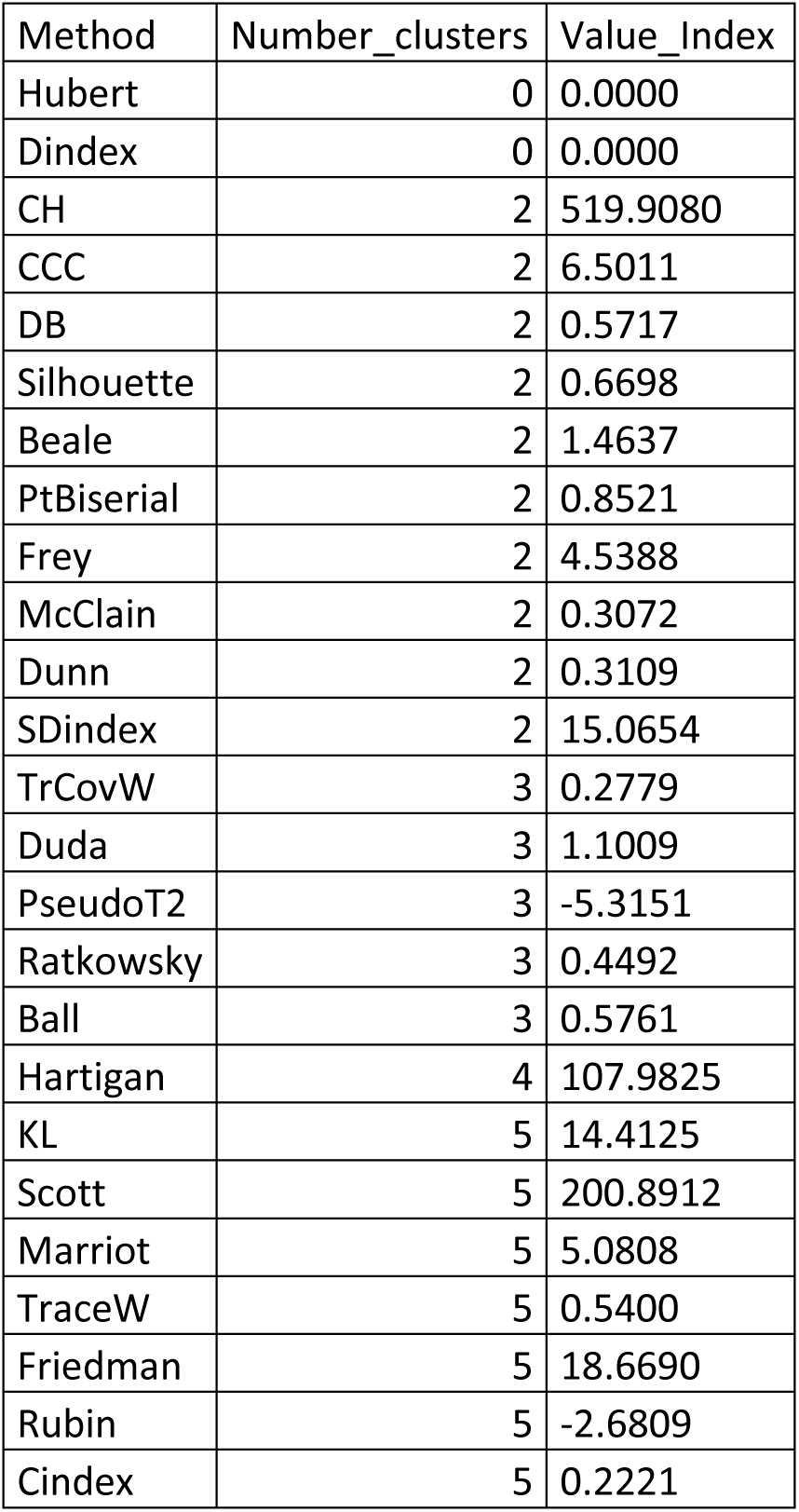
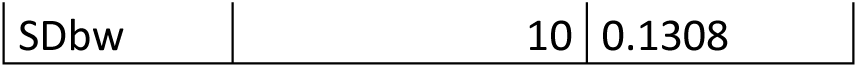
Output of the NbClust R package used to choose the optimal number of clusters of seedling bacterial communities.

| Method | Number_clusters | Value_Index |
| --- | --- | --- |
| Hubert | 0 | 0.0000 |
| Dindex | 0 | 0.0000 |
| CH | 2 | 519.9080 |
| CCC | 2 | 6.5011 |
| DB | 2 | 0.5717 |
| Silhouette | 2 | 0.6698 |
| Beale | 2 | 1.4637 |
| PtBiserial | 2 | 0.8521 |
| Frey | 2 | 4.5388 |
| McClain | 2 | 0.3072 |
| Dunn | 2 | 0.3109 |
| SDindex | 2 | 15.0654 |
| TrCovW | 3 | 0.2779 |
| Duda | 3 | 1.1009 |
| PseudoT2 | 3 | -5.3151 |
| Ratkowsky | 3 | 0.4492 |
| Ball | 3 | 0.5761 |
| Hartigan | 4 | 107.9825 |
| KL | 5 | 14.4125 |
| Scott | 5 | 200.8912 |
| Marriot | 5 | 5.0808 |
| TraceW | 5 | 0.5400 |
| Friedman | 5 | 18.6690 |
| Rubin | 5 | -2.6809 |
| Cindex | 5 | 0.2221 |
| SDbw | 10 | 0.1308 |

**Table S13:** Main metrics of the three co-occurrence networks, divided between entire networks and giant and peripheral components.

| Network | Component | Total node number | Inoculated strain number | Differentially abundant ASVs number | Node degree | Normalized node degree | Density |
| --- | --- | --- | --- | --- | --- | --- | --- |
| Full | Total | 6890 | 36 | 82 | 4443 | 0.64 | 1.87E-04 |
| Full | Giant | 2400 | 29 | 82 | 4017 | 1.67 | 1.40E-03 |
| Full | Periph | 4490 | 7 | 0 | 426 | 0.09 | 4.23E-05 |
| Cluster1 | Total | 3189 | 34 | 77 | 2135 | 0.67 | 4.20E-04 |
| Cluster1 | Giant | 1220 | 27 | 73 | 1885 | 1.55 | 2.53E-03 |
| Cluster1 | Periph | 1969 | 7 | 4 | 250 | 0.13 | 1.29E-04 |
| Cluster2 | Total | 3211 | 32 | 81 | 2213 | 0.69 | 4.29E-04 |
| Cluster2 | Giant | 1263 | 23 | 74 | 1903 | 1.51 | 2.39E-03 |
| Cluster2 | Periph | 1948 | 9 | 7 | 310 | 0.16 | 1.63E-04 |

## Supplementary material and methods: SynCom Composition Space Analysis

### Strain Pool and Experimental Design

The synthetic community (SynCom) strain pool comprised 45 microbial isolates distributed across three kingdoms: 24 bacteria (B1–B24), 10 fungi (F1–F10), and 11 yeasts (Y1–Y11). Each SynCom was assembled under a balanced combinatorial constraint of exactly 4 bacteria, 4 fungi, and 4 yeasts (4B+4F+4Y), yielding a total theoretical design space of:

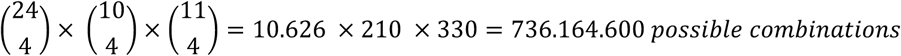

Nineteen SynComs were experimentally designed, including randomly assembled communities (Rand1–Rand10), ecologically informed assemblages (MostAbun, MostAbunSeed, MostAbunSeedling, FastGrowth, Rare1, Rare2, AbunGrad), and phylogenetically diverse assemblages (PhylDiv1, PhylDiv2).

### Theoretical Space Sampling

To represent the full theoretical composition space, 5,000 communities were generated by random sampling without replacement of 4 strains from each kingdom pool (set.seed = 42), representing approximately 0.00068% of all possible combinations. This Monte Carlo approach served as a tractable proxy for the full combinatorial space in all subsequent analyses.

### Compositional Dissimilarity and Ordination

Pairwise compositional dissimilarity between all communities (theoretical and effective) was quantified using the binary Jaccard distance: 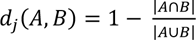

computed via the *vegdist* function (vegan; Oksanen et al, 2007). For two 12-member communities sharing *s* strains, this simplifies to *d_j_* = (24 − 2*s*)/(24 − *s*), yielding distances anchored to discrete integer strain-sharing levels (*e.g., d_j_* = 0.5 at *s* = 8 ; *d_j_* = 0.667 at *s* = 6).

Pairwise Jaccard distances were first computed among the 19 experimentally designed SynComs alone, and subsequently on a combined presence–absence matrix constructed by row-binding the 5,000 theoretical communities and the 19 designed SynComs (5,019 × 45). Both distance matrices were subjected to Principal Coordinates Analysis (PCoA) using the pcoa function (ape package) with Cailliez correction for negative eigenvalues, retaining the first two axes for visualisation. Designed SynComs were overlaid on the theoretical space point cloud using the combined PCoA ordination.

### Dispersion of Designed SynComs Relative to Random Sets

To assess whether the 19 designed SynComs collectively span more compositional space than expected by chance, their mean pairwise Jaccard dissimilarity was computed as an observed dispersion statistic. A permutation null distribution was generated by repeatedly drawing 19 random 4B+4F+4Y communities from the full strain pool (n = 5,000 permutations; set.seed = 42) and computing their mean pairwise Jaccard dissimilarity. A one-sided p-value was estimated as the proportion of permuted sets yielding a mean pairwise Jaccard dissimilarity greater than or equal to the observed value.

### Nearest-Neighbour Distance to Theoretical Composition Space

To quantify how well each designed SynCom was embedded within the theoretical composition space, the Jaccard distance from each of the 19 designed SynComs to its nearest neighbour among the 5,000 theoretical communities was extracted from the full cross-distance submatrix. Lower nearest-neighbour (NN) distances indicate greater compositional representativeness of the theoretical space. As a null reference, the self-NN distance distribution of the theoretical space was computed by identifying, for each of the 5,000 random communities, the distance to its closest neighbour among all remaining random communities. The two distributions were compared visually, and a one-sided p-value was estimated as the proportion of theoretical self-NN distances greater than or equal to the mean NN distance of the designed SynComs. Individual NN distances were additionally ranked per SynCom and stratified by design strategy (A priori vs. Random) to identify which communities occupied peripheral versus representative regions of the composition space, using the mean theoretical self-NN distance as a reference threshold.

