## Supplementary material and methods for "Seed-applied multi-kingdom synthetic communities selectively reshape bacterial communities and highlight key criteria for strain selection"

### Supplementary material and methods: SynCom Composition Space Analysis

**Strain Pool and Experimental Design**

The synthetic community (SynCom) strain pool comprised 45 microbial isolates distributed across three kingdoms: 24 bacteria (B1–B24), 10 fungi (F1–F10), and 11 yeasts (Y1–Y11). Each SynCom was assembled under a balanced combinatorial constraint of exactly 4 bacteria, 4 fungi, and 4 yeasts (4B+4F+4Y), yielding a total theoretical design space of:

$$\binom{24}{4}\times\binom{10}{4}\times\binom{11}{4}=10.626 \times210 \times330=736.164.600 possible combinations$$

Nineteen SynComs were experimentally designed, including randomly assembled communities (Rand1–Rand10), ecologically informed assemblages (MostAbun, MostAbunSeed, MostAbunSeedling, FastGrowth, Rare1, Rare2, AbunGrad), and phylogenetically diverse assemblages (PhylDiv1, PhylDiv2).

**Theoretical Space Sampling**

To represent the full theoretical composition space, 5,000 communities were generated by random sampling without replacement of 4 strains from each kingdom pool (set.seed = 42), representing approximately 0.00068% of all possible combinations. This Monte Carlo approach served as a tractable proxy for the full combinatorial space in all subsequent analyses.

**Compositional Dissimilarity and Ordination**

Pairwise compositional dissimilarity between all communities (theoretical and effective) was quantified using the binary Jaccard distance: $d_{j}\left( A, B \right)=1- \frac{\left| A\cap B \right|}{\left| A\cup B \right|}$
