## Supplementary figures for "Seed-applied multi-kingdom synthetic communities selectively reshape bacterial communities and highlight key criteria for strain selection"

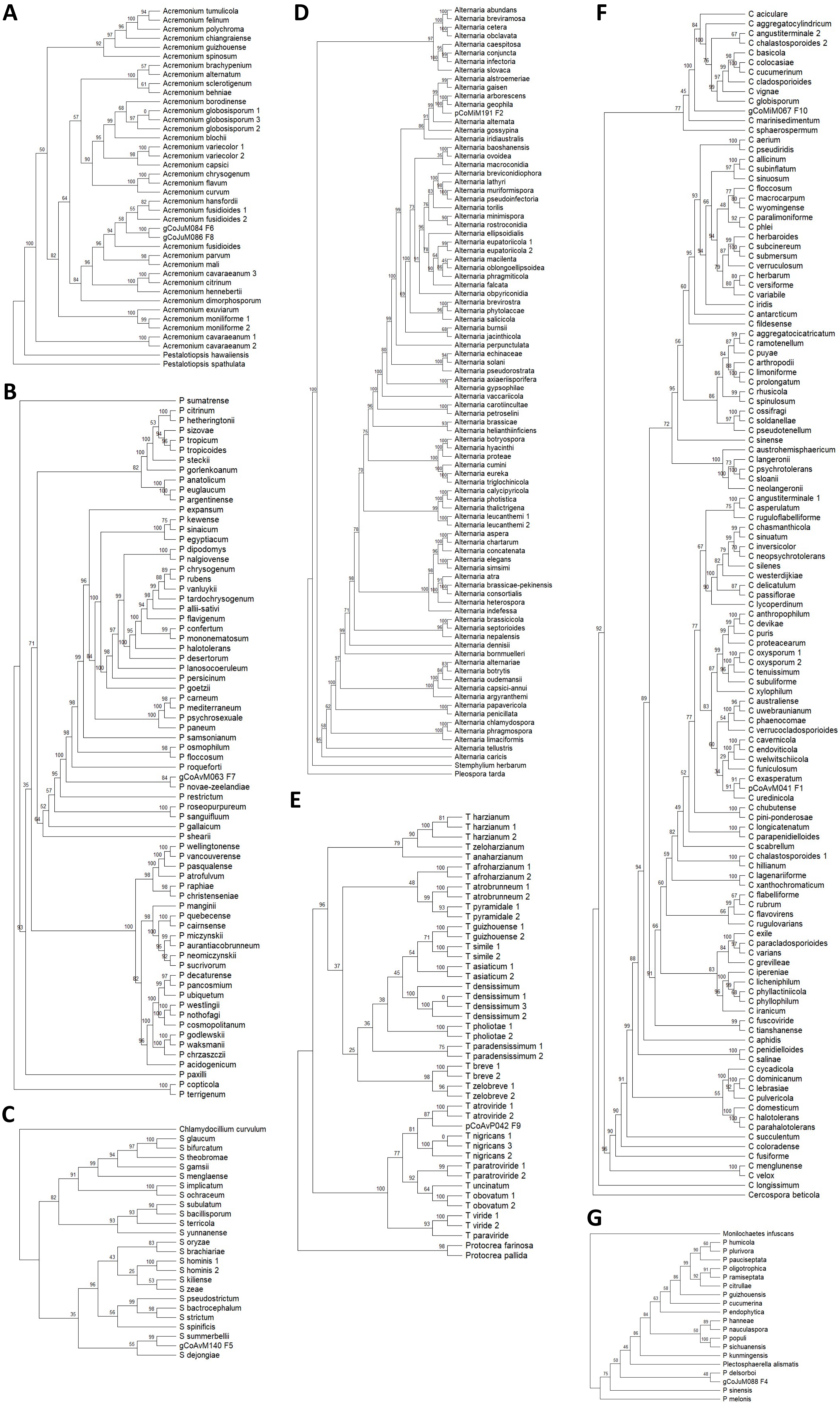


Supplementary Fig. 1 : Phylogenetic trees of the genus (A) *Acremonium*, (B) *Penicillium*, (C) *Sarocladium*, (D) *Alternaria*, (E) *Trichoderma*, (F) *Cladosporium* and (G) *Plectosphaerella* using the IQ-TREE Ultrafast bootstrap (1000 iterations)

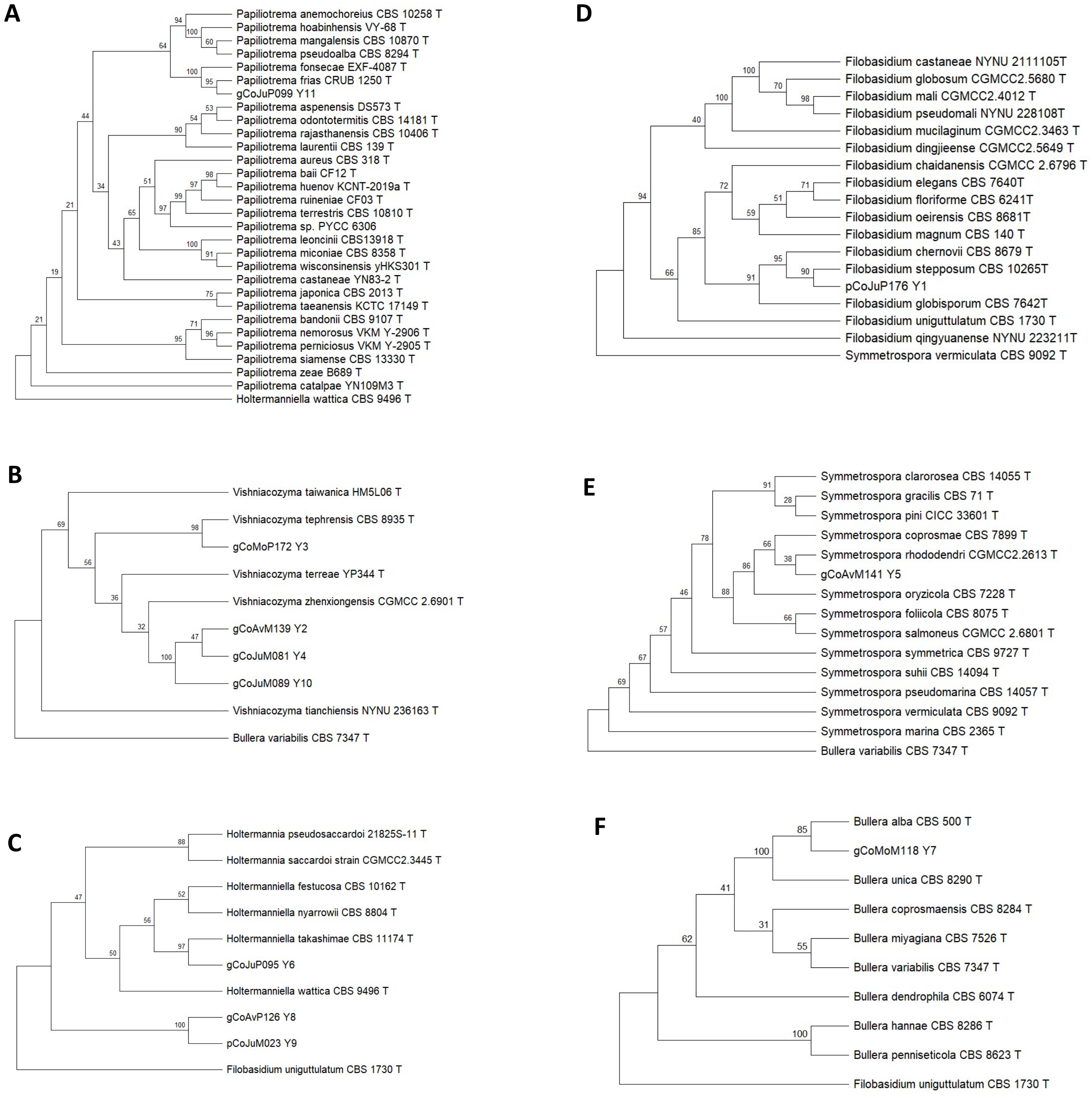


Supplementary Fig. 2: Phylogenetic tree of the genus (A) Papiliotrema, (B) Vishniacozyma, (C) Holtermaniella, (D) Filobasidium, (E) Symmetrospora and (F) Bullera using the IQ-TREE Ultrafast bootstrap (1000 iterations)

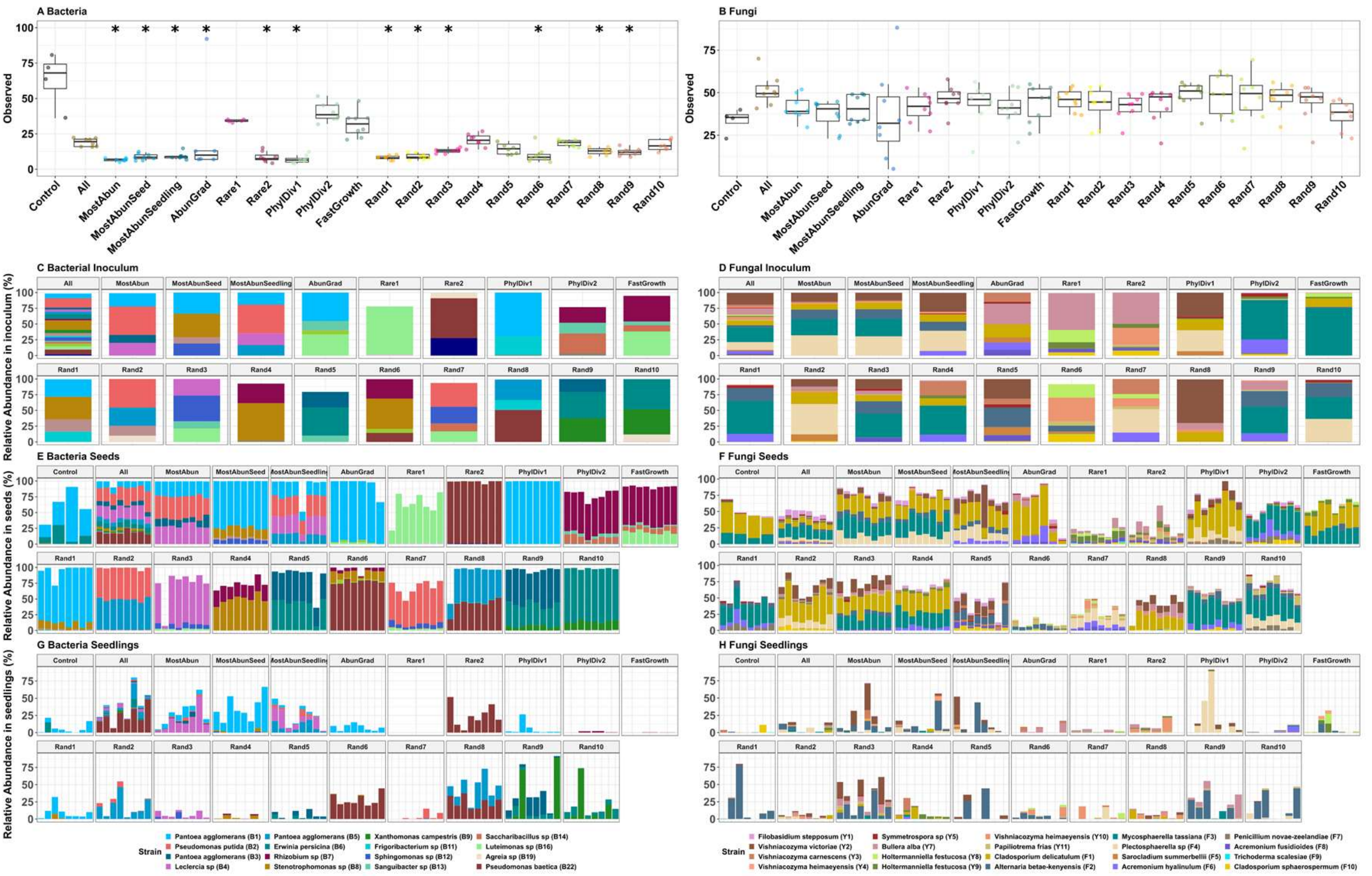


Supplementary Fig. 3: Assessment of microbial alpha diversity and taxonomic composition across seeds and seedlings. (A–B) Observed alpha richness (number of unique ASVs) for (A) bacterial and (B) fungal communities. For the non-inoculated control condition, each replicate point represents a pooled batch of 20 seeds, whereas all other points represent individual inoculated seeds. Asterisks denote significant differences relative to the control group (Pairwise comparisons using Dunn's many-to-one test with Benjamini-Hochberg correction, $p < 0.05$). (C–H) Taxonomic profiling of recovered inoculated strains across different experimental stages for bacterial (left panels: C, E, G) and fungal (right panels: D, F, H) communities. Panels represent relative abundances within the initial inoculum (C, D), the mature seed microbiota (E, F), and the emerged seedling microbiota (H, G).

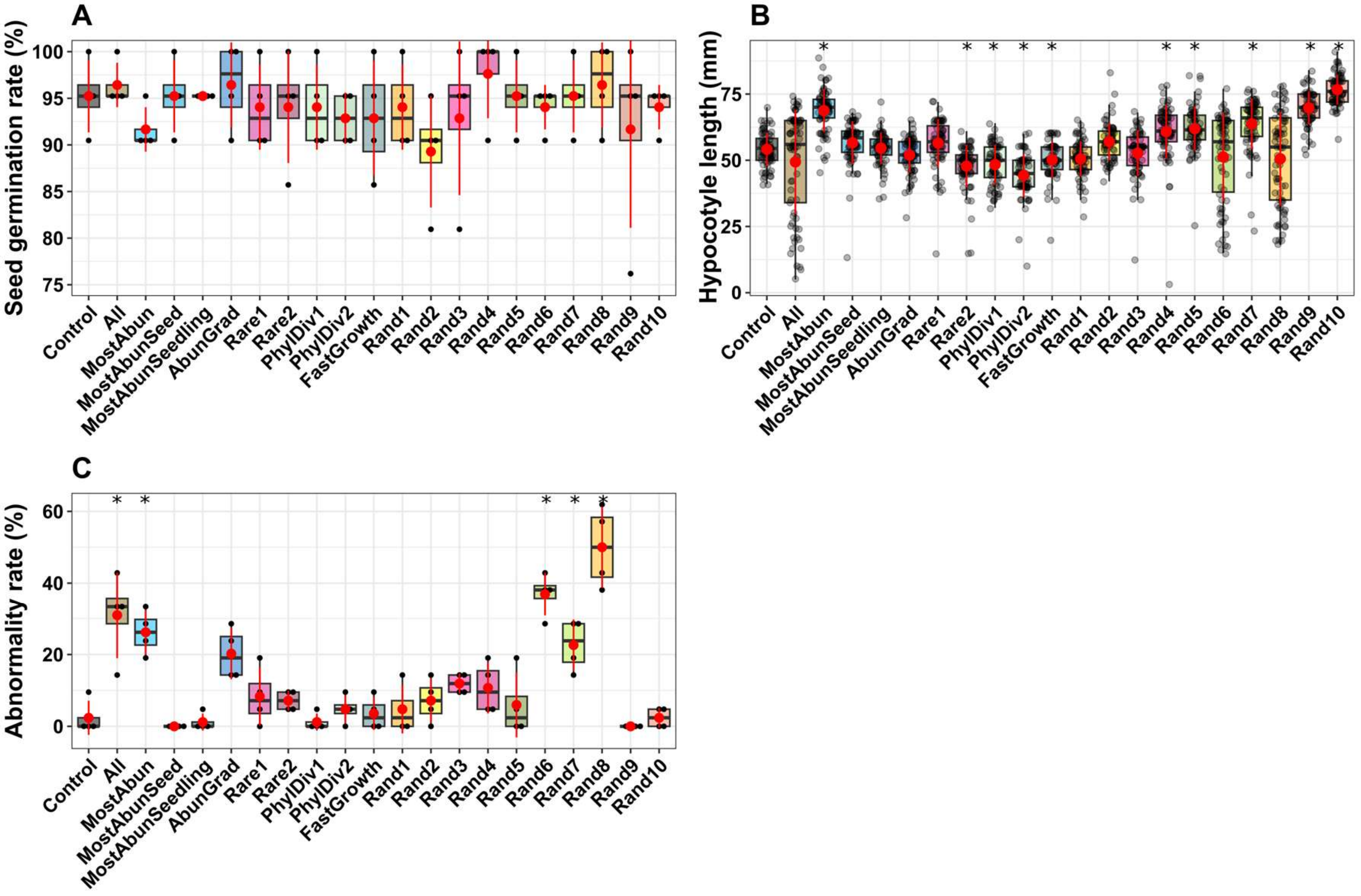


Supplementary Fig. 4: Phenotypic assessment of *Brassica napus* seedlings 15 days post-inoculation with various SynCom formulations. (A) Germination percentages, (B) hypocotyl length, and (C) proportion of morphologically normal seedlings. Asterisks (*) denote statistically significant differences relative to the non-inoculated control group (Pairwise comparisons using Dunn's many-to-one test with Benjamini-Hochberg correction, $p < 0.05$).

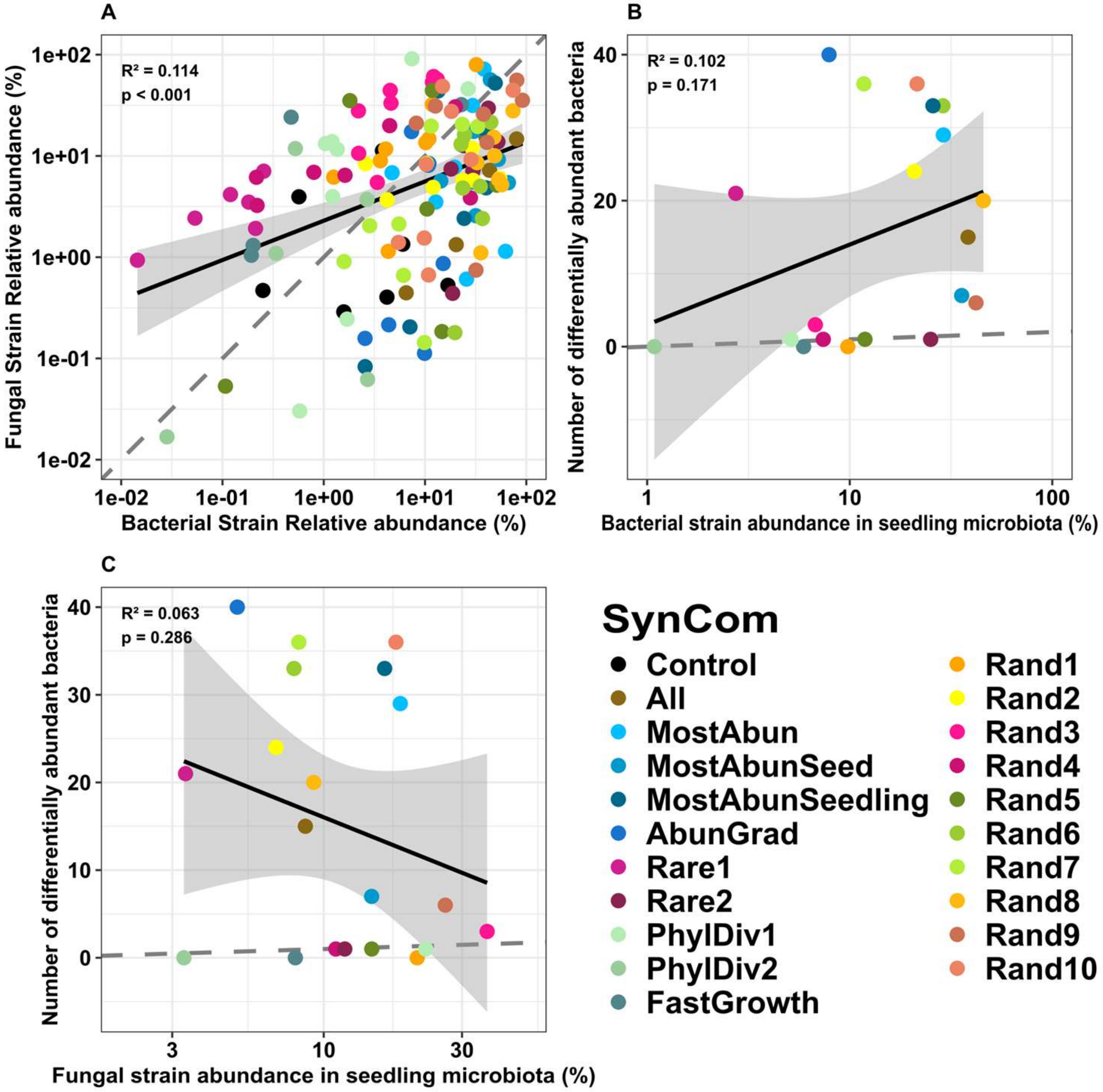


Supplementary Fig. 5: Correlation analyses between colonization dynamics and differential taxonomic abundance in seedlings. (A) Linear regression evaluating the relationship between the cumulative relative abundance of fungal and bacterial SynCom strains within individual seedlings. (B–C) Relationship between the number of differentially abundant host taxa and the cumulative abundance of introduced (B) bacterial or (C) fungal SynCom strains within the seedling matrix.

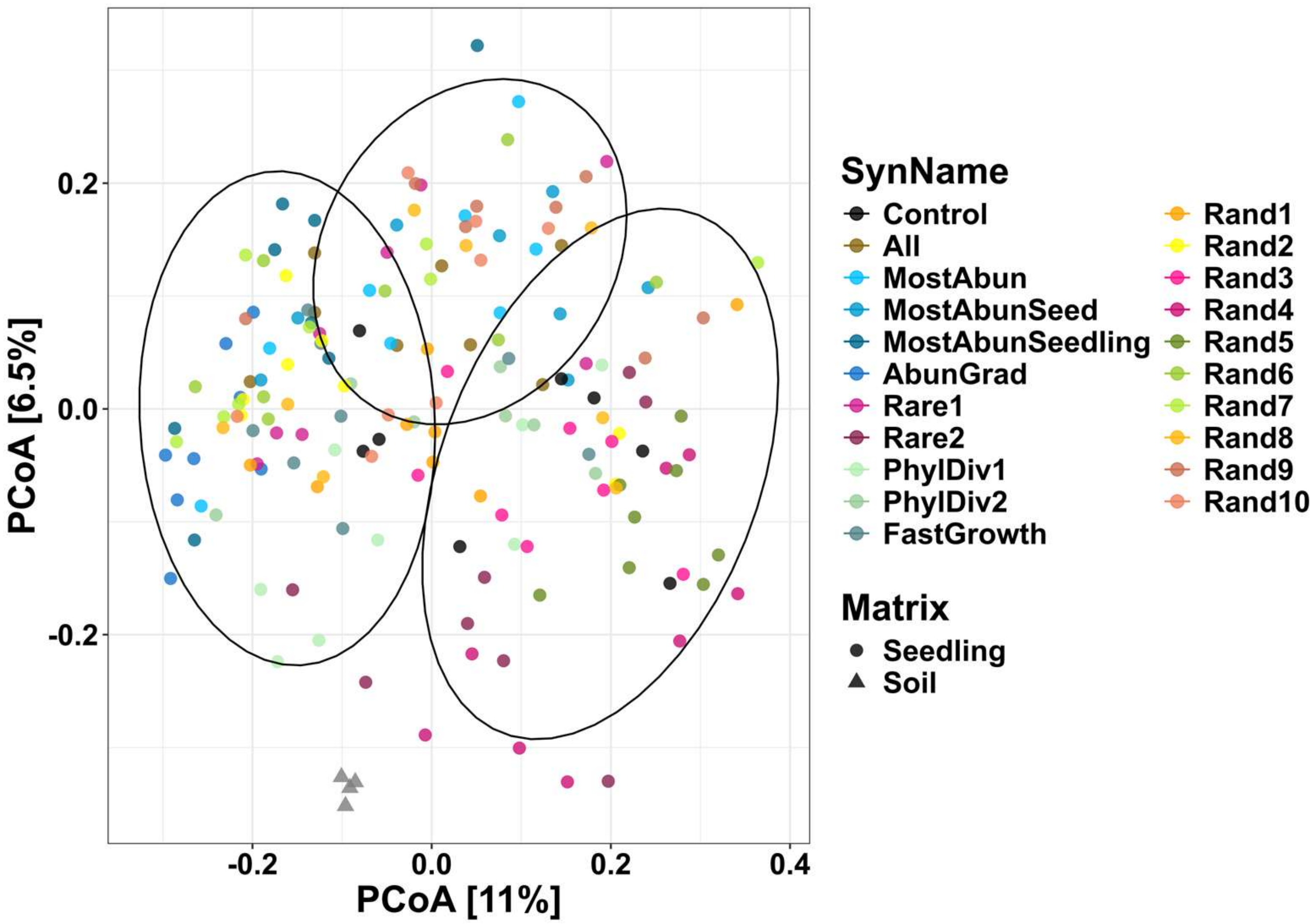


Supplementary Fig. 6: Principal Coordinate Analysis (PCoA) based on Bray-Curtis dissimilarity matrices (log(x+1) transformed) for fungal communities across bulk soil, non-inoculated control seedlings, and SynCom-inoculated seedlings. Cluster overlays are derived from k-means clustering partitioning using the NbClust package in R based on Euclidean distances.

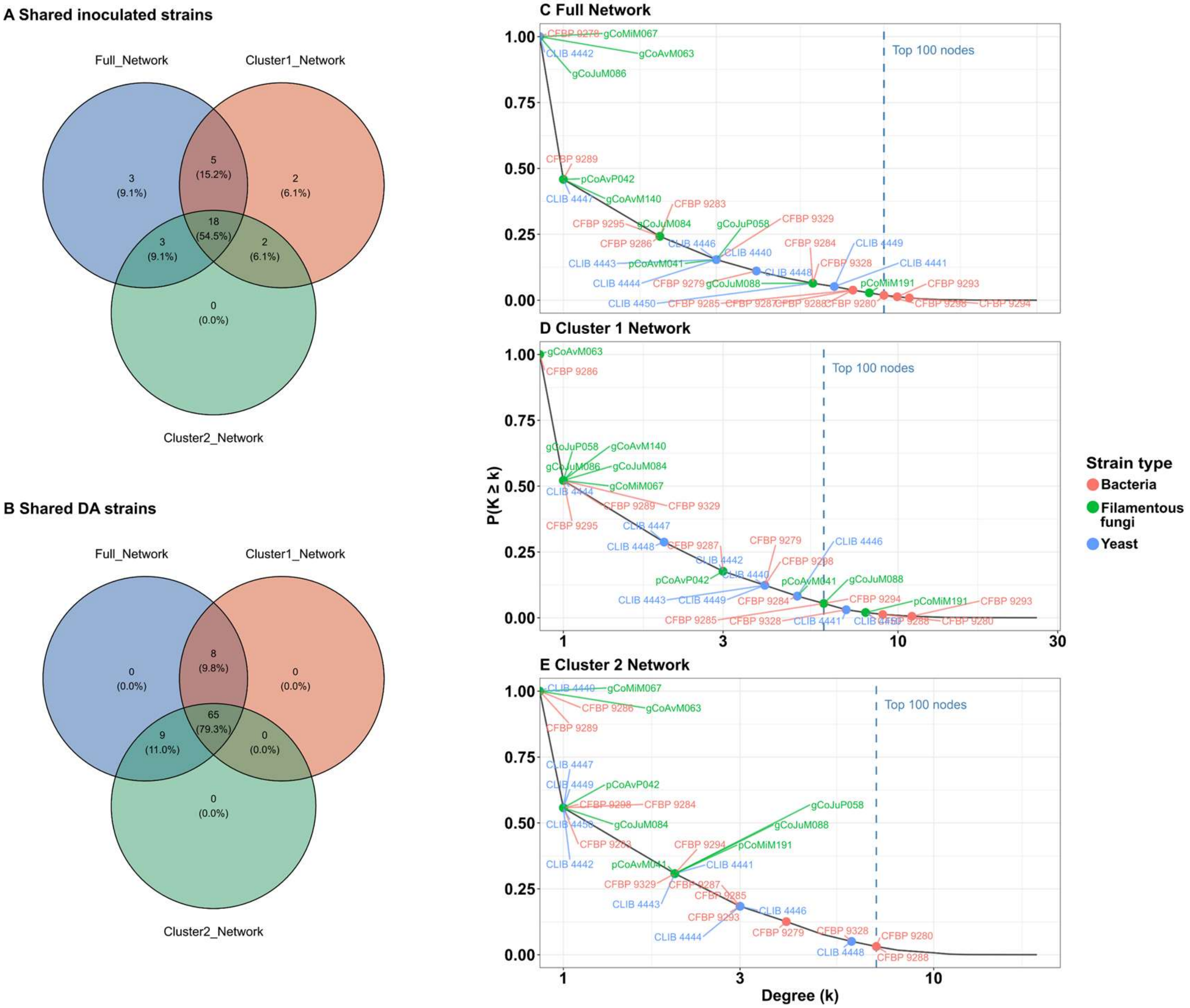


Supplementary Fig. 7: (A–B) Venn diagrams illustrating the overlapping and unique (A) introduced SynCom strains and (B) differentially abundant bacterial ASVs localized within the giant components of the Full, Cluster 1 and Cluster 2 networks. (C–E) Complementary cumulative distribution function (cCDF) curves modeling the topological degree distributions within the (C) full, (D) Cluster 1 and (E) Cluster 2 network models. Individual introduced SynCom strains are highlighted by taxonomic groups: bacteria (green points), filamentous fungi (red points), and yeasts (blue points). The vertical black dashed line marks the threshold designating the top 100 most highly connected nodes within each respective network architecture.
